# Reversible, Orally Bioavailable Small-Molecule Inhibitors Occupy the Substrate Tunnel of the Thioesterase Domain of Fatty Acid Synthase

**DOI:** 10.64898/2026.10.01.756096

**Authors:** Joseph Fromm, Alexander Aleshin, Lester J. Lambert, Laurent J. S. De Backer, Santhi Reddy Ganji, Margrith E. Mattmann, Allison S. Limpert, Palak Gosalia, Carina D. Wimer, Fabiana Izidro Layng, Maria Celeridad, Robert Ardecky, Peter Teriete, David A. Scott, Michael P. Hedrick, Jeffrey W. Smith, Douglas J. Sheffler, Nicholas D. P. Cosford

**Affiliations:** Center for Therapeutics Discovery, Cancer Metabolism and Microenvironment Program, NCI-Designated Cancer Center, Sanford Burnham Prebys Medical Discovery Institute, 10901 North Torrey Pines Road, La Jolla, California 92037, USA; Conrad Prebys Center for Chemical Genomics, Sanford Burnham Prebys Medical Discovery Institute, La Jolla, California, USA; Cancer Metabolism Core, Sanford Burnham Prebys Medical Discovery Institute, 10901 North Torrey Pines Road, La Jolla, California 92037, USA

**Keywords:** FASN, thioesterase, serine hydrolase, de novo lipogenesis, chemical probe, arylthiazole, X-Ray crystallography, MASH

## Abstract

Fatty acid synthase (FASN) is the sole enzyme capable of lipogenesis in cells and is upregulated in many cancers. Herein we report the discovery and development of **SBP-5917** (**78S**), a novel, reversible, small-molecule probe which targets the thioesterase (TE) domain of FASN. **SBP-5917** and other compounds in the series possess IC_50_ < 1 µM and < 5 µM for inhibiting recombinantly purified FASN-TE and cellular FASN respectively. Co-crystal structures of our inhibitors bound to the FASN-TE domain show that they are positioned in the substrate tunnel of the enzyme. Biochemical and biophysical analyses quantify the strength of compound binding and demonstrate their selectivity for FASN-TE. Preliminary pharmacokinetic studies indicate promising in vivo potency for many of the compounds in the series. **SBP-5917** is the first structurally validated, reversible inhibitor of FASN-TE with demonstrated cellular target engagement and can serve as a chemical probe to investigate FASN as a therapeutic target.

**ABSTRACT FIGURE:** 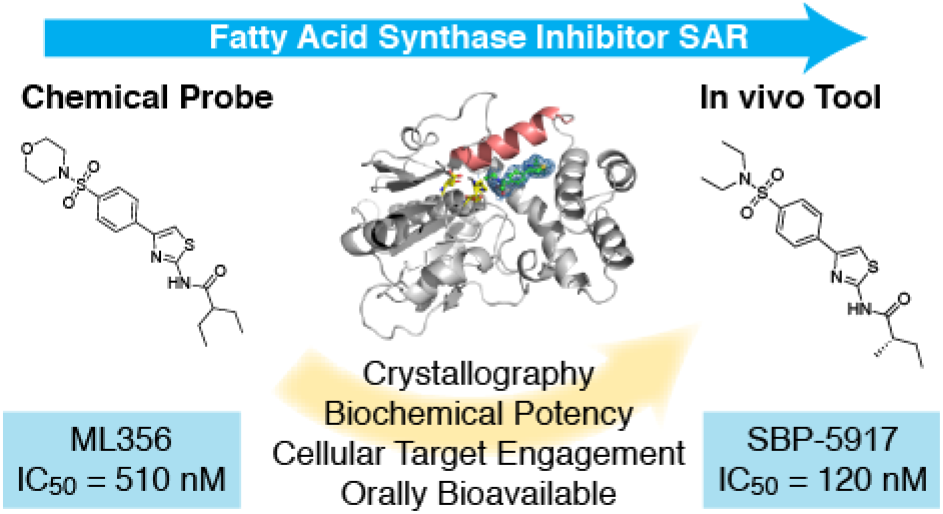

## INTRODUCTION

Fatty acid synthase (FASN) is the human enzyme responsible for the de novo synthesis of the 16-carbon fatty acid palmitic acid (palmitate) from dietary carbohydrates ^1, 2^. Mammalian FASN is a homodimeric polyprotein consisting of six enzymatic subunits: ketoacyl synthase (KS), malonyl/acyl-synthase (MAT), dehydratase (DH), enoyl reductase (ER), ketoacyl reductase (KR), a thioesterase (TE) domain, and an acyl-carrier protein (ACP) ^3–5^. The enzymatic domains of FASN convert acetyl- and malonyl-CoA through multiple cycles of NADPH-dependent sequential condensation into palmitate ^6^. Product release by a finishing hydrolysis “shunt” is the function of the TE domain ^7, 8^. This cleavage reaction between the completed palmitate and the phosphopantetheine of the ACP makes the FASN-TE domain the primary control point for size selection of the product fatty acid ^9^.

There is compelling evidence suggesting that FASN plays a mechanistic role in tumor progression ^10–18^. Increased expression of FASN is a hallmark of a number of major cancers and is linked to poor prognosis ^19–28^. Importantly, since dietary fat provides the nutrients needed for normal physiology, FASN is only of marginal importance in healthy adults ^29, 30^. As first described by Otto H. Warburg, the nutrient deprived core of solid tumors undergoes a metabolic “rewiring” to aerobic glycolysis that fuels high rates of tumor cell growth and proliferation ^31^. Proliferating tumor tissues commonly exhibit an increased rate of lipid metabolism ^28, 32^. This upregulated pool of fatty acids promotes increased membrane biosynthesis for rapid cell division ^3^. Moreover, FASN over-expression has mechanistic consequences that contribute to pro-growth stimuli from major signaling pathways ^15, 33–36^. Inhibition of FASN is therefore an attractive target for therapeutic intervention in the treatment of cancer (**SI Figure 1**).

Beyond cancer, FASN-driven de novo lipogenesis is markedly upregulated in metabolic dysfunction-associated steatohepatitis (MASH), where it contributes up to 26% of hepatic triglyceride accumulation compared to less than 5% in healthy liver, making FASN a compelling target in liver disease as well as oncology ^37^. The recent Phase 2b clinical success of denifanstat (TVB-2640), a FASN inhibitor targeting the ketoreductase (KR) domain, in reducing hepatic steatosis and fibrosis in MASH patients further validates FASN inhibition as a therapeutic strategy across disease contexts ^38^. The thioesterase domain inhibitors described herein, which act at a mechanistically distinct site from KR-domain inhibitors, may therefore have utility in FASN-dependent metabolic diseases.

Multiple groups have previously reported small molecules targeting various catalytic domains of FASN ^39^. For example, thiolactomycin, cerulenin C75, and compound 10v from Merck inhibit the KS domain ^40–42^, while TVB-2640, FT-113, GSK2194069, and JNJ-54302833 have been reported as inhibitors of the KR domain ^33, 43–45^. Currently TVB-2640 is undergoing a Phase 1 clinical study as a combinatorial therapy for metastatic prostate cancer ^46^. It also recently completed a Phase 3 clinical study investigating recurrent glioblastoma ^47^ and several Phase 2 clinical studies as a putative combinatorial cancer therapeutic in non-small lung cancer ^48^, breast cancer ^49^, and astrocytoma ^50^. Furthermore, inhibition of FASN expression has also been investigated in the clinic with the FDA approved proton pump inhibitor omeprazole ^51, 52^.

The 32 kDa FASN-TE is a serine hydrolase responsible for the cleavage of the product palmitate. The discovery and development of inhibitors that target the TE domain of FASN (FASN-TE) has attracted less attention relative to the other FASN domains (**SI Figure 2**). The best characterized FASN-TE inhibitor is Orlistat, an anti-obesity drug that inhibits gastric lipase and also inhibits FASN-TE in vitro ^53^. However, Orlistat is an irreversible substrate analogue that alkylates the active-site serine of FASN-TE with negligible systemic bioavailability, acting locally within the gastrointestinal tract, and this prevents its use for the systemic treatment of tumors ^54^. Another FASN-TE inhibitor described in the literature is the tool compound IPI-9119, which, like Orlistat, is also an irreversible alkylator of the active site Ser of FASN-TE ^55, 56^. Crystal structures of FASN-TE bound to several substrate-mimetic inhibitors have been previously reported ^8, 57, 58^. One of these crystal structures shows that Orlistat binds in the substrate binding tunnel that extends away from the active site ^57^.

Our laboratory previously discovered the first small molecules that reversibly inhibit the TE domain of FASN ^59^. Herein we describe the optimization of these molecules to generate potent, selective, reversible, small-molecule tool compounds that can potentially be utilized to probe the function of the FASN-TE domain and the role of FASN in cancer.

## RESULTS

### Small Molecules Arylthiazoles Potently Inhibit the Thioesterase Domain of FASN

To identify small molecules that inhibit FASN-TE, we performed a high-throughput screen (HTS) of ∼362,000 compounds from the Molecular Libraries Small Molecule Repository (MLSMR) compound collection as part of the National Institutes of Health’s Molecular Libraries Probe Production Centers Network (MLPCN) program ^60^. This HTS employed a biochemical assay utilizing recombinantly purified FASN-TE (2204-2511) and O-methyl fluorescein heptanoate (OMFH) as a fluorogenic substrate ^61^. From this screen, an arylthiazole derivative (**2**) was identified. After limited structure-activity relationship (SAR) studies, compound **3** (**ML356**) (**Scheme 1A**) was identified as a useful first generation FASN-TE inhibitor probe compound ^59^.

**Scheme 1:**
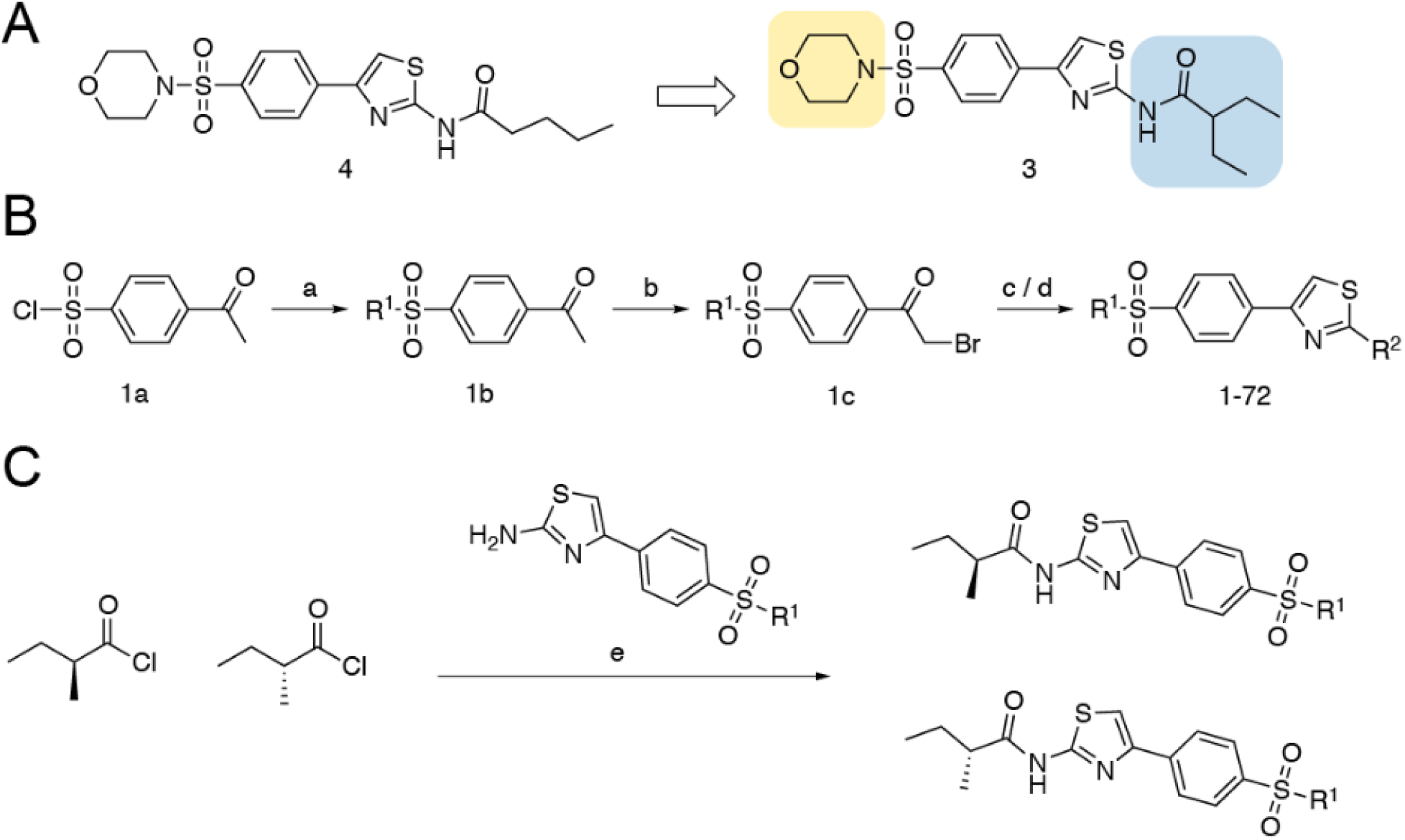
Chemical synthesis of arylthiazole inhibitors of the TE domain of FASN. (A) Initial SAR optimization of Compound **2** from the HTS yielded Compound **6**. The sulfonamide (yellow: R^1^) and lipophilic (blue: R^2^) groups were the focus of further development. (B) *Reagents and Reaction conditions:* (a) NHR^1^, Et_3_N, CH_2_Cl_2_. (b) Br_2_, MeOH:H_2_O. (c) Thiourea, Na_2_CO_3_, THF. (d) R^2^COCl, pyridine, CH_2_Cl_2_. (e) DIPEA, CH_2_Cl_2,_ 60°C. A detailed synthesis for **50** is provided (**SI Scheme 1**). (C) Chemical synthesis of stereopure arylthiazole FASN-TE inhibitors. *Reagents and Reaction conditions:* (*S*)-2-methylbutanoyl chloride or (*R*)-2-methylbutanoyl chloride are reacted with the 2-amine thiazole (100-107) (**SI Scheme 2**) in Et_3_N, CH_2_Cl_2_ or pyridine, CH_2_Cl_2_.

Analysis of the structure of **ML356** (**3**) and similar small molecule hits from the HTS campaign suggested two synthetic strategies for additional SAR studies (**Scheme 1A**). Consequently, our optimization focused on (a) varying the acyl group on the primary amine, and (b) varying the amino substituent on the sulfonamide moiety. For the acylamino moiety, we synthesized analogues with an acyl chain of varied length and/or branching as well as the introduction of cycloalkyl, pyridyl, or fluorophenyl groups. For the sulfonamide moiety, we investigated the effects of varying the size and structure of the cyclic amine and replacement with acyclic amine substituents. Two synthetic routes were employed for the preparation of analogues (**Scheme 1B**). Thus, 4-acetylbenzenesulfonyl chloride (**1a**) is reacted with a secondary amine to provide the sulfonamide derivative (**1b**). The resulting compound is then brominated under standard conditions to afford a bromoketone (**1c**), which is readily converted to the corresponding arylthiazole derivative by reaction with thiourea. Finally, acylation of the arylthiazole with an appropriate acyl chloride provides the target amide derivatives (**1**).

Using this synthetic route, more than 50 analogues were synthesized and tested using the biochemical OMFH assay. Characterization of the compounds from the initial SAR are provided (**Table 1**), wherein we varied the sulfonamide substituent and observed some patterns for improved potency. Replacing the R^1^ morpholine moiety with other six-membered ring derivatives, as exemplified by **22, 24**, **28, 29**, **30,** and **63**, had little effect on potency relative to the morpholine analogues (**4, 3, 8,** and **62** respectively). However, sulfonamides with the substituent (R^1^) diethylamine (**47, 48, 54,** and **67**) or the five-membered pyrrolidine ring (**33, 35**, and **41**) generally exhibited improved potency. Probing the effect of acyl or aryl substitution on the thiazoleamine (R^2^) was also studied. Compounds with a smaller (R^2^) acyl group (**2**, **46**) or distal branching (**6**, **23**) showed reduced affinity relative to the 2-ethyl butanamide present in **ML356** (**3**) and other comparable analogues. Longer acyl groups (**29, 52**) offered no improvement in potency. Replacement of R^2^ with a cycloalkyl group, as in **59** and **61**, or a heteroaromatic moiety (**45**) resulted in less potent compounds. On the other hand, shortening the alkyl branching at the R^2^ α-carbon, as in **7**, **38**, and **50**, led to compounds with similar or improved potencies and affinities compared with the 2-ethyl butanamide.

**Table 1:**
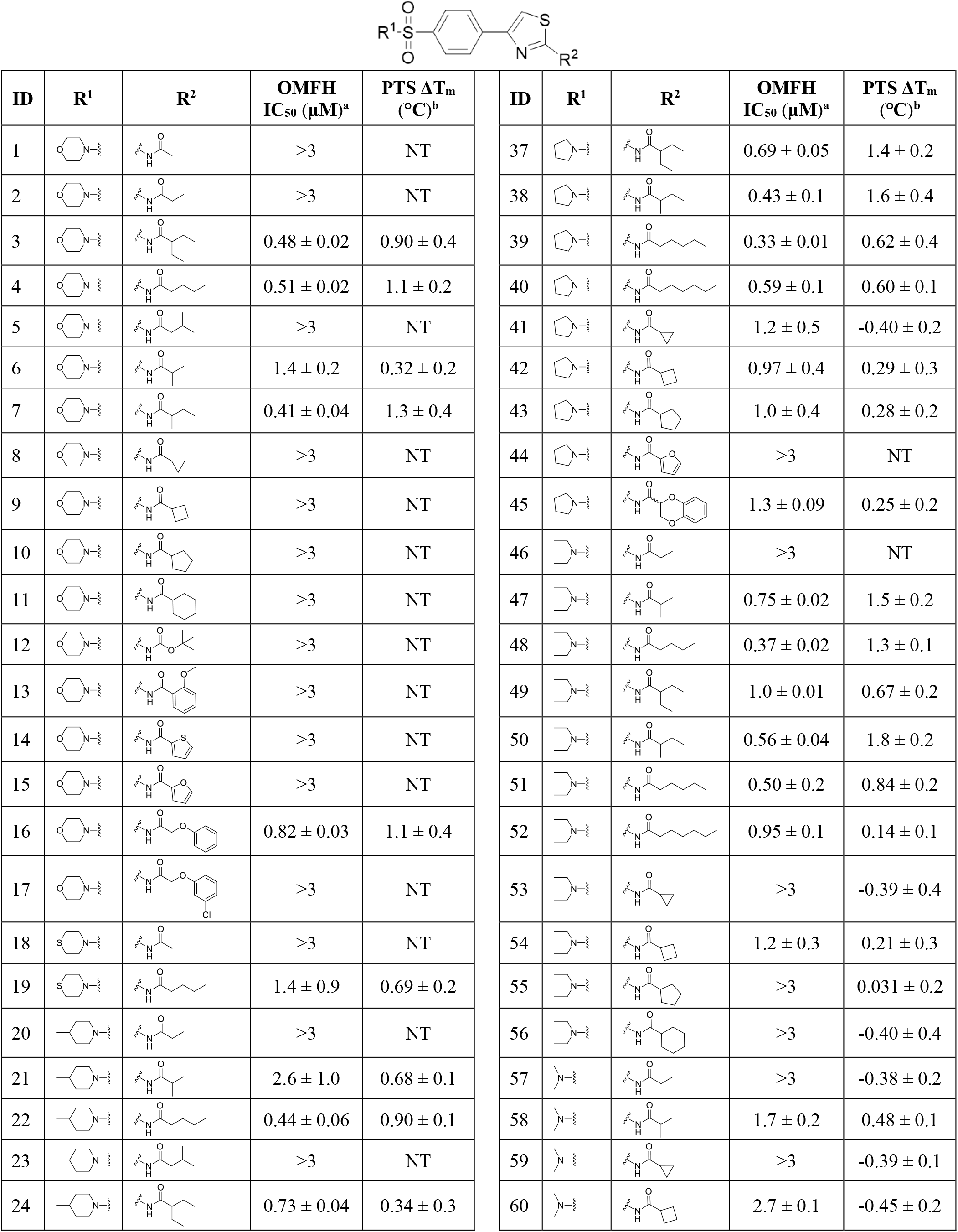
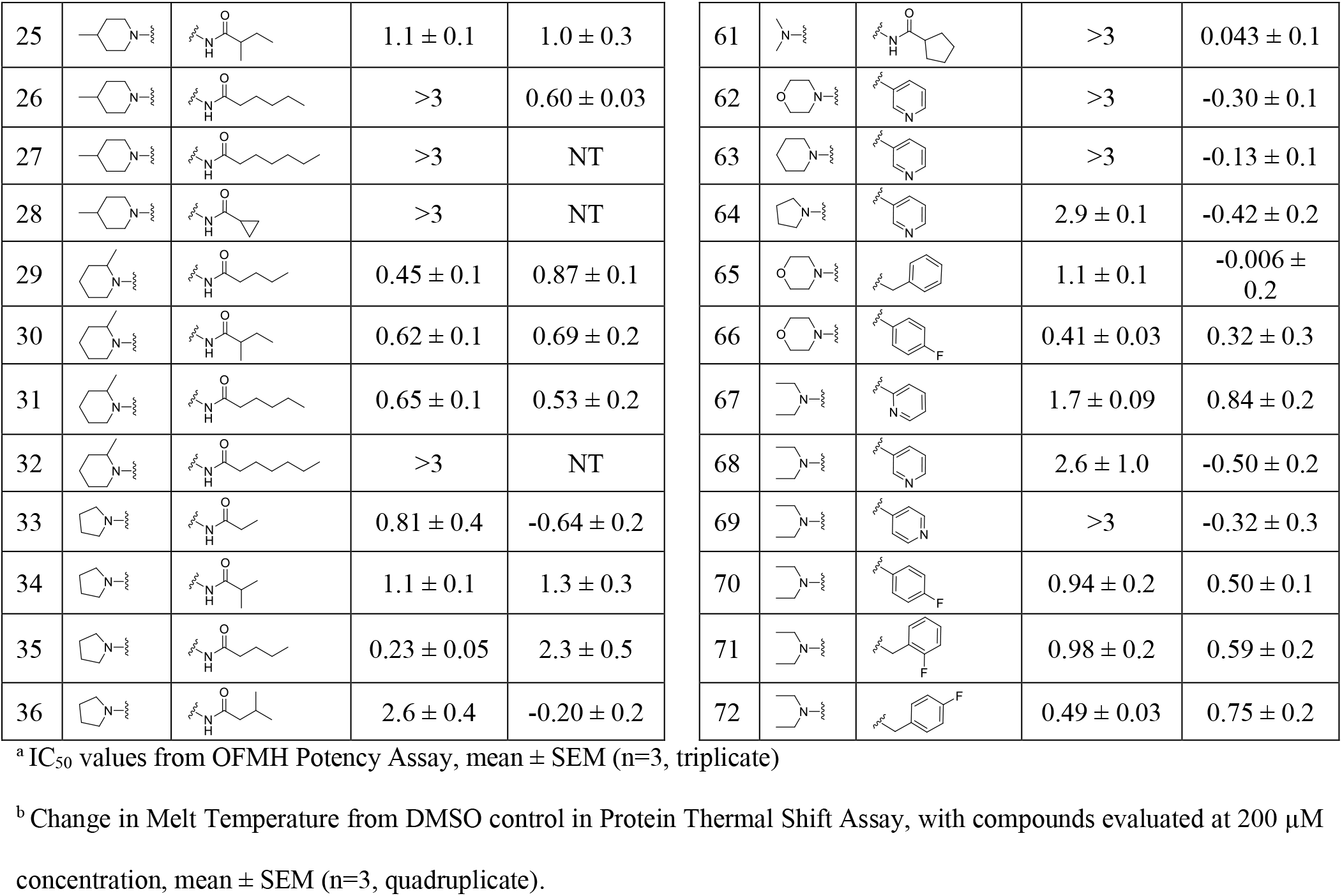
Arylthiazole inhibitors of FASN-TE with biochemical potency and protein stabilization efficacy.

| ID | R<sup>1</sup> | R<sup>2</sup> | OMFH<br>IC<sub>50</sub> (μM)<sup>a</sup> | PTS ΔT<sub>m</sub><br>(°C)<sup>b</sup> | ID | R<sup>1</sup> | R<sup>2</sup> | OMFH<br>IC<sub>50</sub> (μM)<sup>a</sup> | PTS ΔT<sub>m</sub><br>(°C)<sup>b</sup> |
| --- | --- | --- | --- | --- | --- | --- | --- | --- | --- |
| 1 |  |  | >3 | NT | 37 |  |  | 0.69 ± 0.05 | 1.4 ± 0.2 |
| 2 |  |  | >3 | NT | 38 |  |  | 0.43 ± 0.1 | 1.6 ± 0.4 |
| 3 |  |  | 0.48 ± 0.02 | 0.90 ± 0.4 | 39 |  |  | 0.33 ± 0.01 | 0.62 ± 0.4 |
| 4 |  |  | 0.51 ± 0.02 | 1.1 ± 0.2 | 40 |  |  | 0.59 ± 0.1 | 0.60 ± 0.1 |
| 5 |  |  | >3 | NT | 41 |  |  | 1.2 ± 0.5 | -0.40 ± 0.2 |
| 6 |  |  | 1.4 ± 0.2 | 0.32 ± 0.2 | 42 |  |  | 0.97 ± 0.4 | 0.29 ± 0.3 |
| 7 |  |  | 0.41 ± 0.04 | 1.3 ± 0.4 | 43 |  |  | 1.0 ± 0.4 | 0.28 ± 0.2 |
| 8 |  |  | >3 | NT | 44 |  |  | >3 | NT |
| 9 |  |  | >3 | NT | 45 |  |  | 1.3 ± 0.09 | 0.25 ± 0.2 |
| 10 |  |  | >3 | NT | 46 |  |  | >3 | NT |
| 11 |  |  | >3 | NT | 47 |  |  | 0.75 ± 0.02 | 1.5 ± 0.2 |
| 12 |  |  | >3 | NT | 48 |  |  | 0.37 ± 0.02 | 1.3 ± 0.1 |
| 13 |  |  | >3 | NT | 49 |  |  | 1.0 ± 0.01 | 0.67 ± 0.2 |
| 14 |  |  | >3 | NT | 50 |  |  | 0.56 ± 0.04 | 1.8 ± 0.2 |
| 15 |  |  | >3 | NT | 51 |  |  | 0.50 ± 0.2 | 0.84 ± 0.2 |
| 16 |  |  | 0.82 ± 0.03 | 1.1 ± 0.4 | 52 |  |  | 0.95 ± 0.1 | 0.14 ± 0.1 |
| 17 |  |  | >3 | NT | 53 |  |  | >3 | -0.39 ± 0.4 |
| 18 |  |  | >3 | NT | 54 |  |  | 1.2 ± 0.3 | 0.21 ± 0.3 |
| 19 |  |  | 1.4 ± 0.9 | 0.69 ± 0.2 | 55 |  |  | >3 | 0.031 ± 0.2 |
| 20 |  |  | >3 | NT | 56 |  |  | >3 | -0.40 ± 0.4 |
| 21 |  |  | 2.6 ± 1.0 | 0.68 ± 0.1 | 57 |  |  | >3 | -0.38 ± 0.2 |
| 22 |  |  | 0.44 ± 0.06 | 0.90 ± 0.1 | 58 |  |  | 1.7 ± 0.2 | 0.48 ± 0.1 |
| 23 |  |  | >3 | NT | 59 |  |  | >3 | -0.39 ± 0.1 |
| 24 |  |  | 0.73 ± 0.04 | 0.34 ± 0.3 | 60 |  |  | 2.7 ± 0.1 | -0.45 ± 0.2 |

|  |  |  |  |  |
| --- | --- | --- | --- | --- |
| 25 |  |  | 1.1 ± 0.1 | 1.0 ± 0.3 |
| 26 |  |  | >3 | 0.60 ± 0.03 |
| 27 |  |  | >3 | NT |
| 28 |  |  | >3 | NT |
| 29 |  |  | 0.45 ± 0.1 | 0.87 ± 0.1 |
| 30 |  |  | 0.62 ± 0.1 | 0.69 ± 0.2 |
| 31 |  |  | 0.65 ± 0.1 | 0.53 ± 0.2 |
| 32 |  |  | >3 | NT |
| 33 |  |  | 0.81 ± 0.4 | -0.64 ± 0.2 |
| 34 |  |  | 1.1 ± 0.1 | 1.3 ± 0.3 |
| 35 |  |  | 0.23 ± 0.05 | 2.3 ± 0.5 |
| 36 |  |  | 2.6 ± 0.4 | -0.20 ± 0.2 |
| 61 |  |  | >3 | 0.043 ± 0.1 |
| 62 |  |  | >3 | -0.30 ± 0.1 |
| 63 |  |  | >3 | -0.13 ± 0.1 |
| 64 |  |  | 2.9 ± 0.1 | -0.42 ± 0.2 |
| 65 |  |  | 1.1 ± 0.1 | -0.006 ± 0.2 |
| 66 |  |  | 0.41 ± 0.03 | 0.32 ± 0.3 |
| 67 |  |  | 1.7 ± 0.09 | 0.84 ± 0.2 |
| 68 |  |  | 2.6 ± 1.0 | -0.50 ± 0.2 |
| 69 |  |  | >3 | -0.32 ± 0.3 |
| 70 |  |  | 0.94 ± 0.2 | 0.50 ± 0.1 |
| 71 |  |  | 0.98 ± 0.2 | 0.59 ± 0.2 |
| 72 |  |  | 0.49 ± 0.03 | 0.75 ± 0.2 |

<sup>a</sup> IC<sub>50</sub> values from OFMH Potency Assay, mean ± SEM (n=3, triplicate)
<sup>b</sup> Change in Melt Temperature from DMSO control in Protein Thermal Shift Assay, with compounds evaluated at 200 μM concentration, mean ± SEM (n=3, quadruplicate).

Derivatizing the thiazoleamine as either primary or secondary arylthiazole derivatives was well tolerated. The greatest biochemical potency was achieved by the 4-fluorobenzyl (**72**) and the 4-fluorophenyl (**66**), which were both comparable to **ML356** (**3**). However, other fluorophenyl derivatives (**70** and **71**) exhibited slightly reduced potency compared with their counterparts. Pyridine groups at the R^2^ arylthiazole position (**61**, **62**, and **64**) had significantly reduced potency and affinity relative to **ML356** (**3**).

As a complementary approach to benchmark this series of FASN-TE inhibitors, we developed and employed a differential scanning fluorimetry/protein thermal shift (PTS) assay using recombinant FASN-TE domain protein to probe the ability of the arylthiazoles to stabilize the FASN-TE domain, with data shown in **Table 1**. Evaluating those compounds which possessed an IC_50_ < 3.0 µM in the biochemical OMFH assay, along with a few select additional compounds, we observed 12 compounds that were able to stabilize the FASN-TE with a ΔTm > 1.0°C when tested at a concentration of 200 µM. These included several potent branched alkyl chain compounds **7**, **38** and **50**. Branched chain pyrrolidines **37**, **38** and straight chain diethylamine compounds (**48**) were among those observed to afford increased stability to the melt curve of FASN-TE. Although the modest stabilizing ΔTm PTS values of these compounds are to be interpreted with caution, we observed a consistent relationship between biochemical potency and FASN-TE stabilization (**SI Figure 3**). Notable is that the lipophilic dye sypro orange did not exert a significant effect on protein stability and was not an inhibitor in the OMFH assay. Furthermore, biochemical and PTS potency did not correlate with logP discounting the possibility of nonspecific assay interference. These combined biochemical and PTS results informed subsequent SAR.

### The Structural Basis for Arylthiazole Binding to the FASN-TE Domain

To improve our understanding of the structural basis for binding to the TE domain of FASN by our arylthiazole inhibitors, we initiated crystallization trials using recombinantly expressed TE domain. For reference we solved the structure of the apo form of FASN-TE (2204-2511) to 2.6 Å by molecular replacement using PDBID 3TJM as the search model (PDBID 9ZLX). Crystalizing as space group P 2_1_2_1_2_1_ with one molecule per asymmetric unit, a different crystal form than that reported previously, the structure of the apo-enzyme conforms closely to that previously described ^8^. Most importantly and consistent with previous findings, the electron density for the segment containing gatekeeper residues (2341-2358) in the apo-TE domain was not detected ^8^. The gatekeeper helix forms the outer shell to an amphipathic tunnel which harbors the growing acyl chain and flexibility is functionally intrinsic to the “scan and release” activity of the TE domain of FASN ^8^. Therefore, small molecule inhibitors that bind to this segment of the TE domain of FASN must have the capacity to conform to, and potentially stabilize, a malleable and dynamic binding pocket.

From a representative panel of our arylthiazole FASN-TE inhibitors, we obtained each of the co-crystal structures of reversible inhibitors **71** and **66** bound to the TE domain of FASN (**Figure 1A** and **1B**). The bound structures were solved by molecular replacement to 2.0 Å and 2.8 Å respectively. The overall structure of the bound conformations of the TE domain of FASN is unperturbed from the apo-state with a rmsd of 0.35 (7MHD) and 0.56 (7MHE) for main chain α-carbons. The ligands **71** and **66** were fit based on an extended tube of electron density enveloped by the ligand tunnel. As the terminal morpholine group projects from the binding pocket, the electron density for **66** is incomplete so the chair conformation is modeled. Furthermore, the gatekeeper helix residues (2341-2358) in the bound state structure are not fully resolved. However, there are fragments of electron density that allow it to be positioned enclosing the bound ligands.

**Figure 1:**
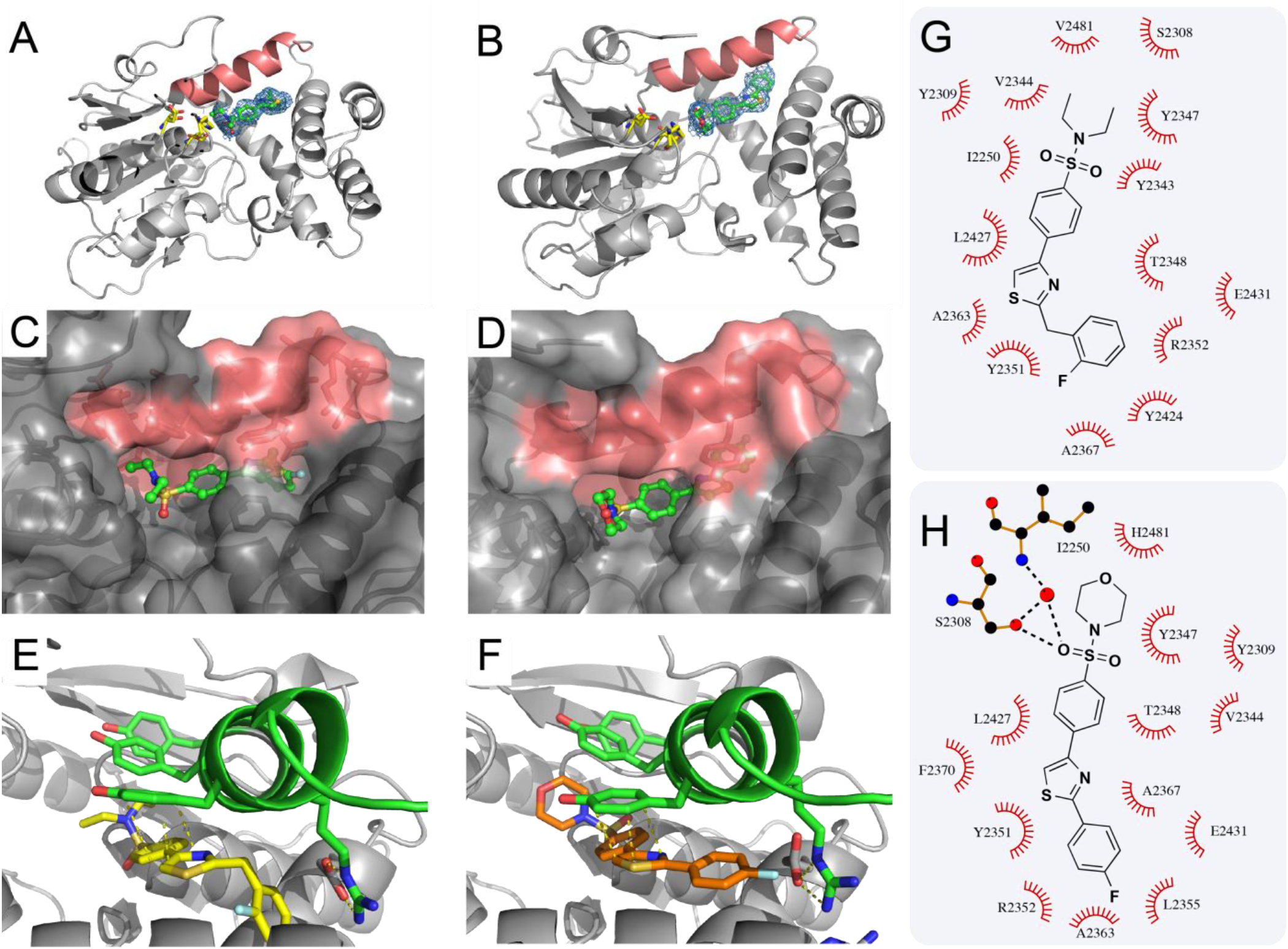
Arylthiazole reversible inhibitors bind in the ligand tunnel of FASN-TE. X-ray co-crystal structures of the TE domain of FASN bound in complex with reversible inhibitors **71** (PDBID 7MHD) and **66** (PDBID 7MHE) (A) Overall structure of the TE domain of FASN solved to 2.0 Å bound to **71** with the secondary structure of the α/β hydrolase subdomain indicated in grey. The catalytic triad residues are indicated as yellow sticks. The **71** inhibitor is represented as balls and sticks with carbon atoms colored green. An Fo - Fc omit map (blue) indicating the electron density contoured at 3σ within 3 Å of the bound ligand is shown. The position of the gatekeeper helix (2341-2358) is indicated in red. (B) The TE domain structure solved to 2.8 Å bound to **66.** A similar electron density for the ligand was fitted as shown in A. (C) Surface diagram representation of the bound conformation of **71**. Gatekeeper resides are colored red and the **71** inhibitor is represented as balls and sticks with carbon atoms colored green. (D) Surface diagram representation of the bound **66**. (E) Alignment of Y2351 with arylthiazole **71**. (F) Alignment of Y2351 with arylthiazole **66**. (G) Schematic showing the < 4.0 Å interactions by **71** to specific residues of the TE domain indicated as non-polar red arcs and/or polar interactions to indicated atoms. This diagram was made using LIGPLOT. (H) Interactions to TE domain by **66**.

The FASN-TE inhibitors **71** and **66** bind in the fatty acid acyl chain binding tunnel occupying dimensions that are like the lipid chain of palmitate. The overall orientation of these inhibitors is with the morpholino- or diethylamine-sulfonamide positioned adjacent to the catalytic triad, the phenyl arylthiazole extended into the substrate tunnel, and the fluorophenyl ring probing deeply towards flanking residues E2431 and A2363 in a deformable inner pocket (**Figure 1C** and **1D**). The thiazole rings of both **71** and **66** align with the aromatic ring of Y2351, separated by 3.5 Å (**Figure 1E** and **1F**) using a binding mode consistent with other thiazole-aromatic π-stacking interactions ^62^. The hydrophobic tunnel, spanning the two subdomains of the TE domain, extends further to terminate into a cluster of charged residues R2352, K2354, E2360 and E2362 that creates what is effectively a charged barrier for a growing alkyl chain.

Inhibitors **71** and **66** engage several hydrophobic contacts in the binding tunnel with residues in both the gatekeeper helix and the groove formed by helices (2412-2432) and (2360-2373) (**Figure 1G** and **1H**). These contacts are comparable for the phenyl linked arylthiazole core scaffold for both **71** and **66**. Some differences in the embedded structures of the two ligands are evident. The 2-fluorobenzyl substituent of **71** kinks towards favorable contacts with a pocket produced by the peptide chain of A2367, R2352 and Y2424. These contact residues are different for the 4-fluorophenyl ring of **66** which forms a nonpolar inner surface contact with A2363 and L2355. That the plastic dimensions of the binding tunnel can accommodate the conformation of both ligands is consistent with the versatile lipid processing role of the TE domain of FASN.

The invariant residues of the side chains of the catalytic triad (S2308, H2481 and D2338) in both the bound **71** and **66** structures are proximal to the sulfonamide for these inhibitors. However, no direct polar contact is made to the catalytic S2308. Rather a network of ordered water molecules is detected for **66** in the bound state, and this ultimately serves as the basis for polar interaction with the activity determining residues. The morpholino- group of **66** bends out of the catalytic site but the lack of electron density does not allow the chair conformation to be established.

### Comparison of the Bound Conformation of Reversible Inhibitor 66 to Covalent Inhibitors Orlistat and Methyl γ-Linolenylphosphonate

The enclosed binding tunnel of the TE domain of FASN can accommodate several fatty acyl chains, with the (C16:0) palmitoyl determined to be an optimal fit ^58^. The reversible inhibitors **71** and **66** are of a similar length to this substrate. When the TE domain backbones of the covalent adduct structures for Orlistat (PDBID 2PX6-A) or methyl γ-linolenylphosphonate (MGLP) (PDBID 3TJM) are superimposed on that of **66** (PDBID 7MHE), differences in the conformational space adopted by the bound ligands are instructive (**SI Figure 4**) ^57, 58^. The first eight carbons of the Orlistat and MGLP chain occupy a groove pocket that the more planar arylthiazole core scaffold does not enter. Deeper into the ligand tunnel, flexibility in the relative position of the Orlistat and MGLP alkyl chains is evident. Compound **66** however adopts a rigid pose with striking differences in conformation proximal to the catalytic residues. Orlistat extends its second hexanoyl tail into a second groove adjacent to the catalytic site that neither **66** nor MGLP access. Similarly, the covalent conformation of the peptidyl moiety of Orlistat significantly differs from **66** and MGLP. The position of the morpholine moiety in **66** resembles the methylphosphonate head group of MGLP but pivots away from the catalytic triad rather than the covalently bound MGLP. Overall, the bound conformations of **71** and **66** are unlike either Orlistat or MGLP. Comparison of the multiple structural pharmacophores could permit rational design of inhibitor molecules that form increased contacts to this pliant and adaptive binding site.

### Effects of Arylthiazoles on the TE Domain of FASN Structure Probed by HSQC

To confirm a common binding mode between the arylthiazole series and the solved arylthiazole structures, we employed protein NMR spectroscopy to probe effects of ligand binding on the TE domain structure. Accordingly, uniformly ^15^N-labeled FASN-TE (2204-2518) was prepared, and an HSQC spectrum of the domain was obtained (**SI Figure 5A**). When either **50** or **66** are titrated into the ^15^N-labeled TE domain a small set of discrete crosspeaks were perturbed by the compounds (**SI Figure 5B-C**). This finding is consistent with the crystal structure results, which suggests that ligand binding produces minimal backbone changes between the bound and unbound forms. That the same subset of peaks is shifted suggests a shared binding mode for both compounds and is consistent with a common binding mode for related arylthiazole analogues. New crosspeaks that could possibly be assigned to gatekeeper helix tyrosine residues (based on the reported assignment ^63^) emerge in the bound state for **50** and **66** however these cannot be unambiguously assigned by inspection. The TE domain of FASN has previously been shown to be a challenging target for NMR studies as spectral overlap and relaxation dynamics frustrate full backbone assignment ^63^.

Overall, our conclusions for the observed bound state structures, the PTS stabilization data and the NMR studies are as follows. The close correspondence between the apo- and bound crystal forms of the FASN-TE domain is indicative of a well-folded domain. The PTS thermal melt data show that unfolding occurs at a discrete temperature transition in the typical range for < 100 kDa proteins which supports the crystallographic observations ^64^. Furthermore, the NMR HSQC data do not show obvious signs of multiple states in the form of peak doubling and/or broad line shape. There is, however, a rigidifying of the gatekeeper helix (2341-2358) in the bound state crystal structure. We conclude that the apo- gatekeeper helix (2341-2358) is dynamic and disordered to allow access for fatty acid chains. Upon binding our arylthiazole inhibitors, we observe the emergence of only a subset of distinct NMR crosspeaks, which is supportive of this rigidifying model. As the ordering that occurs upon ligand binding is confined to the binding tunnel, we also observe only modest thermal shift ΔTm stabilization consistent with low affinity ligand binding and the intrinsic limits that the binding tunnel can impart onto the overall fold ^64^. Importantly the stabilizing interaction of our most potent arylthiazoles, combined with biochemical potency, is a useful metric to characterize ligands on the TE domain. We utilized these properties to evaluate close analogues in the series.

### Synthesis of Stereopure Arylthiazole FASN-TE Inhibitors

Informed by the initial SAR and crystal structures, we synthesized a focused panel of stereopure arylthiazoles where we fixed the N-acyl 2-methyl butanamide at the R^2^ position and explored the effect of systemic substitutions to the sulfonamide substituent (R^1^) (**Scheme 1C**). To benchmark the effect of these changes we employed the OMFH and PTS assays (**Table 2**). The *S-*stereoisomer of 2-methyl butanamide with an R^1^ diethylamine group, **78S**, had a 4.7-fold improvement in potency in the OMFH assay (IC_50_ 0.12 µM) relative to the racemate, **50**. This increased potency correlates with the increased thermal stability ΔT_m_ of 2.9°C from 1.8°C. In contrast, the binding of the *R*-stereoisomer, **78R**, was less favored in both the OMFH enzymatic assay (IC_50_ of 1.2 µM) and PTS protein stability assay (ΔT_m_ of 1.1°C) than the racemate, **50**. An *S*-stereoisomer with a morpholine substituent at the R^1^ position, **73S,** had similar activity relative to the diethylamine, with an OMFH IC_50_ of 0.13 µM and ΔT_m_ of 2.5°C). This compound was also more potent (3.2-fold increase) and produced a stronger thermal shift than its racemate counterpart **7**, a trend which held true for all S-stereoisomer compounds. We explored the introduction of a piperidine (**80S**) (IC_50_ = 0.37 µM) as either 2-methyl (**76S**) or 4-methyl (**75S**) analogues; these shared similar potencies (0.44 and 0.29 µM respectively). A pyrrolidine, **77S**, retained potency as did the dimethyl amine, **79S**, consistent with the FASN-TE crystal structure pose indicating that the sulfonamide can expand into the solvent. The bulkier thiomorpholine moiety, as in **74S**, is not well tolerated at this position, with a 14-fold lower potency (IC_50_ 1.7 µM) than **78S** and displayed the weakest, though still substantial, thermal shift (ΔT_m_ = 1.5°C) of the *S-*enantiomer 2-methyl butanamides.

**Table 2:**
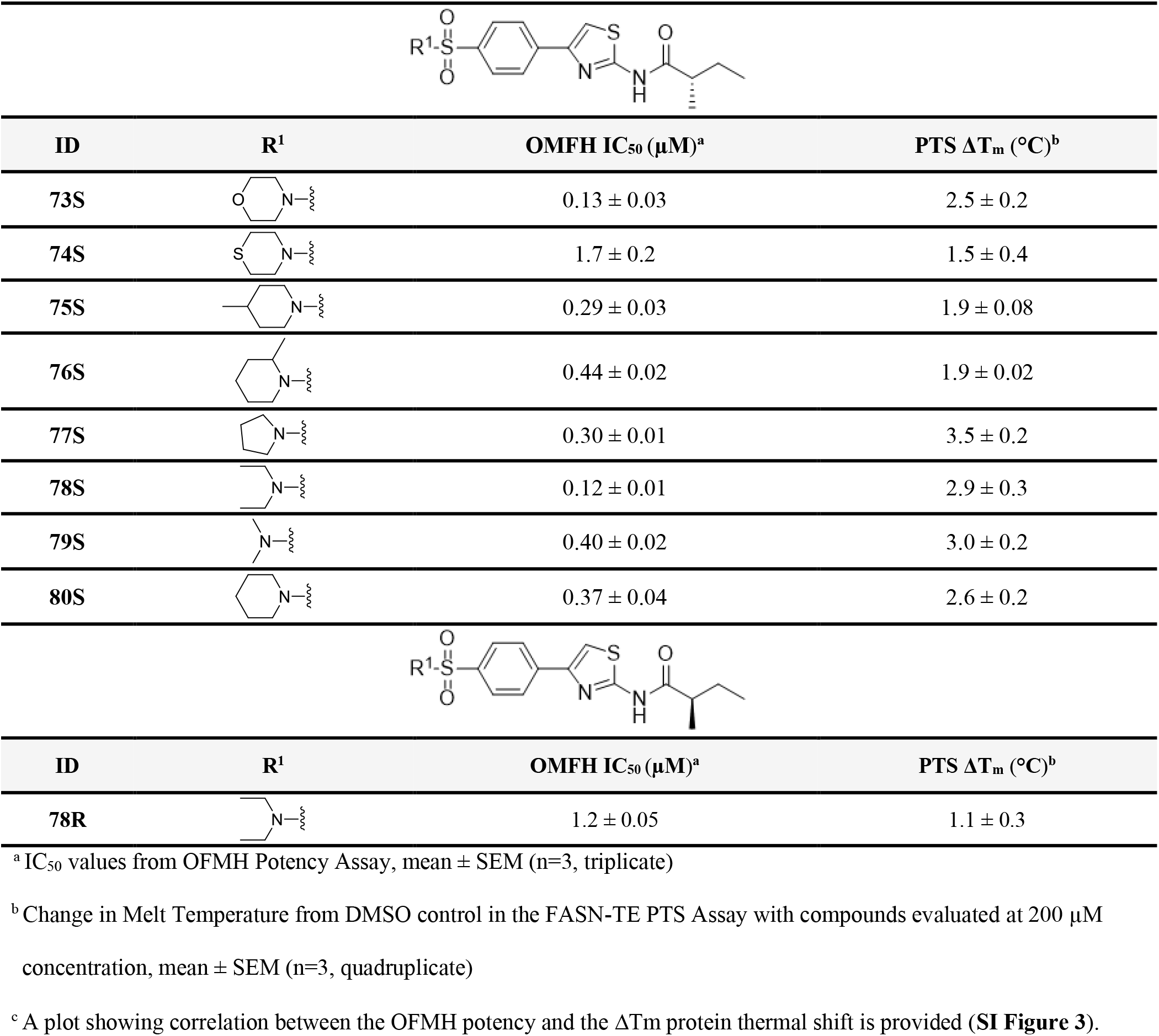
Stereopure inhibitors of FASN-TE, R^1^ biochemical potency and protein stability.

|  |  |  |  |
| --- | --- | --- | --- |
| ID | R<sup>1</sup> | OMFH $IC_{50}$ ($\mu M$)<sup>a</sup> | PTS $\Delta T_m$ ($^{\circ}C$)<sup>b</sup> |
| **73S** | | $0.13 \pm 0.03$ | $2.5 \pm 0.2$ |
| **74S** | | $1.7 \pm 0.2$ | $1.5 \pm 0.4$ |
| **75S** | | $0.29 \pm 0.03$ | $1.9 \pm 0.08$ |
| **76S** | | $0.44 \pm 0.02$ | $1.9 \pm 0.02$ |
| **77S** | | $0.30 \pm 0.01$ | $3.5 \pm 0.2$ |
| **78S** | | $0.12 \pm 0.01$ | $2.9 \pm 0.3$ |
| **79S** | | $0.40 \pm 0.02$ | $3.0 \pm 0.2$ |
| **80S** | | $0.37 \pm 0.04$ | $2.6 \pm 0.2$ |
| ID | R<sup>1</sup> | OMFH $IC_{50}$ ($\mu M$)<sup>a</sup> | PTS $\Delta T_m$ ($^{\circ}C$)<sup>b</sup> |
| **78R** | | $1.2 \pm 0.05$ | $1.1 \pm 0.3$ |

<sup>a</sup> $IC_{50}$ values from OFMH Potency Assay, mean ± SEM (n=3, triplicate)
<sup>b</sup> Change in Melt Temperature from DMSO control in the FASN-TE PTS Assay with compounds evaluated at $200 \mu M$ concentration, mean ± SEM (n=3, quadruplicate)
<sup>c</sup> A plot showing correlation between the OFMH potency and the $\Delta T_m$ protein thermal shift is provided (**SI Figure 3**).

### Microsomal Stability and Pharmacokinetic Properties of Arylthiazole FASN-TE Inhibitors

We examined the potential in vivo utility of a panel of potent FASN-TE inhibitors by evaluating their stability upon incubation in vitro with pooled mouse, rat, or human liver microsomes for 60 min (**Table 3**). In general, the FASN-TE inhibitors exhibited high to moderate stability in this assay. While **66** exhibited favorable stability in this assay with 100% remaining after 60 min in human microsomes, **50** was less stable with only 58.0% remaining. The stereopure dimethylamine **79S** possessed the least stability of those tested, with moderate stability against mouse and rat microsomes and low stability in human microsomes. Both **73S** and **75S** remained stable, with 100% retention in both rat and human microsomes. We also measured the plasma stability for this panel of inhibitors at 60 min (**Table 4**). Each compound exhibited a moderate to high degree of plasma stability. Notable was the reduced level of plasma stability for both the stereopure **73S** and **75S** with 49% and 52% remaining respectively in human plasma. The other stereopure compounds tested, however, were stable with 100% remaining.

**Table 3:** Microsomal Stability of Arylthiazole FASN-TE inhibitors.

| Microsomal Stability (Percent remaining at 60 min) |  |  |  |
| --- | --- | --- | --- |
| Compound ID | Mouse | Rat | Human |
| **50** | 63 | 96 | 58 |
| **66** | 100 | 86 | 100 |
| **73S** | 63 | 100 | 100 |
| **75S** | 100 | 100 | 100 |
| **77S** | 60 | 100 | 85 |
| **78S** | 72 | 99 | 56 |
| **78R** | 70 | 100 | 64 |
| **79S** | 52 | 65 | 0 |

**Table 4:**
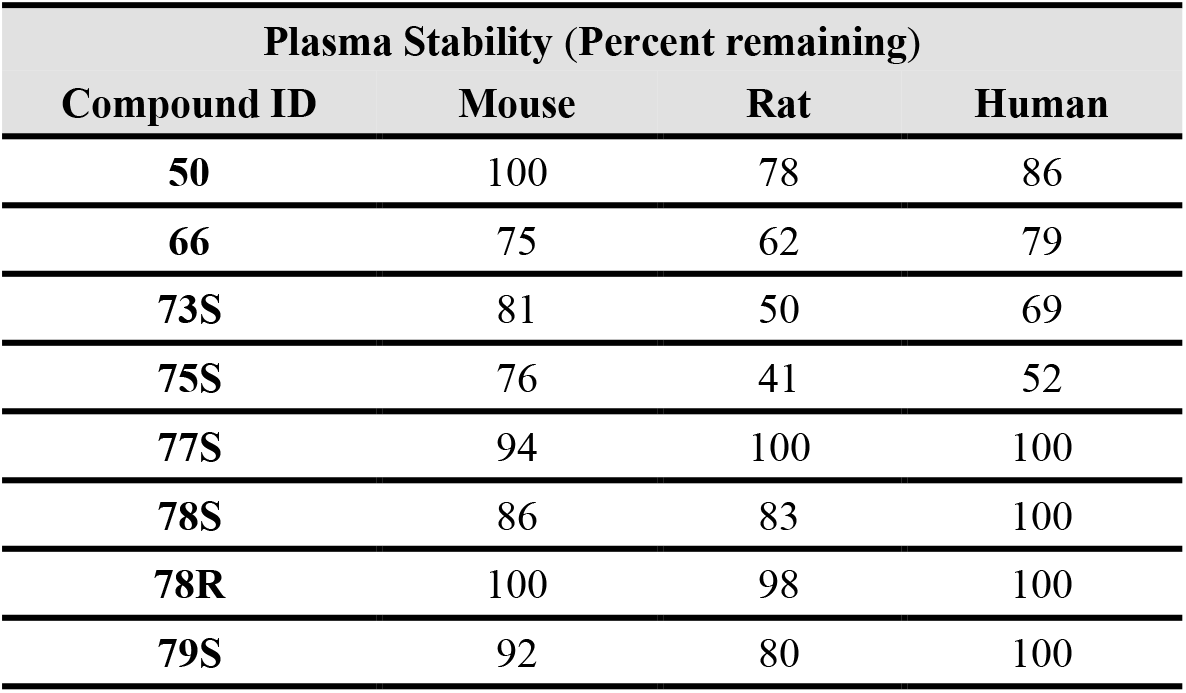
Plasma Stability of Arylthiazole FASN-TE inhibitors.

| Plasma Stability (Percent remaining) |  |  |  |
| --- | --- | --- | --- |
| Compound ID | Mouse | Rat | Human |
| **50** | 100 | 78 | 86 |
| **66** | 75 | 62 | 79 |
| **73S** | 81 | 50 | 69 |
| **75S** | 76 | 41 | 52 |
| **77S** | 94 | 100 | 100 |
| **78S** | 86 | 83 | 100 |
| **78R** | 100 | 98 | 100 |
| **79S** | 92 | 80 | 100 |

We then subjected this panel of compounds to in vivo analysis to evaluate their systemic exposure in C57BL/6J mice. We conducted a rapid pharmacokinetic assessment by oral gavage (Snapshot PK) of a cohort of mice dosed at 10 mg/kg and measured plasma concentrations at 30 and 60 min (**Table 5**). Plasma exposure for these FASN-TE inhibitors ranged widely in this preliminary evaluation. The stereopure **78S** demonstrated the highest overall exposure, with values of 1.35 µM at 30 min and 0.92 µM at 60 min. Compounds **66** and **75S** also showed moderate exposure with observed concentrations of 0.95 and 0.87 µM at 30 min and 1.1 and 0.88 µM at 60 min.

**Table 5:** In vivo snapshot pharmacokinetic exposure of arylthiazole FASN-TE inhibitors in C57BL/6J mice.

| Plasma Pharmacokinetics (nM) |  |  |
| --- | --- | --- |
| Compound ID | 30 min | 60 min |
| 50 | < 2 | < 2 |
| 66 | 950 ± 100 | 1100 ± 220 |
| 73S | 90 ± 40 | 50 ± 25 |
| 75S | 870 ± 150 | 880 ± 50 |
| 77S | 12 ± 0 | 15 ± 8 |
| 78S | 1350 ± 200 | 920 ± 80 |
| 78R | 250 ± 110 | 130 ± 30 |
| 79S | 70 ± 50 | 3 ± 1 |

To further quantify the potential of our arylthiazole inhibitors for in vivo studies we assessed the time-course PK profile of **78S** in C57BL/6J mice. Following a single dose of **78S** administered either intravenously (IV, 1.0 mg/kg) or by oral gavage (PO, 10 mg/kg), we measured the concentration of **78S** in plasma over a seven-point time-course (**Figure 2**). We observe a favorable profile for **78S** with a C_max_ of 2.56 µM following oral administration with an oral bioavailability of 84%. The pharmacokinetic properties of **66**, **75S**, and **78S** suggest a basis for improvement but are especially promising when compared to the existing FASN-TE inhibitor, Orlistat^54^. Importantly, the PK exposure data suggest that the new FASN-TE inhibitors may serve as viable in vivo tool compounds for the study of FASN.

**Figure 2:**
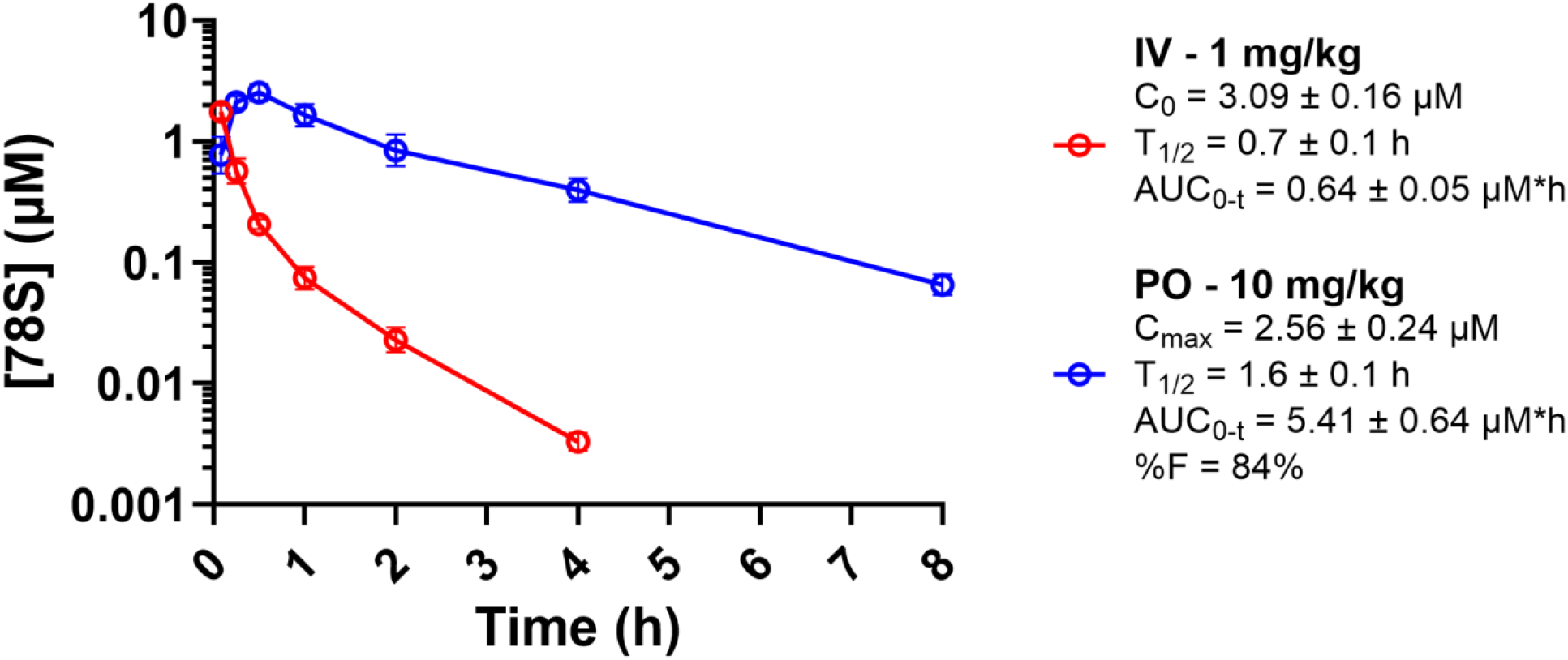
PK analyses of stereopure FASN-TE inhibitor 78S. For time-dependent PK studies, the stereopure arylthiazole **78S** was administered to mice both IV and PO. Male C57BL/6J mice (N=3) were treated with **78S** by a single dose via tail injection (IV; 1.0 mg/kg, red) or oral gavage (PO; 10 mg/kg, blue). Plasma concentrations for **78S** were obtained at the seven time points indicated as measured by LC-MS/MS. Data are presented as the geometric mean ± SD of the N=3 on a log10 scale and were analyzed using the Microsoft Excel plugin PKSolver 2.0 ^65^. C_0_, initial concentration; C_max_, the maximum plasma concentration post dosing; T_1/2_ terminal elimination half-life; AUC, area under the curve and %F, percentage bioavailability are provided.

FASN-TE inhibitor compounds were formulated as 5% DMSO, 10% TWEEN-80 and 85% water as 1 mg/mL stock solutions and orally administered to at 10 mg/kg to adult female C57BL/6J mice in a 10 mL/kg dosing volume. Mice (n=3) were dosed by oral gavage. Blood samples were collected at 30 min and 60 min post-treatment. The concentration of FASN-TE inhibitors was determined by LC-MS.

### Cellular Target Engagement of Stereopure Arylthiazole Inhibitors of FASN-TE

We developed a method to measure cellular target engagement with FASN-TE based on the principles of the cellular thermal shift assay (CETSA) ^66^. Focusing on FASN-TE allows for the detection of small thermal melt temperature changes induced by small molecules which would be masked in the full-length FASN homodimer. A plasmid encoding a tagged FASN-TE protein was transfected into HEK293 cells after which the cells were exposed to a thermal gradient to induce protein unfolding and depletion. The remaining soluble protein expression level of FASN-TE in cell lysates was quantified using a coupled chemiluminescent reporter system. Treatment with arylthiazole inhibitors produces a measurable shift in the thermal profile of soluble FASN-TE relative to DMSO (**Figure 3A**). This is correlated with the compound’s ability to affect the unfolding and precipitation of FASN-TE in the cell. The thermal profiles are quantified using differences in the melt temperature of the protein (ΔT_m_), as defined by the temperature at which 50% of the maximal signal is recorded (EC_50_), or the fold change in chemiluminescence at a single temperature (ΔL_Iso_). Optimization was performed to determine that 54°C was a reliable isothermal temperature to detect differences in cellular target engagement of the stereopure arylthiazole inhibitors.

**Figure 3:**
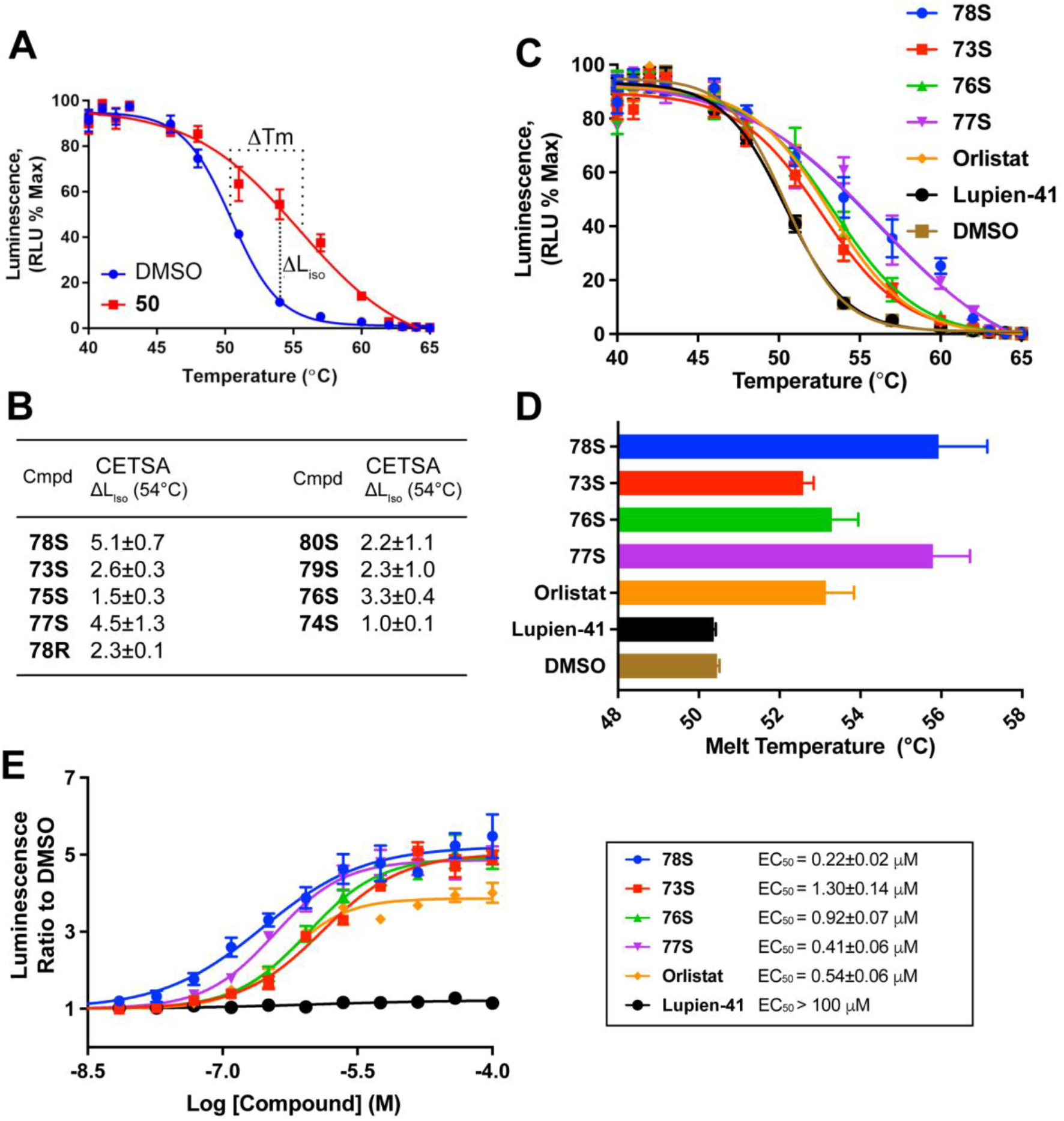
Stereopure arylthiazole inhibitors show target engagement with the TE domain of FASN in a cellular environment. (A) Representative thermal profiles of the TE domain of FASN transfected in HEK293 cells treated with vehicle (DMSO, blue) and an arylthiazole compound (**50,** (red) with relative ΔT and ΔL_Iso_, mean ± SEM (n=3, triplicate). (B) Table of the change in luminescence relative to DMSO control (ΔL_Iso_) of stereopure arylthiazoles treated at 10 µM, mean ± SEM (n=3, quintuplicate). (C) Effect of inhibitors at 100 µM for 1 h, on the thermal profile of the TE domain of FASN transfected into HEK293 cells, mean ± SEM (n=4, triplicate). (D) Quantification of the change in cellular melt temperature of the TE domain of FASN in the presence of inhibitors, calculated using curves shown in C, mean ± SEM (n=4, triplicate). (E) Dose-response of inhibitors bound to the TE domain of FASN transfected into HEK293 cells as measured by the change in signal relative to DMSO at 53°C, mean ± SEM (n=3, quintuplicate). Inset: Calculated EC_50_ values from E.

When the panel of stereopure FASN-TE inhibitors were profiled with CETSA to measure ΔL_Iso_ (**Figure 3B**), the general potency trends observed in the prior OMFH and PTS assays were also seen. This demonstrates that on-target activity is not hindered by other components of the cellular environment. **78S** was the most potent stereopure arylthiazole in the OMFH assay and also produced the largest stabilization increase in ΔL_Iso_, 5.1-times relative to DMSO. **78S** was once again found to be more active than the *R*-enantiomer **78R**, which only produced a fold change of 2.3, though, surprisingly this was still a stronger response than other stereopure compounds. For example, **75S**, which was 4-fold more potent than **78R** in the biochemical OMFH assay, only produced a ΔL_Iso_ fold change of 1.5-times DMSO. Also of interest, the pyrrolidine **77S**, which produced the largest PTS shift of 3.5°C, had the second largest ΔL_Iso_ of 4.5-times. Meanwhile, **76S**, which only produced a PTS shift of 1.9°C yielded a ΔL_Iso_ of 3.3-times that of DMSO. Stereopure arylthiazole compounds found to have a ΔL_Iso_ greater than a 2.5-fold difference in luminescence from DMSO with CETSA (**Figure 3B**) were subjected to more quantitative comparative affinity analysis to determine a CETSA ΔT_m_. Two previously published inhibitors were used to benchmark the stereopure arylthiazoles: Orlistat ^53^, and Compound 41 (Lupien-41) reported by Lupien et al. to be an inhibitor of FASN-TE ^36^ (**SI Figure 2**). Mechanistically, Orlistat binds irreversibly while Lupien-41 is a reversible inhibitor. We obtained Lupien-41 and characterized this compound using our primary testing strategy and determined it to possess an IC_50_ of 2.7 ± 0.62 µM using the OMFH biochemical assay and a moderately negative thermal shift of −0.91 ± 0.26°C using the FASN-TE PTS assay.

CETSA thermal profiles of the TE domain were generated with cells treated with 100 µM of each compound (**Figure 3C**). A concentration of 100 µM was used for the thermal profile comparison to ensure maximal signal, and that the dose-response CETSA (**Figure 3E**, EC_50_ = 220 nM for **78S**) confirms on-target activity at pharmacologically relevant concentrations. From the thermal profiles, the stabilizing effect on the melting temperature (T_m_) of the domain in the cellular environment was measured (**Figure 3D**). **78S** and **77S** were found to have the largest stabilizing effect on the melting temperature, with an increase of 5.9 and 5.3°C respectively over DMSO. The inhibitors **73S** and **76S** had roughly half the effect, similar to the results of Orlistat (2.7°C greater than vehicle). Lupien-41 had no effect on the melt temperature. We performed an isothermal dose-response titration of compounds at 53°C and found that compound **78S**, **73S**, **76S**, and **77S** all had EC_50_ values in the high nanomolar to low micromolar range (**Figure 3E**) with the most potent compound being **78S** (220 nM). Under these assay conditions, Orlistat also had an EC_50_ value in the high nanomolar range (540 nM) though, the maximum response generated by the highest concentration of Orlistat was < 80% of that observed for the arylthiazoles.

### Stereopure Arylthiazole Inhibitors Bind Reversibly to FASN-TE

We used a serine hydrolase activity-based probe TAMRA-FP, to gauge the reversibility of the stereopure arylthiazole inhibitors ^39^. The fluorophosphonate probe relies on serine hydrolase enzymatic activity to bind irreversibly and nonspecifically to members of the serine hydrolase family to which the TE domain of FASN belongs ^39^. The probe is unable to bind to proteins that lack serine hydrolase activity, or whose activity is inhibited by small molecule inhibition, or heat inactivation (HI). Bound TAMRA-FP produces a fluorescent signal that can be used to visualize labeled proteins by imaging of an SDS-PAGE gel (**Figure 4A**). For these experiments we also benchmarked our compounds against an analogue of the irreversible FASN-TE inhibitor IPI-9119 ^30^.

**Figure 4:**
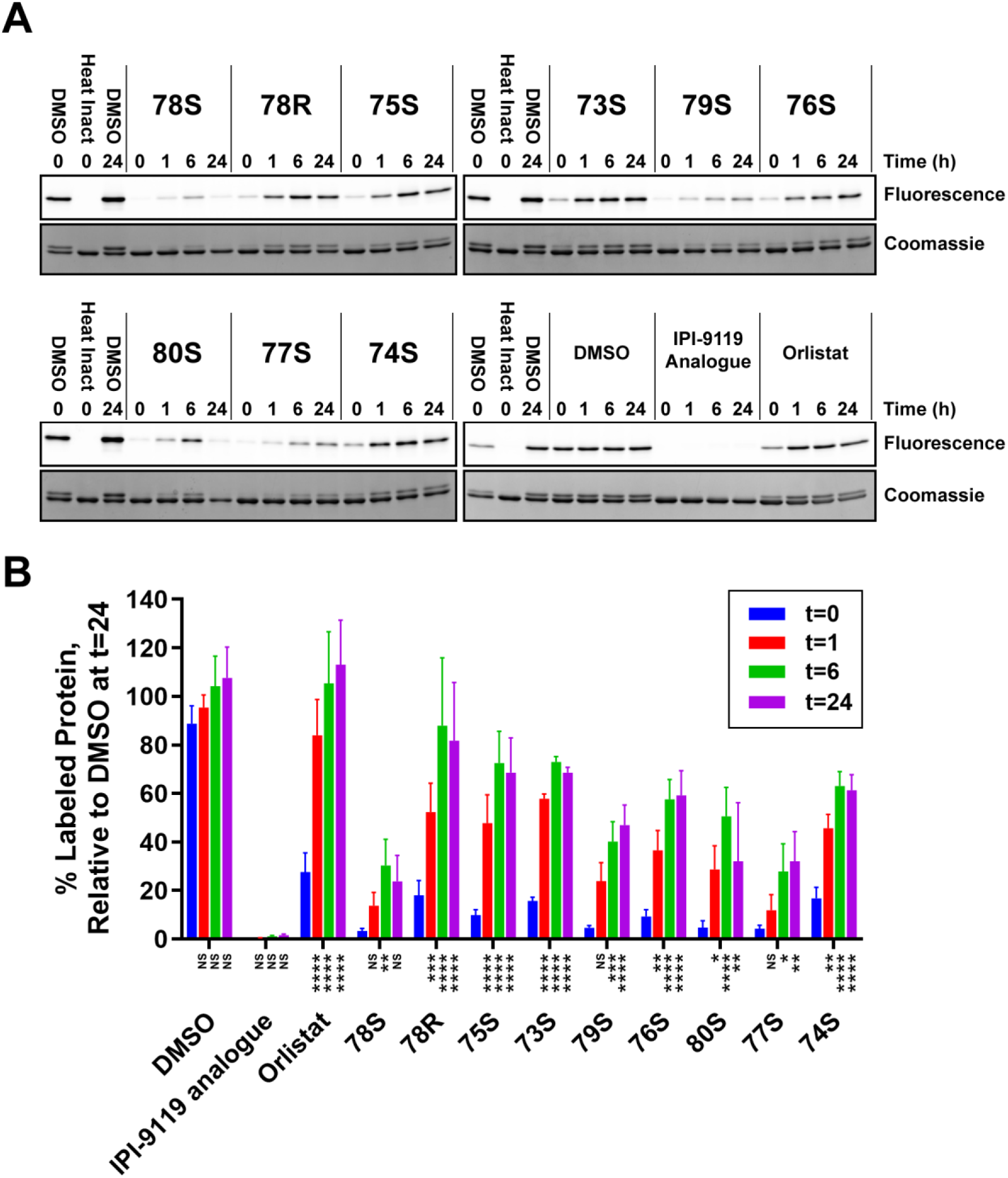
Arylthiazoles reversibly inhibit binding of the non-specific activity-based serine hydrolase probe ActivX TAMRA-FP. Arylthiazole and competitor inhibitors were tested at 200 µM. (A) A representative fluorescence image and Coomassie stained gel showing inhibition of TAMRA-FP binding to FASN-TE over a 24-hour time course. Fluorescent gels indicate probe-bound protein complexes while Coomassie stained gels demonstrate equal protein loading in all samples. (B) Quantification of TAMRA-FP fluorescent signal in A, mean ± SEM (n=3, singlicate). Statistical significance was calculated using 2-way ANOVA multiple comparisons test relative to the t = 0 timepoint for each treatment condition using GraphPad Prism, Version 11. *, p<0.05; **, p<0.01; ***, p<0.001; ****, p<0.0001.

At t=0 h, the stereopure arylthiazole inhibitors block >80% of the probe from binding to recombinantly purified FASN-TE relative to DMSO, with the most potent, **78S**, blocking 97%. The IPI analogue blocks 99% of binding, while orlistat only blocks 72% (**Figure 4B**). Over time, the probe binds to more protein, indicating the arylthiazole inhibitors are released, and therefore reversible. All arylthiazole analogues inhibit significantly less binding of the probe after 6 h. For example, immediately after the addition of **75S**, only 10% of protein is bound to the probe, relative to DMSO, whereas after 6 h, 73% of the probe is bound to FASN-TE. While **78S** continues to strongly inhibit the probe from binding after 6 h (30% bound, relative to DMSO), it still demonstrates release over time, up from an initial value of 3% protein bound to probe relative to DMSO. Alternatively, the IPI-9119 analogue, which binds irreversibly, persists at inhibiting 98% of the probe after 24 hours.

### Stereopure FASN-TE Inhibitors Serine Hydrolase Selectivity Profile

When we incubated TAMRA-FP with cellular lysates and analyzed the lysate via SDS-PAGE, the probe produced a signal at multiple bands consistent with the nonspecific, activity-based targeting of serine hydrolases in the lysate (**Figure 5A**). The source of cellular lysate chosen for these experiments was derived from SK-MEL5 cells, a melanoma cell line, on the basis that FASN inhibition has been determined to be a melanoma vulnerability and is therefore a potential therapeutic target ^67^. The probe-protein complex that produced the strongest signal was FASN (verified by Western blotting **SI Figure 6A**). Several other serine hydrolases were labeled by the probe with the next strongest signals arising from proteins with molecular weights of 81.5, 29, and 26 kDa (**SI Figure 6B**). In this competition assay all compounds tested, except for Lupien-41, inhibited >95% of the signal from FASN (**Figure 5B**). The nonselective IPI-9119 analogue also reduced labeling by the probe to all three of the other proteins analyzed. In contrast, the arylthiazoles were highly selective for FASN (**SI Figure 7**) with only a minor reduction in the labeling of TAMRA-FP to the other serine hydrolases. Using a similar activity-based probe strategy, we compared the selectivity of **50** for the FASN-TE when compared to recombinant acyl-coenzyme A thioesterase 4 (ACOT-4) (**SI Figure 8**). Arylthiazoles show no off-target binding activity even for closely related thioestease ACOT-4.

**Figure 5:**
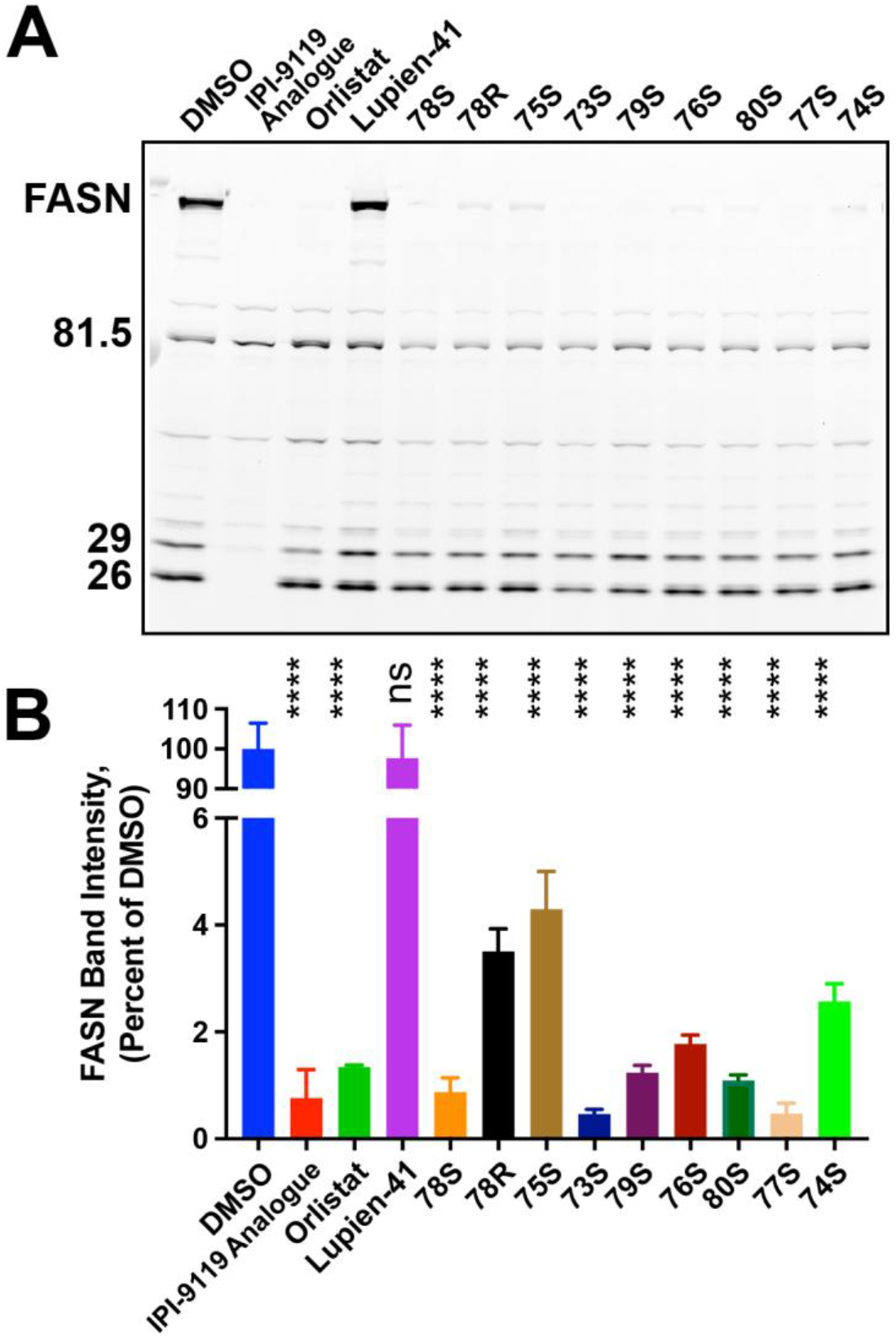
Arylthiazoles selectively inhibit FASN in cellular lysates. (A) SK-Mel5 lysate was incubated with inhibitors (200 µM) for 30 min before treatment with FP-TAMRA, the reaction was allowed to proceed for 10 min, and the reaction was quenched. A representative fluorescent gel shows the activity-based probe bound to multiple serine hydrolase proteins in the DMSO/control treated lysate. The relative ability of inhibitors to reduce the reactivity of the probe to these proteins is evident. (B) Quantification of fluorescent signal from FASN in (A), relative to the signal from DMSO, mean ± SEM (n=3, duplicate). Significance was calculated using Dunnett’s multiple comparisons test compared with vehicle (DMSO) control using GraphPad Prism, Version 11. *, p<0.05; **, p<0.01; ***, p<0.001; ****, p<0.0001.

**Figure 6:**
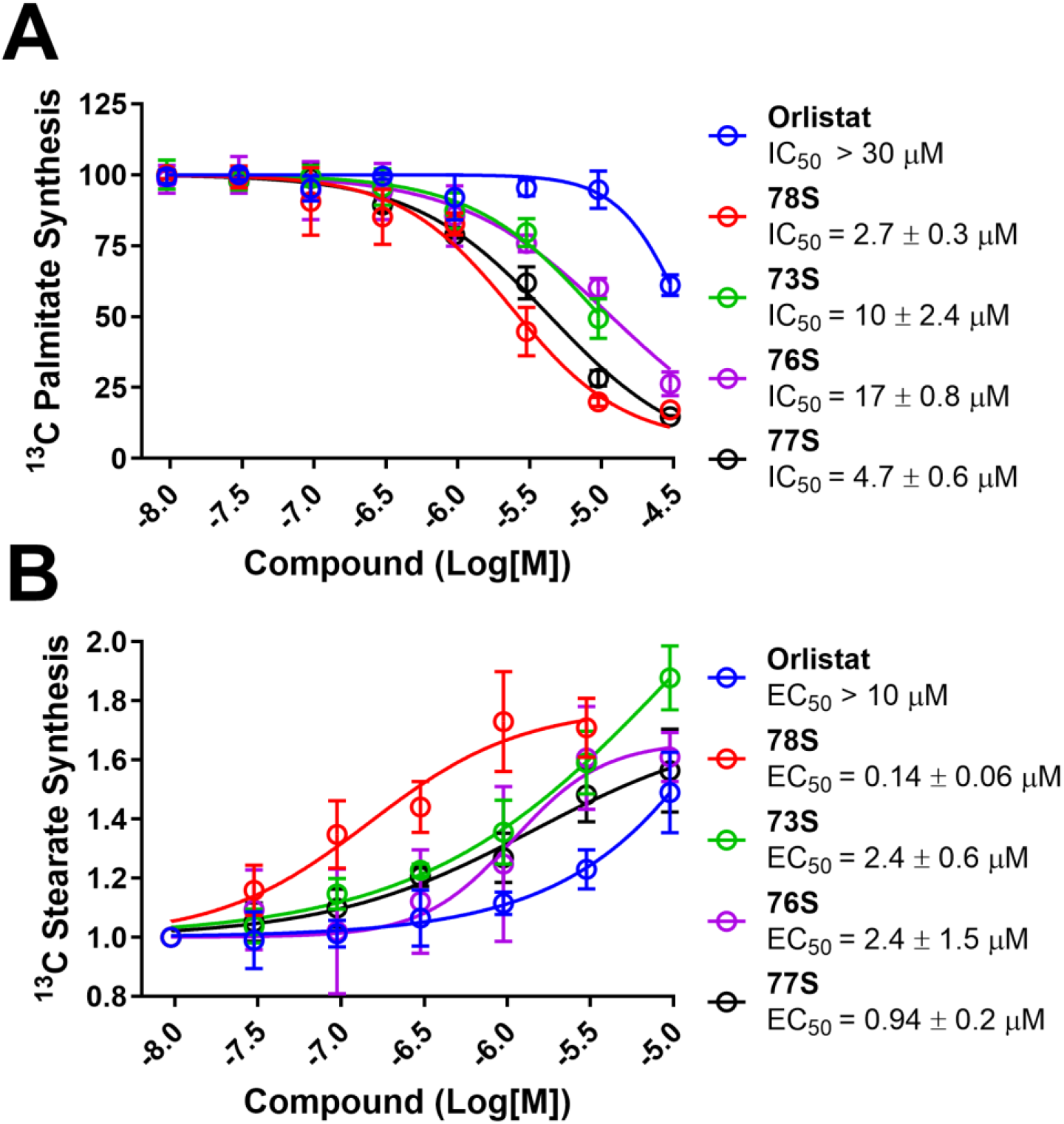
Stereopure arylthiazole FASN inhibitors block the production of palmitate in cells. (A) Dose-response inhibition of de novo palmitate synthesis in SK-Mel5 cells measured by mass-spectroscopy analysis of ^13^C incorporation into palmitate 72 h post treatment with ^13^C glucose and compound, mean ± SEM (n=3, duplicate). (B) Inhibition of the TE domain of FASN simultaneously induces the de novo synthesis of stearate, a fatty acid with 18 carbons, as measured by ^13^C incorporation of stearate in the same samples collected and analyzed in (A), mean ± SEM (n=3, duplicate).

Lupien-41 was previously reported to potently inhibit purified FASN-TE in biochemical assays yet exhibit only weak cytotoxicity in cancer cells, a discrepancy attributed to poor pharmaceutical properties rather than loss of target affinity ^36^. Consistent with this, the absence of a CETSA thermal shift and TAMRA-FP competition signal for Lupien-41 in our hands most likely reflects insufficient intracellular compound accumulation rather than a fundamentally different binding site on FASN-TE.

In addition, we evaluated the ability of **50** to inhibit a panel of proteases at Reaction Biology (Malvern, PA) (**SI Table 2**). For these studies, time-course measurements of the increase in fluorescence signal from fluorescently labeled peptide substrates were monitored and the initial linear portion of slope (signal/min) was analyzed. Compound **50** was inactive against all the proteases tested, except for weak activity at chymotrypsin (1.7 µM IC_50_). We conclude that compounds in this chemical series display highly limited off-target protease activity.

### Stereopure Arylthiazole FASN Inhibitors Inhibit Palmitate Synthesis but increase Stearate Synthesis

We also tested the ability of the arylthiazoles to inhibit the cellular catalytic activity of full length FASN by measuring de novo synthesis of palmitate. SK-Mel5 cells were grown in ^13^C-glucose enriched media with the incorporation of isotope into newly synthesized lipids quantified by mass spectroscopy. Dose dependent inhibition of palmitate synthesis by the arylthiazoles was measured in SK-Mel5 cells over 72-h (**Figure 6A**).

Inhibition of palmitate biosynthesis by the panel of FASN-TE inhibitors correlated well with the EC_50_ values recorded using the CETSA assay, with **78S** possessing the best potency (2.7 µM). The exception to this was Orlistat, which, while recording an EC_50_ value of 0.54 µM in the CETSA assay, had an IC_50_ value for the inhibition of cellular palmitate synthesis weaker than the experimental window could quantify.

Interestingly the inhibition of FASN-TE activity results in the de novo synthesis of stearic acid (**Figure 6B**). Stearic acid is an 18-carbon fatty acid and the observation of dose-dependent increase in its concentration is mechanistically consistent with the model that the FASN-TE inhibitors interfere with the size-selective role of FASN-TE. Inhibition of FASN-TE would be predicted to cause FASN to produce larger lipid products; it has been shown that when the TE domain of FASN is enzymatically cleaved from the rest of the protein, fatty acids of up to 22 carbons are synthesized ^68^. Our findings demonstrate that small molecule inhibition of the TE domain mimics the cleaved forms of FASN.

Because the background level of stearate synthesized by FASN is significantly lower than palmitate in the cells, detection is much more sensitive to small changes. This results in a several fold lower EC_50_ value for the production of stearate than the IC_50_ value determined for the inhibition of palmitate. For the production of stearate, **78S** remained the most potent compound, with a recorded EC_50_ value of 0.14 µM demonstrating that the high inhibitory potency correlates with this detected activity. The rank order for the other stereopure arylthiazoles tested was consistent with both inhibition of palmitate synthesis and stearate production. Notably, chemical probe **78S** has a biochemical IC_50_ of 0.12 µM against recombinant FASN-TE but a cellular IC_50_ of 2.7 µM for palmitate synthesis inhibition. This discrepancy may be due to competition with the natural palmitoyl-ACP substrate at physiological concentrations, intracellular protein binding, or the high intracellular FASN concentration in the cellular assay relative to the recombinant assay, or contributions from all three factors.

The combined inhibition of palmitate synthesis and induction of stearate synthesis distinguishes the mode of action of our TE domain inhibitors from those that target either the KS or KR domains. Dietary stearate has been shown to induce apoptosis in cancer cells and reduce metastasis burden in murine models, purportedly through inhibition of cell cycle checkpoints ^69–74^, suggesting that endogenously synthesized stearate may contribute to the anti-tumor activity of TE domain inhibitors, a mechanistic question that, to our knowledge, has not yet been directly investigated. Furthermore, the mechanistically distinct lipid remodeling signature of FASN-TE inhibition, the simultaneous reduction of palmitate and dose-dependent induction of stearate, may have therapeutic relevance in MASH, where palmitate-driven lipotoxicity is a primary driver of hepatocyte injury, inflammation, and fibrosis ^75^. **SBP-5917** thus provides a structurally validated chemical probe to investigate the consequences of selective TE-domain inhibition in both cancer and metabolic liver disease models.

## DISCUSSION AND CONCLUSIONS

We have designed, synthesized and characterized a series of stereopure arylthiazole small molecule inhibitors of the TE domain of FASN. Our best compound, stereopure **SBP-5917** (**78S**), is potent yet reversible, and possesses both on-target affinity and efficacy in cellular models. The testing funnel we employed, beginning with screens of compound potency and binding affinity against the recombinantly purified TE domain of FASN followed by confirmation of cellular target engagement, led to a series of improved lead compounds. The crystal structures we solved indicate that the potent activity of **SBP-5917** is due to its binding within the substrate tunnel. Moreover, competition with an activity-based probe indicates that arylthiazole inhibition is reversible, a potentially desirable drug-like property. This activity probe also provides evidence that **SBP**-**5917** binds to the full length FASN protein in addition to the recombinantly purified TE domain. This finding is further validated through measurement of de novo palmitate synthesis and efficacy against full length FASN in cells.

The structural enablement insights provided by the bound structures of **66** and **71,** which we advanced to the stereopure **SBP-5917**, invites some evaluation of the determinants of binding. At the resolution of the co-crystal structures, we observe that polar interactions to S2308 and I2250 to be largely water-mediated interactions. Polar water networks are not to be thought of as mere “filler”; greater selectivity and potency can sometimes be achieved by designing ligands to induce bridging mimicry and/or strategic displacement of high energy waters ^76^. Furthermore, the chiral N-acyl 2-methyl butanamide substituent which extends into the substrate tunnel invites further optimization as this is indicative of a shape-selective bound conformation that not an isostere of a flexible fatty acid side chain. Access to this property was not obvious in previous structural studies ^57^. With the aim of improving our FASN-TE inhibitors, we plan to develop this series further.

The stereopure arylthiazole inhibitor we have described and characterized is a new chemical probe that can be used to investigate the function of the TE domain of FASN. **SBP-5917** binds to the substrate pocket of the TE domain, possesses promising pharmacokinetic properties, and allows the inhibition of both biochemical and cellular forms of FASN. **SBP-5917** should serve as a useful cellular and in vitro probe of FASN and can guide the future development of FASN inhibitors that target FASN dependent diseases.

## EXPERIMENTAL SECTION

### Chemical methods

Commercial sourced sources were the origin of all reagents and were used without further purification unless otherwise stated. All reactions were performed under dry N_2_ unless otherwise stated. Chromatographic purification of compounds used several methods. Compound purification by preparative HPLC was performed on a Waters purification system C18 column (reverse phase, 50 X 150 mm, 5 μm) with an ESI mass spectrometer. Flash chromatography was performed using either; a WelFlash SiO2-II or irregular silica gel (40–63 μm) column cartridges and compounds were eluted using a Biotage Sweden AB box B, or RediSep columns using a Teledyne ISCO.

Purity of compounds was established by high-pressure liquid chromatography (HPLC) with UV detection and liquid chromatography-mass spectroscopy (LC-MS). Characterization and analysis by nuclear magnetic resonance (NMR) was performed using a JEOL400 spectrophotometer which was used to obtain 1H and 13C spectra at 400 MHz and 101 MHz respectively. Chemical shifts are reported in δ (ppm) referenced to tetramethylsilane, proton integration, multiplicity and coupling constants (Hz). NMR spectra were analyzed using MestReNova version 14.3.0.

The purity of final products for biological testing in vivo and in cells was determined by analytical HPLC-MS to be greater than 95%. HPLC analysis was performed on a Shimadzu 2010 EV LC-MS using the following conditions: Kromasil C18 column (reverse phase, 4.6 mm, 50 mm). Chiral characterization HPLC and High-resolution ESI-TOF mass spectra were provided by the Scripps Research Institute Center for Metabolomics and Mass Spectrometry (La Jolla, CA, USA).

### N-(4-(4-(N,N-diethylsulfamoyl)phenyl)thiazol-2-yl)-2-methylbutanamide (50)

A solution of 4-(2-Amino-1,3-thiazol-4-yl)-N,N-diethylbenzenesulfonamide (**83**) (2.0 g, 9.5 mmol) in dry DCM (100 mL) was provided with 2-methylbutanoyl chloride (1.2 g, 9.5 mmol) and pyridine (1.12, 14.2 mmol). The reaction was stirred for 2 h at ambient temperature. The reaction was quenched and extracted with EA/brine. The organic layer was dried over Na_2_SO_4_ and concentrated to yield the crude product. The compound was purified by flash column chromatography (silica gel, eluting with 33% to 50% EA/hexanes). The purified yield of (**50**) was (1.03 g, 27%) as a white solid. HPLC/LC purity > 95%. LC-MS (ESI): 396.33 [M+H]^+^. ^1^H NMR (400 MHz, CHLOROFORM-*D*) δ 9.66 (s, 1H), 7.96 (d, J = 8 Hz, 2H), 7.93 (d, J = 8 Hz, 2H), 7.27 (s, 1H), 3.26 (dd J = 6.8 Hz, J = 6.8 Hz, 4H), 2.44 (m, 1H), 1.80 (m, 1H), 1.56 (m, 1H), 1.23 (m, 3H), 1.14 (m, 6H), 0.94 (m, 3H). ^13^C NMR (101 MHz, CHLOROFORM-*D*) δ 174.90, 158.52, 148.01, 139.26, 138.20, 127.70, 126.55, 110.22, 42.78, 42.13, 27.19, 17.02, 14.20, 11.81. HRMS (ESI-TOF) calcd for C_18_H_25_N_3_O_3_S_2_ [M + H]^+^, 396.1337; found 396.1365.

### 4-fluoro-N-(4-(4-(morpholinosulfonyl)phenyl)thiazol-2-yl)benzamide (66)

A solution of 2-Bromo-1-[4-(4-morpholinylsulfonyl)phenyl]ethanone (88 mg, 0.25 mmol) in THF was combined with 4- Fluorobenzenecarbothioamide (68 mg, 0.44 mmol) and sodium bisulfate monohydrate (38 mg, 0.28 mmol) in 10 mL THF. The reaction was allowed to proceed for 16 h at ambient temperature. The THF was removed in vacuo and the solids dissolved in DCM, washed against aqueous solution and dried over Na_2_SO_4_. The compound was purified by preparative HPLC and concentrated to yield (**66**) (68.3 mg, 0.17 mmol) in 67% yield as a white powder. ^1^H NMR (400 MHz, CDCl_3_) δ, ppm: 8.17 (d, 2H), 8.03 (dd. 2H), 7.83 (d, 2H), 7.64 (s, 1H), 7.18 (t, 2H), 3.75 (dd, 4H), 3.04 (dd, 4H). ^13^C NMR (400 MHz, CDCl_3_) δ, ppm: 167.45, 165.40, 162.82, 154.28, 138.72, 134.19, 128.66, 128.56, 128.41, 126.92, 116.33, 116.01, 115.15, 66.11, 46.02. HRMS (ESI): *m/z* calcd for C_19_H_17_FN_2_O_3_S_2_ 404.48, found (M + H)^+^ 405.08.

### *N*-(4-(4-(*N*,*N*-diethylsulfamoyl)phenyl)thiazol-2-yl)-2-(2-fluorophenyl)acetamide (71)

A solution of 4-(Bromoacetyl)-N,N-diethylbenzenesulfonamide (72.5 mg, 0.22 mmol) in dry THF (2.2 mL) was provided with Na_2_HSO_4_ (33 mg, 0.24 mmol) and 2-(2-Fluorophenyl)ethanethioamide (64 mg, 0.38 mmol). The reaction was stirred for 12 h at ambient temperature. The THF was removed in vacuo and dissolved in DCM. Extraction against H_2_O and then brine provided an organic fraction which was dried over Na_2_SO_4_ and then concentrated. The compound (**71**) was purified by flash column chromatography (silica gel, eluting with 25% EA in hexanes). The isolated white solid yield was 22.9 mg (97%). ^1^H NMR (400 MHz, CDCl_3_) δ, ppm: 8.02 (d, 2H), 7.86 (d, 2H), 7.64 (s, 1H), 7.51 (m, 4H), 4.45 (s, 2H), 3.26 (m, 4H), 1.13 (m, 6H). ^13^C NMR (400 MHz, CDCl_3_) δ, ppm: 169.32, 153.71, 139.54, 138.35, 137.89, 132.42, 129.30, 127.51, 126.74, 125.76, 124.13, 115.31, 41.95, 39.36, 14.06. HRMS (ESI): *m/z* calcd for C_20_H_21_FN_2_O_2_S_2_ 404.52, found (M + H)^+^ 405.11.

### (*S*)-2-methyl-*N*-(4-(4-(morpholinosulfonyl)phenyl)thiazol-2-yl)butanamide (73S)

To a solution of (*S*)-2-methylbutanoyl chloride (55.5 mg, 0.46 mmol) in dry-DCM (10 mL) were added 4-(4- (morpholinosulfonyl)phenyl)thiazol-2-amine (**100**) (100 mg, 0.31 mmol) and *N*,*N*-Diisopropylethylamine (119.3 mg, 0.92 mmol). The mixture was stirred at 55°C for 18 h under N_2_. After cooling, the mixture was concentrated to obtain a crude product which was purified by preparative-HPLC to afford the compound *(S)*-2-methyl-*N*-(4-(4-(morpholinosulfonyl) phenyl)thiazol-2-yl)butanamide (**73S**) (32.8 mg, 26.1%) as a white solid HPLC/UV purity > 98% (214.254 nm); LC-MS (ESI): 410.2 (M + H)^+^. ^1^H NMR (400 MHz, CDCL_3_-*D*) δ 9.85 (s, 1H), 8.00 (d, *J* = 8.2 Hz, 2H), 7.82 (s, 1H), 7.31 (s, 1H), 3.79 – 3.72 (m, 5H), 3.04 (t, *J* = 4.7 Hz, 5H), 2.53 – 2.45 (m, 1H), 1.83 (dt, *J* = 14.3, 7.3 Hz, 1H), 1.62 (dt, *J* = 13.8, 7.0 Hz, 1H), 1.30 (d, *J* = 6.9 Hz, 2H), 0.98 (t, *J* = 7.4 Hz, 4H). ^13^C NMR (101 MHz, CDCL_3_-*D*) δ 174.61, 128.58, 126.53, 110.30, 66.10, 46.01, 42.95, 27.01, 16.82, 11.70. HRMS (ESI-TOF) *m/z* calcd for C_18_H_23_N_3_O_4_S_2_ [M + H]^+^, 410.1130; found 410.1206.

### (*S*)-2-methyl-*N*-(4-(4-(thiomorpholinosulfonyl)phenyl)thiazol-2-yl)butanamide (74S)

To a solution of (*S*)-2-methylbutanoyl chloride (88 mg, 0.73 mmol) in dry-DCM (10 mL) were added 4-(4- (thiomorpholinosulfonyl)phenyl)thiazol-2-amine (**101**) (125 mg, 0.37 mmol) and *N*,*N*-Diisopropylethylamine (142.2 mg, 1.10 mmol). The mixture was stirred at 55°C for 18 h under N_2_. After cooling, the mixture was concentrated to give the crude product which was purified by prep-HPLC to afford the compound (*S*)-2-methyl-*N*-(4-(4-(thiomorpholinosulfonyl)phenyl) thiazol-2-yl)butanamide (5.1 mg, 3.3%) as a white solid HPLC/UV purity > 98% (214.254 nm); LC-MS (ESI): 426.2 (M + H)^+^. ^1^H NMR (400 MHz, CHLOROFORM-*D*) δ 11.35 (s, 1H), 7.97 (d, *J* = 8.4 Hz, 2H), 7.86 (d, *J* = 8.5 Hz, 2H), 7.30 (s, 1H), 3.06 – 2.99 (m, 4H), 2.64 (m, 6H), 1.87 (dt, *J* = 14.4, 7.3 Hz, 1H), 1.72 – 1.61 (m, 4H), 1.45 (td, *J* = 6.4, 3.6 Hz, 2H), 1.32 (s, 1H), 1.00 (t, *J* = 7.5 Hz, 2H). ^13^C NMR (101 MHz, CHLOROFORM-*D*) δ 175.25, 161.90, 128.79, 126.57, 109.16, 46.98, 42.94, 40.99, 26.68, 25.15, 23.45, 16.44, 11.53. HRMS (ESI-TOF) calcd for C_18_H_23_N_3_O_3_S_3_ [M + H]^+^, 426.0901; found 426.0969.

### (*S*)-2-methyl-*N*-(4-(4-((4-methylpiperidin-1-yl)sulfonyl)phenyl)thiazol-2-yl)butanamide (75S)

To a solution of (*S*)-2-methylbutanoyl chloride (72.1 mg, 0.6 mmol) in dry-DCM (10 mL) were added 4-(4-((4-methylpiperidin-1-yl)sulfonyl)phenyl)thiazol-2-amine (**102**) (135 mg, 0.40 mmol) and *N*,*N*-Diisopropylethylamine (155.3 mg, 1.2 mmol). The mixture was stirred at 55°C for 18 h under N_2_. After cooling, the mixture was concentrated to give the crude product which was purified by preparative-HPLC to afford the compound (*S*)-2-methyl-*N*-(4-(4-((4-methylpiperidin-1-yl)sulfonyl)phenyl)thiazol-2-yl)butanamide (47.3 mg, 28.0%) as a white solid HPLC/UV purity > 98% (214.254 nm); LC-MS (ESI): 422.2 (M + H)^+^. ^1^H NMR (400 MHz, CHLOROFORM-*D*) δ 7.97 (s, 7H), 7.31 (s, 1H), 7.27 (s, 2H), 3.28 (td, *J* = 5.8, 3.0 Hz, 7H), 2.79 – 2.71 (m, 2H), 1.84 – 1.76 (m, 3H), 1.36 (dd, *J* = 8.3, 6.6 Hz, 4H). ^13^C NMR (101 MHz, CHLOROFORM-*D*) δ 198.51, 174.68, 128.44, 126.40, 109.89, 46.47, 42.95, 33.32, 30.13, 26.98, 21.46, 16.78, 11.68. HRMS (ESI-TOF) calcd for C_20_H_27_N_3_O_3_S_2_ [M + H]^+^, 422.1493; found 422.1568.

### (2S)-2-methyl-N-(4-(4-((2-methylpiperidin-1-yl)sulfonyl)phenyl)thiazol-2-yl)butanamide (76S)

To a solution of (*S*)-2-methylbutanoyl chloride (92.6 mg, 0.77 mmol) in dry-DCM (10 mL) were added 4-(4-((2-methylpiperidin-1-yl)sulfonyl)phenyl)thiazol-2-amine (**103**) (130 mg, 0.39 mmol) and *N*,*N*-Diisopropylethylamine (149.6 mg, 1.16 mmol). The mixture was stirred at 55°C for 18 h under N_2_. After cooled, the mixture was concentrated to obtain the crude product which was purified by prep-HPLC to afford the compound (2*S*)-2-methyl-*N*-(4-(4-((2-methylpiperidin-1-yl)sulfonyl)phenyl)thiazol-2-yl)butanamide (89.1 mg, 59.7%) as a white solid HPLC/UV purity > 98% (214.254 nm); LC-MS (ESI): 422.2 (M + H)^+^. ^1^H NMR (400 MHz, CHLOROFORM-*D*) δ 10.79 (s, 1H), 7.97 – 7.87 (m, 4H), 7.28 (s, 1H), 4.28 (d, *J* = 6.8 Hz, 1H), 3.79 – 3.71 (m, 1H), 3.02 (td, *J* = 13.1, 2.7 Hz, 1H), 2.56 (p, *J* = 7.0 Hz, 1H), 1.89 – 1.81 (m, 1H), 1.68 – 1.49 (m, 6H), 1.46 (s, 1H), 1.32 (d, *J* = 6.9 Hz, 2H), 1.28 (d, *J* = 6.9 Hz, 0H), 1.10 (s, 2H), 1.08 (s, 2H), 0.99 (t, *J* = 7.3 Hz, 3H). ^13^C NMR (101 MHz, CHLOROFORM-*D*) δ 174.85, 127.86, 126.48, 109.41, 48.69, 42.97, 40.43, 30.36, 26.87, 25.25, 18.11, 16.67, 15.50, 11.62. HRMS (ESI-TOF) calcd for C_20_H_27_N_3_O_3_S_2_ [M + H]^+^, 422.1494; found 422.1559.

### (*S*)-2-methyl-*N*-(4-(4-(pyrrolidin-1-ylsulfonyl)phenyl)thiazol-2-yl)butanamide (77S)

To a solution of (*S*)-2-methylbutanoyl chloride (233 mg, 1.94 mmol) in dry-DCM (10 mL) were added 4-(4-(pyrrolidin- 1-ylsulfonyl)phenyl)thiazol-2-amine (**104**) (400 mg, 1.29 mmol) and *N*,*N*-Diisopropylethylamine (501.9 mg, 3.88 mmol) at 0 °C. The mixture was stirred at 55°C for 18 h under N_2_. After cooled, the mixture was concentrated to give the crude product which was purified by prep-HPLC to afford the compound *N*-(4-(4-(pyrrolidin-1-ylsulfonyl)phenyl)thiazol-2-yl)-2,3-dihydrobenzo[*b*][1,4]dioxine-2-carboxamide (298.4 mg, 58.7%) as a white solid HPLC/UV purity > 98% (214.254 nm); LC-MS (ESI): 393.9 (M + H)^+^. ^1^H NMR (400 MHz, CHLOROFORM-*D*) δ 10.91 (s, 1H), 8.01 – 7.90 (m, 3H), 7.31 – 7.25 (m, 1H), 3.29 (tq, *J* = 6.7, 2.7 Hz, 4H), 2.73 (s, 4H), 2.59 (qd, *J* = 6.9, 2.1 Hz, 1H), 1.89 – 1.74 (m, 4H), 1.71 – 1.57 (m, 1H), 1.30 (ddd, *J* = 16.5, 7.0, 2.2 Hz, 3H), 0.98 (qd, *J* = 7.0, 2.0 Hz, 3H). ^13^C NMR (101 MHz, CHLOROFORM-*D*) δ 174.93, 160.30, 154.05, 128.40, 127.68, 126.53, 109.56, 48.02, 42.93, 42.65, 27.06, 26.86, 25.29, 16.85, 16.65, 11.69, 11.61. HRMS (ESI-TOF) calcd for C_18_H_25_N_3_O_3_S_2_ [M + H]^+^, 394.1181; found 394.1254.

### (*R*)-*N*-(4-(4-(*N*,*N*-diethylsulfamoyl)phenyl)thiazol-2-yl)-2-methylbutanamide (78R)

To a solution of (*R*)-2-methylbutanoyl chloride (250 mg, 2.08 mmol) in dry-DCM (10 mL) were added 4-(2-aminothiazol-4-yl)-N,N-diethylbenzenesulfonamide (**105**) (431.9 mg, 1.39 mmol) and Pyridine (0.5 mL). The mixture was stirred at 60°C for 18 h under N_2_. After cooling, the mixture was concentrated to give the crude product which was purified by prep-HPLC to afford the compound (*R*)-*N*-(4-(4-(*N*,*N*- diethylsulfamoyl)phenyl)thiazol-2-yl)-2-methylbutanamide (85.4 mg, 15.0%) as a white solid HPLC/UV purity > 98% (214.254 nm); the enantiomeric excess was determined by chiral HPLC analysis (89.8%); (LC-MS (ESI): 396.0 (M + H)^+^. ^1^H NMR (400 MHz, CHLOROFORM-*D*) δ 10.49 (s, 1H), 7.97 – 7.87 (m, 4H), 7.27 (d, *J* = 3.6 Hz, 1H), 3.27 (q, *J* = 7.2 Hz, 4H), 2.53 (q, *J* = 6.9 Hz, 1H), 1.61 (dt, *J* = 14.0, 7.1 Hz, 1H), 1.32 – 1.23 (m, 3H), 1.19 – 1.13 (m, 5H), 1.12 (s, 2H), 0.99 – 0.91 (m, 3H). ^13^C NMR (101 MHz, CHLOROFORM-*D*) δ 174.84, 159.71, 140.30, 128.42, 127.82, 127.15, 126.50, 109.67, 42.88, 42.13, 42.08, 27.06, 26.94, 16.84, 16.73, 14.20, 14.14, 11.69, 11.64.

### (*S*)-*N*-(4-(4-(*N*,*N*-diethylsulfamoyl)phenyl)thiazol-2-yl)-2-methylbutanamide (78S)

To a solution of (*S*)-2-methylbutanoyl chloride (480 mg, 4.0 mmol) in dry-DCM (10 mL) were added 4-(2-aminothiazol- 4-yl)-N,N-diethylbenzenesulfonamide (**105**) (622 mg, 2.0 mmol) and Pyridine (740 mg, 10.0 mmol). The mixture was stirred at 60°C for 18 h under N_2_. After cooling, the mixture was concentrated to get a crude product which was purified by prep-HPLC to afford the compound (*S*)-*N*-(4-(4-(*N*,*N*- diethylsulfamoyl)phenyl)thiazol-2-yl)-2-methylbutanamide (403.5 mg, 51.1%) as a white solid HPLC/UV purity > 98% (214.254 nm); the enantiomeric excess was determined by chiral HPLC analysis (97.7%); LC-MS (ESI): 396.2 (M + H)^+^. ^1^H NMR (400 MHz, CHLOROFORM-*D*) δ 10.26 (s, 1H), 7.94 (qd, *J* = 8.7, 2.0 Hz, 5H), 3.27 (qd, *J* = 7.6, 2.0 Hz, 5H), 1.24 (dd, *J* = 7.0, 2.1 Hz, 2H), 1.14 (td, *J* = 7.3, 1.8 Hz, 6H), 0.97 – 0.89 (m, 4H). ^13^C NMR (101 MHz, CHLOROFORM-*D*) δ 198.15, 174.96, 159.04, 146.82, 139.55, 137.21, 127.70, 126.49, 109.99, 42.68, 42.06, 27.06, 16.84, 14.13, 11.69. HRMS (ESI-TOF) calcd for C_18_H_25_N_3_O_3_S_2_ [M + H]^+^, 396.1337; found 396.1361.

### (*S*)-*N*-(4-(4-(*N*,*N*-dimethylsulfamoyl)phenyl)thiazol-2-yl)-2-methylbutanamide (79S)

To a solution of (*S*)-2-methylbutanoyl chloride (97.5 mg, 0.81 mmol) in dry-DCM (10 mL) were added 4-(2-aminothiazol-4-yl)-*N*,*N*-dimethylbenzenesulfonamide (**106**) (115 mg, 0.41 mmol) and *N*,*N*-Diisopropylethylamine (157.5 mg, 1.22 mmol). The mixture was stirred at 55°C for 18 h under N_2_. After cooled, the mixture was concentrated to obtain the crude product which was purified by prep-HPLC to afford the compound (*S*)-*N*-(4-(4-(*N*,*N*-dimethylsulfamoyl)phenyl)thiazol-2-yl)-2-methylbutanamide (89.1 mg, 59.7%) as a white solid HPLC/UV purity > 98% (214.254 nm); LC-MS (ESI): 368.2 (M + H)^+^. ^1^H NMR (400 MHz, CHLOROFORM-*D*) δ 10.15 (s, 2H), 8.03 – 7.96 (m, 3H), 7.90 – 7.84 (m, 3H), 7.30 (s, 2H), 2.74 (s, 1H), 2.55 – 2.47 (m, 1H), 1.86 – 1.80 (m, 1H), 1.65 – 1.56 (m, 1H), 1.28 (d, *J* = 6.9 Hz, 3H), 0.97 (t, *J* = 7.5 Hz, 5H). ^13^C NMR (101 MHz, CHLOROFORM-*D*) δ 174.79, 159.28, 145.97, 136.89, 135.01, 128.50, 126.48, 110.06, 42.87, 37.97, 27.00, 16.81, 11.68. HRMS (ESI-TOF) calcd for C_16_H_21_N_3_O_3_S_2_ [M + H]^+^, 368.1024; found 368.1094.

### (*S*)-2-methyl-*N*-(4-(4-(piperidin-1-ylsulfonyl)phenyl)thiazol-2-yl)butanamide (80S)

To a solution of (*S*)-2-methylbutanoyl chloride (278.6 mg, 2.32 mmol) in dry-DCM (10 mL) were added 4-(4-(piperidin-1-ylsulfonyl)phenyl)thiazol-2-amine (**107**) (500 mg, 1.55 mmol) and *N*,*N*-Diisopropylethylamine (600.2 mg, 4.64 mmol). The mixture was stirred at 55°C for 18 h under N_2_. After cooled, the mixture was concentrated to give the crude product which was purified by prep-HPLC to afford the compound (*S*)-2-methyl-*N*-(4-(4-(piperidin-1-ylsulfonyl)phenyl)thiazol-2-yl)butanamide (40.4 mg, 6.4%) as a white solid HPLC/UV purity > 98% (214.254 nm); LC-MS (ESI): 408.2 (M + H)^+^. ^1^H NMR (400 MHz, CHLOROFORM-*D*) δ 8.00 (d, *J* = 8.5 Hz, 2H), 7.91 (d, *J* = 8.3 Hz, 2H), 7.31 (s, 1H), 3.43 – 3.36 (m, 4H), 3.00 (s, 1H), 2.80 – 2.70 (m, 4H), 2.66 (s, 3H), 1.91 (dt, *J* = 14.3, 7.2 Hz, 1H), 1.78 – 1.66 (m, 1H), 1.38 (d, *J* = 6.9 Hz, 3H), 1.01 (t, *J* = 7.4 Hz, 3H). ^13^C NMR (101 MHz, CHLOROFORM-*D*) δ 175.27, 161.92, 141.01, 137.83, 132.43, 128.80, 126.57, 109.15, 46.98, 42.95, 40.98, 26.65, 25.15, 23.44, 16.43, 11.52. HRMS (ESI-TOF) calcd for C_19_H_25_N_3_O_3_S_2_ [M + H]^+^, 408.1337; found 422.1408.

### FASN-TE Protein Expression and Purification

For biochemical and PTS studies, FASN-TE was expressed recombinantly, incorporating a His_8_ tag at the N-terminus using boundary conditions of (2204-2511). Transformed host strain BL21(DE3) *E. coli* was cultured in Terrific Broth (TB) medium at 37°C until an OD_600_ of ∼0.7 was reached. Protein expression was induced with addition of isopropyl β-D-1thiogalactopyranoside (IPTG) to a final concentration of 0.7 mM, after which cells were grown overnight at room temperature. Cells were collected by centrifugation and lysed by sonication and recombinant FASN-TE was purified using Ni^2+^-affinity chromatography followed by size exclusion chromatography. The eluted proteins were buffer exchanged into 25 mM Tris (pH 8.0), 150 mM NaCl, 0.5 mM TCEP, and 10% glycerol.

### Biochemical FASN-TE Inhibition Assay

Titrated compounds in DMSO (200 nL) were transferred to Greiner 784076 plates by an Echo Liquid Handler (Labcyte, San Jose, CA). 10 µL of buffered 1.8 µM recombinant TE domain of FASN was added followed by 10 µL of 7.0 µM O-methyl fluorescein heptanoate (OMFH) in 10% DMSO as previously described ^59^. Fluorescence, developed by the cleavage of OMFH by the TE domain of FASN, was measured every 5 minutes for 1 hour. Rates of cleavage were calculated from the initial slope data and plotted against the compound dose response to calculate IC_50_ values.

### Differential Scanning Fluorimetry/Protein Thermal Shift Assay

Temperature induced melting of the FASN-TE was carried out at 12 µM in 24 µM Tris (pH 7.4), 24 µM NaCl, 4.8% glycerol, 4.8x Sypro Orange (Invitrogen, Waltham, MA #S6651) 4% DMSO. Melt curves were obtained in individual wells using a MicroAmp® 384-well real-time PCR plate (Applied Biosystems, Waltham, MA #4483285) on a ViiA 7 Real-Time PCR instrument (Applied Biosystems, Waltham, MA) and analyzed using Protein Thermal Shift software v1.2 (Thermo Fisher, Waltham, MA) as previously described ^77^. Compounds were probed at 200 µM in quadruplicate and the change in melting temperature (PTS) value was quantified by subtraction of the reference value for DMSO.

### InCell Pulse Cellular Target Engagement Assay

The optimized miniaturized form of the InCell Pulse target engagement assay was adapted from detailed protocols described recently ^66, 78^. HEK293 cells were grown in Dulbecco’s modified Eagle’s growth medium (GlutaMAX, 10% FBS, antibiotic/antimycotic, 20 µM HEPES, sodium pyruvate) and transiently transfected to express FASN-TE(2204-2511) with a C-terminus ePL-tag using jetPRIME buffer and reagent (Polyplus-transfection, Strasbourg, France #101000046). Analyte compound (75 nL) was transferred to an armadillo PCR plate (Thermo Scientific, Waltham, MA #AB3384) by an Echo Liquid Handler (Labcyte, San Jose, CA). Cells were detached via trypsinization 24-h post-transfection and washed in growth media before plating 7.5 µL of 1.25 x 10^5^ cells/mL onto the plate. After a 1-hour incubation at 37°C, 5% CO_2_ the cells were pulsed for 3 minutes at a specified temperature on a thermocycler (X50h Mastercycler, Eppendorf, Hamburg, Germany). For thermal profile experiments a gradient from 40-65°C was used. For lysis and visualization, 7.5 µL of enzyme fragment complementation lysis and detection reagent (DiscoverX, Luxembourg) was added formulated as previously described ^66^. After a 30-minute incubation in the dark at room temperature, chemiluminescence was measured using a TECAN Spark multimode microplate reader (Tecan; Mannedorf, Switzerland).

### Isothermal CETSA Panel

Measurement of single concentration isothermal cellular target engagement was adapted from the miniaturized protocol described previously ^66^. Transiently transfected cells were prepared as described above. Assay plates were prepared using an Echo Liquid Handler with transfer of 75 nL of 10 mM inhibitor compound in DMSO to the plate. Detached cells (7.5 µL of 1.25 x 10^5^ cells/mL) were applied to the plate, incubated with compound for 1 h, and were subjected to an isothermal heat pulse at 54°C. Assay plates were provided with chemiluminescence InCell imaging reagent and compound chemiluminescence measured as described above. For dose response experiments, 75 nL of compounds diluted by serial dilution were transferred into the assay plate and the assay was performed with an isothermal heat pulse as described above.

### FASN-TE Activity Probe Competition Assay

Pellets of SK-MEl5 cells were resuspended in phosphate buffered saline (PBS) and were lysed by sonication. The cellular debris was pelleted by centrifugation and the soluble lysate obtained for experiments. Lysate was diluted to 1 mg/ mL of total protein as determined by bicinchoninic (BCA) analysis. Assays were performed with 50 µL of 1 mg/mL SK-Mel5 cell lysate with the addition of 200 µM compound for 30 minutes before adding 1 µL of 100 µM TAMRA-FP (ActiveX) Serine Hydrolase probe (Thermo Scientific, Waltham, MA #88318) for a final concentration of 2 µM as previously described ^79^. Activity of the probe was stopped with the addition of SDS buffer and heating at 72°C for 10 minutes. The samples were resolved by SDS-PAGE and either subjected to immunostaining using anti-FASN antibodies (Cell Signaling Technologies, #C20G5) or analyzed by the fluorescence of the Tamra-FP probe. Quantification of labeled proteins utilized Image Lab Software (*Bio-Rad Laboratories*, Hercules CA). Gels were quantified by measuring the intensity of each band and normalized to the total protein loaded in the lane, as visualized by Coomassie staining. Experiments with recombinant FASN-TE were performed in a similar manner as described above with the assay at 50 µL of 100 µg/mL recombinant TE domain of FASN-TE.

### Palmitate Synthesis Inhibition

SK-Mel5 cells were grown in RPMI 1640 growth medium (10% FBS, antibiotic/antimycotic), detached with trypsin and washed with PBS. The cells were then resuspended in Minimum Essential Media (10% FBS, antibiotic/antimycotic, GlutaMAX, non-essential amino acids) with 1 g/L U-^13^C_6_ (Cambridge Isotope Laboratories Inc., Tewksbury, MA #CLM-139-PK) and plated out 5x10^5^ cells in 2 mL media with 6 µL of titrated compound in DMSO in a 6-well plate. After 72 h of incubating at 37°C with 5% CO_2_, the media was aspirated, and cells were washed with PBS. 450 µL of 50% methanol was added to cells and the cells were frozen on dry ice for 30 minutes and then were thawed on wet ice for 10 minutes before being manually scraped and collected. Chloroform was added to these cell extracts to separate the fatty acids before they are derivatized and separated by GC-MS where ^13^C-labelled palmitate was isolated to measure *de novo* synthesis levels.

### Microsomal Stability in Vitro Assay

Pooled mouse (XenoTech, LLC, M1000), rat (XenoTech, LLC, R1000), or human (XenoTech, LLC, H0610) liver microsomes were preincubated with test compounds at 37 °C for 5 min. The reaction was initiated by addition of NADPH with final assay concentrations of 4 μM test compound, 2 mM NADPH, and 1 mg/mL (total protein) liver microsomes. Zero timepoint samples (100 µL) were taken immediately and quenched with 100 µL of 5:1 ACN/H2O containing internal standard. Compounds were incubated with microsomes and NADPH for one hour at 37 °C with shaking. Endpoint samples (100 µL) were mixed with 100 μL of 5:1 ACN/H2O containing internal standard. Samples were centrifuged at approximately 13 000 rpm for 10 min. Supernatants were removed and transferred into a deep 96-well plate and the amount of test compound was quantified using a Shimadzu Nexera X2 High-performance liquid chromatography (HPLC) (Shimadzu Scientific Instruments; Kyoto, Japan) coupled to an AB Sciex 6500 QTRAP (AB Sciex LLC; Toronto, Canada). The change of the AUC (area under the curve) of the test compound as a function of time was used as a measure of microsomal stability. Compounds were run in duplicate with a positive control. Negative controls were run with no NADPH addition.

### Plasma Stability in Vitro Assay

A 20 μL aliquot of a 10 mM solution in DMSO of the test compound was added to 2.0 mL of heparinized mouse plasma (Bioreclamation IVT, MSEPLNAHP-C57-M), rat plasma (Bioreclamation IVT, RATPLNAHP-M), or human plasma (Bioreclamation IVT, HUMANPLNHYNN) to obtain a 100 μM final solution. The mixture was incubated for 1 h at 37°C while shaking. Aliquots of 100 μL were taken at 15 min intervals and diluted with 100 μL of 5:1 ACN/H2O containing internal standard. After mixing, samples were centrifuged at approximately 13 000 rpm for 10 min. The supernatants were transferred into a deep 96-well plate, and the amount of test compound was quantified using a Shimadzu Nexera X2 High-performance liquid chromatography (HPLC) (Shimadzu Scientific Instruments; Kyoto, Japan) coupled to an AB Sciex 6500 QTRAP (AB Sciex LLC; Toronto, Canada). The change of the AUC of the test compound as a function of time was used as a measure of plasma stability. Test compounds were run in duplicate with positive and negative controls.

### In vivo studies

All experimental protocols were approved by the Institutional Animal Care and Use Committee (IACUC) of Sanford Burnham Prebys Medical Discovery Institute and performed in conformity with regulations and guidelines of the National Institutes of Health (NIH) and the Association for Assessment and Accreditation of Laboratory Animal Care (AAALAC). The Institute’s Animal Care and Use Program is accredited by AAALAC International and a Multiple Project Assurance A3053-1 is on file in the Office for Protection from Research Risks (OPPR) a division of the U.S. Department of Health and Human Services (DHHS). Adult female C57BL/6J mice were purchased from Jackson Laboratories and were housed with free access to food and water on a 12 h light/dark cycle. For snapshot PK studies, compounds were formulated in 5% DMSO, 10% Tween-80, and 85% H_2_O and dosed orally into a cohort of 3 mice at 10 mg/kg (1 mg/mL, 10 mL/kg). Blood samples were collected retro-orbitally at the indicated times and plasma was prepared by centrifugation. For full PK studies, **78S** was formulated in 5% DMSO + 5% Kolliphor HS15 + 90% saline at 0.2 mg/mL for the IV arm of the study and were dosed at 1 mg/kg (5 mL/kg) via tail vein injection (n=3 mice). For the PO arm of the full PK study, **78S** was formulated in 5% DMSO + 10% Tween 80 + 85% water at 1 mg/mL and was dosed at 10 mg/kg (10 mL/kg) via oral gavage (n=3 mice). Approximately 30 µL blood/time point was collected into K_2_EDTA tube via the facial vein for bleeding. Plasma samples were placed on ice and centrifuged to obtain plasma sample 3,000 × g, 5 min under 4℃) within 15 minutes of collection. For quantification of FASN-TE inhibitor compounds, plasma samples were extracted with acetonitrile:water 4:1 with 0.05% formic acid. This solution contained verapamil as an internal standard. Samples were clarified by centrifugation, and the supernatants were diluted with acetonitrile:water and analyzed by LC-MS/MS using a Shimadzu Nexera X2 High-performance liquid chromatography (HPLC) (Shimadzu Scientific Instruments; Kyoto, Japan) coupled to an AB Sciex 6500 QTRAP (AB Sciex LLC; Toronto, Canada). PK analysis was performed by noncompartmental analysis utilizing the Microsoft Excel plugin PKSolver 2.0.

## ANCILLARY INFORMATION

### Accession Codes

The atomic coordinates and structure factors of the FASN-TE domain in the apo form and bound to inhibitors 66 and 71 have been deposited in the Protein Data Bank (PDB) under accession numbers 9ZLX, 7MHD and 7MHE respectively. Authors will release the atomic coordinates upon article publication.

### Supporting Information

Supporting information is available in the file FASN-TE-SI.pdf and is provided free of charge. **Figure S1**, an overview of the role of FASN in cancer. **Figure S2**, a figure showing the structures of selected FASN-TE inhibitors. **Supplemental protocol** methods for the molecular cloning of the expression construct for X-ray studies. **Supplemental protocol** methods for the expression and purification of FASN-TE (2204-2511), methods for the crystallization of FASN-TE and structure determination. **Table S1**, a table providing the data collection and refinement statistics (molecular replacement) for the x-ray crystal structure. **Figure S3**, a correlation plot showing the trend of stereopure arylthiazole compound biochemical potency compared to PTS stabilization of the FASN-TE domain **Figure S4**, a figure showing a structural comparison of arylthiazole FASN-TE inhibitor conformation to bound lipid inhibitors. **Figure S5**, a figure showing chemical shift perturbation resulting from FASN-TE inhibitor binding to FASN-TE. **Figure S6**, a figure demonstrating that FASN reacts with the serine hydrolase activity probe TAMRA-FP. **Table S2**, a table showing off-target activity against a panel of proteases for our FASN-TE inhibitor series. **Figure S7**, a figure showing that stereopure arylthiazoles have reduced off-target inhibitory activity against serine hydrolases in cell lysates compared to FASN. **Figure S8**, a figure showing selectivity of arylthiazoles for FASN compared to ACOT-4. **Scheme S1**, a scheme showing the general procedure for synthesis of arylthiazoles described in Table 1. **Scheme S2**, a scheme showing the chemical synthesis of intermediates for stereopure S- R- aminiothiazole FASN-TE inhibitors. **Chemistry Experimental**, additional chemistry experimental methods, chromatograms showing the HPLC and chiral purity of compounds used for in vivo studies and references specific to the supporting information. We have provided a molecular string file JMC_FASN_A1.CSV containing a list of all the compounds studied in this manuscript. All other data supporting the findings of this study are available from the corresponding author upon reasonable request.

## AUTHOR INFORMATION

### Corresponding Author

* **Nicholas D. P. Cosford** – Center for Therapeutics Discovery, Cancer Metabolism and Microenvironment Program, Sanford Burnham Prebys Medical Discovery Institute, 10901 North Torrey Pines Road, La Jolla, California 92037, USA;

### Present Addresses

LJSDB is currently an employee of Chemium, Ottignies-Louvain-la-Neuve, Belgium. MEM is currently an employee of Maravai LifeSciences, San Diego, CA, United States. PG is currently an employee of Bio-Techne, San Marcos, CA, United States. CDW is currently an employee of Thermo Fisher Scientific, San Diego, CA, United States. PT is currently an employee of IDEAYA Biosciences, San Francisco, CA, United States. MPH is currently an employee of Vividion Therapeutics. Inc. San Diego, CA, United States.

### Author Contributions

The manuscript was written through contributions of all authors. All authors have given approval to the final version of the manuscript.

## ACKNOWLEDGMENTS

We would like to thank Jessica De Ingeniis, Martina Felderman and Darren Finlay for helpful advice about experimental design, Robert Liddington for advice and assistance with X-ray crystallography, and Bobby R. Lucero and Kokila Shankar for their assistance in supporting production of data needed for the resubmission of the manuscript. We are grateful to Jilei Duan, Jing Dong, Olga Zagnitko (Viva Biotech Ltd.) for synthetic chemistry support.

## ABBREVIATIONS

BCA: bicinchoninic acid
CETSA: cellular thermal shift assay
DMSO: dimethyl sulfoxide
EC50: half- maximal effective concentration
ePL: enhanced ProLabel®
FASN: fatty acid synthase
FBS: fetal bovine serum
GC-MS: gas chromatography–mass spectrometry
HEK293: human embryonic kidney 293 cells
HEPES: 4-(2-hydroxyethyl)-1-piperazineethanesulfonic acid
HPLC: high-performance liquid chromatography
HSQC: heteronuclear single quantum coherence
HTS: high-throughput screen
IC50: half-maximal inhibitory concentration
IPTG: isopropyl β-D-1-thiogalactopyranoside
LC-MS: liquid chromatography–mass spectrometry
MGLP: methyl γ-linolenylphosphonate
MLPCN: Molecular Libraries Probe Production Centers Network
MLSMR: Molecular Libraries Small Molecule Repository
NADPH: nicotinamide adenine dinucleotide phosphate
NMR: nuclear magnetic resonance
OMFH: *O*- methyl fluorescein heptanoate
PTS: protein thermal shift
SAR: structure–activity relationship
SDS- PAGE: sodium dodecyl sulfate–polyacrylamide gel electrophoresis
SEM: standard error of the mean
TB: terrific broth
TCEP: tris(2-carboxyethyl)phosphine
TE: thioesterase.

