## Supplemental Data for "Reversible, Orally Bioavailable Small-Molecule Inhibitors Occupy the Substrate Tunnel of the Thioesterase Domain of Fatty Acid Synthase"

*AUTHOR ADDRESS*

***Corresponding Author**

*Supporting Information*

### Table of Contents

1. SI Figure 1: Overview: Fatty Acid Synthase (FASN) in cancer…………………………... S3
2. SI Figure 2: Selected FASN-TE inhibitors……………………………………………..S4
3. Molecular cloning, expression, purification of FASN-TE for biophysical studies…………S5
4. Crystallization conditions for co-crystal structures of FASN-TE…………………………...S6
5. SI Table 1: Data collection and refinement statistics (molecular replacement)……………..S7
6. SI Figure 3: Correlation plot: biochemical potency and protein thermal shift………………S8
7. SI Figure 4: Comparative conformation of arylthiazole inhibitors…………………………S9
8. SI Figure 5: Chemical shift perturbation of the FASN-TE indicates that the binding of **66** and **50** to the FASN-TE utilize common determinants……………………………………………..S10
9. SI Figure 6: FASN reacts with serine hydrolase activity probe TAMRA-FP………………..S11
10. SI Table 2: Off-target protease activity of **50**………………………………………………..S12
11. SI Figure 7: Stereopure arylthiazoles have reduced off-target inhibitory activity against serine hydrolases in cell lysates compared to FASN……………………………………………….S13
12. SI Figure 8: Arylthiazoles have minimal off target activity for ACOT-4 thioesterase……..S14
13. Chemistry experimental…………………………………………………………S15–S35
14. SI Scheme 1. General procedure for synthesis of arylthiazoles described in Table 1………..S15
15. SI Scheme 2: Chemical synthesis of intermediates for stereopure S- R-aminiothiazole FASN-TE inhibitors………………………………………………………………………..S27
16. HPLC purity of compounds used for in vivo studies…………………………………...S36-S42
17. Stereochemistry of **50**, **78R** and **78S** by chiral HPLC……………………………………….S43
18. References……………………………………………………………………..S44-S46

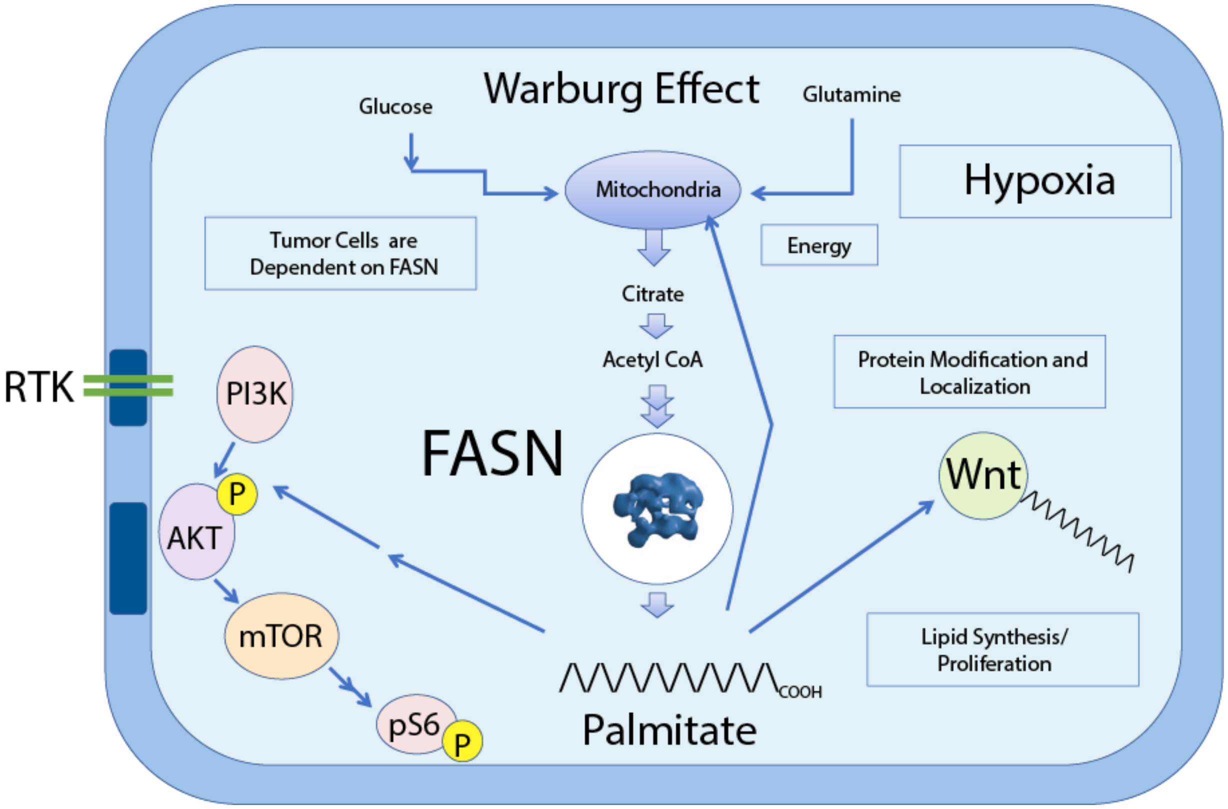

**SI Figure 1. The Role of Fatty Acid Synthase (FASN) in Cancer:** First recognized as the prognostic biomarker OA-519 in breast cancer in 1994, increased FASN catalytic activity in cancer cells correlates with higher malignancy and poor prognosis [1]. As the core enzyme capable of de novo fatty acid synthesis, a number of mechanistic consequences of upregulated fatty acid synthesis have been identified as potentially contributing to this linkage. Of primary importance is the long-recognized dependence of tumor cells on aerobic glycolysis, cf. the Warburg effect. This shift to aerobic glycolysis is necessary both for increased energy but also provides the lipid precursors for cell division. However the increased fatty acid pool perhaps has other growth promoting effects from major signaling pathways such as Wnt6/β-catenin [2]. Perturbations in tumor cell membrane composition, expansion of lipid raft microdomains and membrane fluidity that promote receptor tyrosine kinases (RTK) have also been described to be the result of increased FASN synthesis and is sensitive to FASN inhibition [3-6].

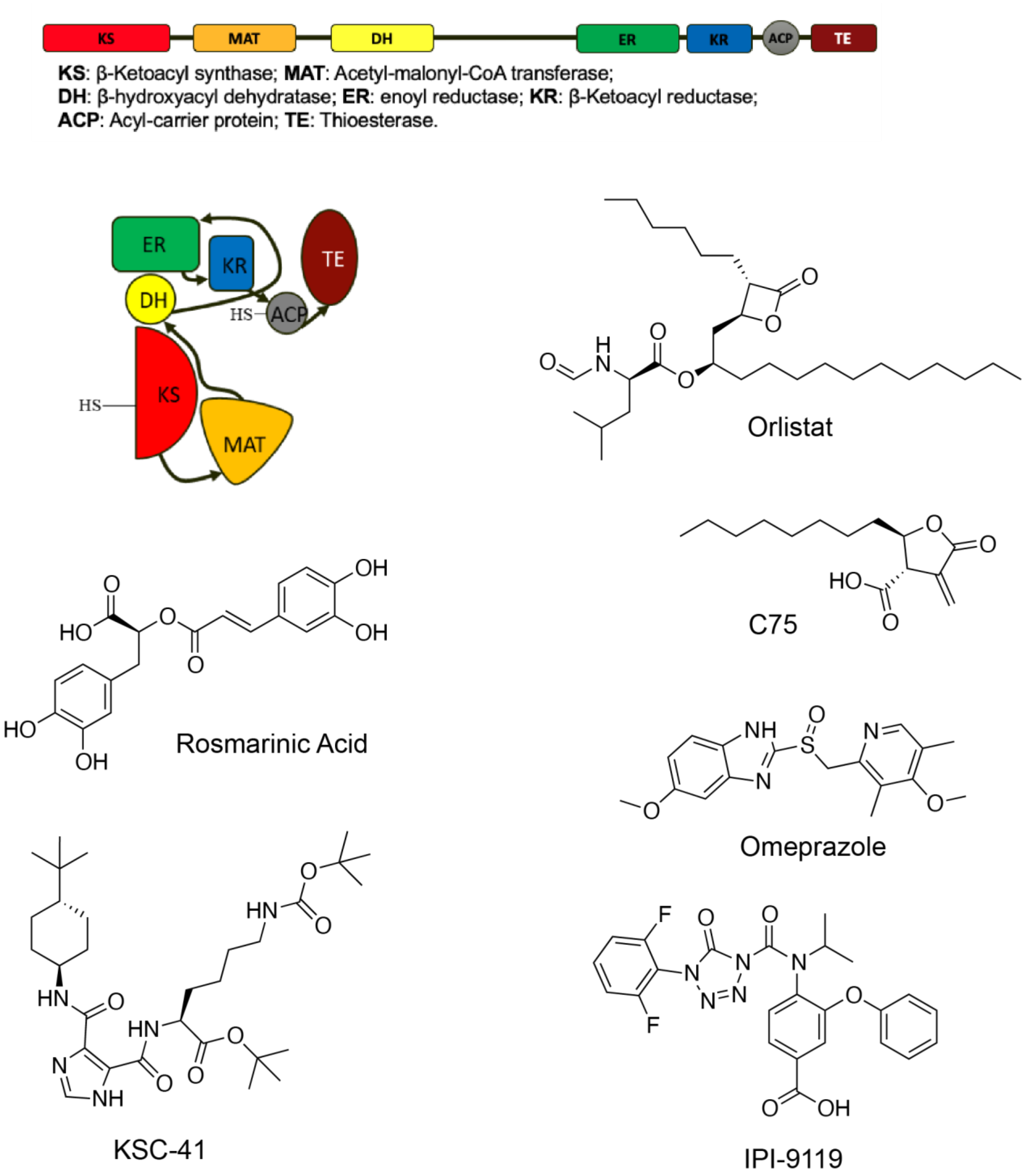

**SI Figure 2. Selected FASN-TE inhibitors.** An essential part of the FASN polyprotein, the thioesterase domain of FASN is recognized as a target for inhibitory compounds. A select number of known FASN-TE inhibitors and their structures are shown. Orlistat [7], cerulenin C75 [1], Rosmarinic Acid [8], Omeprazole [9], KSC-41 [10], and IPI-9119 [11], have been described to inhibit the FASN-TE domain and have been investigated for efficacy in cancer.

**Molecular Cloning of the Expression Construct for X-ray Studies**

The recombinant plasmid of FASN-TE pNHisFAS (2204-2511) encodes a form of the fatty acid synthase thioesterase domain that includes N- and C- terminal disordered segments when compared to that co-crystallized by Zhang et al. [12]

**Expression and Purification of FASN-TE (2204-2511)**

For the ligand-bound crystal structure trial recombinant expression of FASN-TE (2204-2511) was achieved by transformation into BL21(DE3) with growth in LB media/Kanamycin to mid log-phase and induction overnight (0.7 mM IPTG) at 16°C. Homogenization with Ni column binding buffer (30 mM Tris pH 8.0, 300 mM NaCl, 30 mM Imidazole, 10% glycerol) was followed by lysis of cells using treatment with lysozyme (0.1 mg/mL) and sonication. For a 2 L growth the resuspended volume was 250 mL. Clarification by centrifugation (15,000 X g) was performed to produce a soluble protein rich lysate. This solution was applied to Ni-sepharose (GE Healthcare) (40 mL resin volume) by gravity. The resin was transferred to a Tricorn column for purification by FPLC. The column was washed with 5 volumes of Ni column binding buffer (1.0 mL/min) and was then subjected to the development of a 120 mL imidazole gradient 30-300 mM. Peak fractions were supplemented with TCEP to 10 mM. The Ni column rich fractions were pooled (100 mL, approximately 1 mg/mL) and dialyzed against a thrombin cleavage buffer (20 mM Tris pH 7.8, 150 mM NaCl, 1 mM $CaCl_2$, 10 mM DTT) at 4°C using 1 μg/mg bovine thrombin. The protein was concentrated by stirred cell ultrafiltration to 5 mg/mL (MWCO 10,000) and was then subjected to purification by gel filtration chromatography. The Superdex S200 16/60 column (GE Healthcare) purification employed a flow buffer of 50 mM Tris pH 7.8, 50 mM NaCl at 16°C. The FASN-TE eluted at a column volume of 90 mL. Peak fractions were pooled, adjusted to 10 mM TCEP and were concentrated by ultrafiltration (MW 10,000) (Millipore, Billerica, MA) to 10-15 mg/mL. The FASN-TE sample was immediately subjected to crystallization trials. For the structure of the Apo-form of FASN-TE (2204-2511) purification followed the above purification but final purification was achieved with the use of an ion exchange monoQ 4.6/100 column (GE Healthcare) using a NaCl gradient. The protein was eluted into a buffer with 100 mM NaCl, 20 mM Tris, 2 mM DTT and concentrated using MWCO 10 kDa-cutoff concentrators (Millipore, Billerica, MA) and subjected to crystallization trials without freezing.

**Expression and Purification of FASN-TE (2204-2511) for NMR Studies**

For NMR studies the $^{15}NH_4SO_4$ enriched minimal media was employed to express the uniformly labeled FASN-TE (2204-2511) using methods and media as previously described [13]. Purification by Ni-sepharaose afforded the enriched protein FASN-TE (2204-2511) that was dialyzed against 1 X PBS with 2 mM TCEP. Ultrafiltration using a MWCO 10,000 membrane was performed to prepare a protein sample at 0.2 mM. NMR samples were prepared at 10% $D_2O$, 0.6% DMSO and HSQC spectra were recorded using an Avance 600 MHz instrument equipped with a cryoprobe.

**Crystallization of FASN-TE and Structure Determination**

The apo-form of FASN-TE (2204-2511) and its complexes with **66** and **71** were crystallized using the hanging drop vapor diffusion method and conditions identified by the 96-reagent JBScreen Kinase (MiTeGen, Ithaca, NY, USA) and Phoenix drop dispenser (Art Robbins Instruments, Sunnyvale, CA, USA). The apo-protein crystallized using 0.3 µL of 8 mg/ml FASN-TE in 100 mM NaCl, 50 mM BisTris pH 6.0, 10 mM DTT and 0.2 µL of well solution 10% PEG400, 50 mM Tris-Cl pH 8.5, 1.0 mM DTT, 1.0 mM ethylenediaminetetraacetic acid disodium salt (EDTA), 300 mM NaCl. The crystals of the protein-inhibitor complexes were obtained using same conditions, but the protein solution also contained 0.5 mM of an inhibitor and 1% DMSO. The crystals grew in 3-5 days at $20^0$C. The crystals were flash-frozen in liquid nitrogen without addition of cryoprotectant and the data were collected from single cryo-cooled crystals (100 K) at the beamline 9-2 with the wavelength 1.03316 Å and processed using XDS [14] or HKL-2000 [15] and the CCP4i program suite [16]. The structure of FASN-TE with **66, 71** and the apo form was solved using the molecular replacement program PHASER [17] and employed PDB ID 3TJM as the search model. Model building was done with COOT [18] and the refinement by PHENIX [19] and REFMAC 5.8 [20, 21] Figures of the protein structures and alignments were prepared using PyMOL (Molecular Graphics System, Version 1.7.2.1, Schrodinger, LLC) and the Blender Plugin Molecular Nodes (Version 0.12.0., Zenodo). 2D plots made use of LIGPLOT.[22] The data collection and refinement statistics are presented in **SI Table 1**.

**SI Table 1. Data collection and refinement statistics (molecular replacement)**

| | **Apo** | **71** | **66** |
|---|---|---|---|
| **PDB-ID** | **9ZLX** | **7MHD** | **7MHE** |
| **Data collection** | | | |
| Space group | P $2_12_12_1$ | P $2_12_12_1$ | P $2_12_12_1$ |
| Cell dimensions | | | |
| α, β, χ (Å) | 50.95 61.45 76.16 | 53.89 61.49 77.45 | 52.68 56.50 75.64 |
| α, β, γ (°) | 90 90 90 | 90 90 90 | 90 90 90 |
| Resolution (Å) | 76.2-2.6 (2.72-2.6)* | 35.9-2.0 (2.08-2.03) | 45.3-2.8 (2.92-2.77) |
| $R_{merge}$ | 0.165 (0.75)* | 0.049 (0.62) | 0.221 (1.19) |
| $I / \sigma I$ | 5.3 (1.3) | 15.1 (2.1) | 5.2 (1.4) |
| Completeness (%) | 96.2 (93.7) | 98.2 (82.5) | 90.7 (89.2) |
| Redundancy | 3.6 (3.4) | 4.5 (3.1) | 4.6 (4.6) |
| **Refinement** | | | |
| Resolution (Å) | 47.8 - 2.6 | 35.9 - 2.0 | 45.3 - 2.8 |
| $R_{work}$ / $R_{free}$ | 0.196/0.260 | 0.191/0.250 | 0.213/0.280 |
| No. atoms | | | |
| Protein | 2098 | 2261 | 2154 |
| Ligand/ion | - | 27 | 31 |
| Water | 25 | 99 | 16 |
| *B*-factors | | | |
| Protein | 44.4 | 47.2 | 47.3 |
| Ligand/ion | - | 55.9 | 54.2 |
| Water | 44.3 | 46.2 | 45.2 |
| Wilson | 47.0 | 39.2 | 57.8 |
| R.m.s. deviations | | | |
| Bond lengths (Å) | 0.013 | 0.016 | 0.015 |
| Bond angles (°) | 1.63 | 1.98 | 1.72 |
| Ramachandran favor/outliers (%) | 96.2/0.4 | 95.4/0.0 | 96.3/0.0 |

*Values in parentheses are for highest-resolution shell.
Rfree test size = 5%
Apo-structure has the resolution anisotropy (in the shell *I/sdI*=1.5) 3.0x3.1x2.6 Å
FASN-TE/**66** structure has the resolution anisotropy (in the shell *I/sdI*=1.5) 2.0x2.0x2.0 Å
FASN-TE/**71**-structure has the resolution anisotropy (in the shell *I/sdI*=1.5) 3.3x3.0x2.8 Å

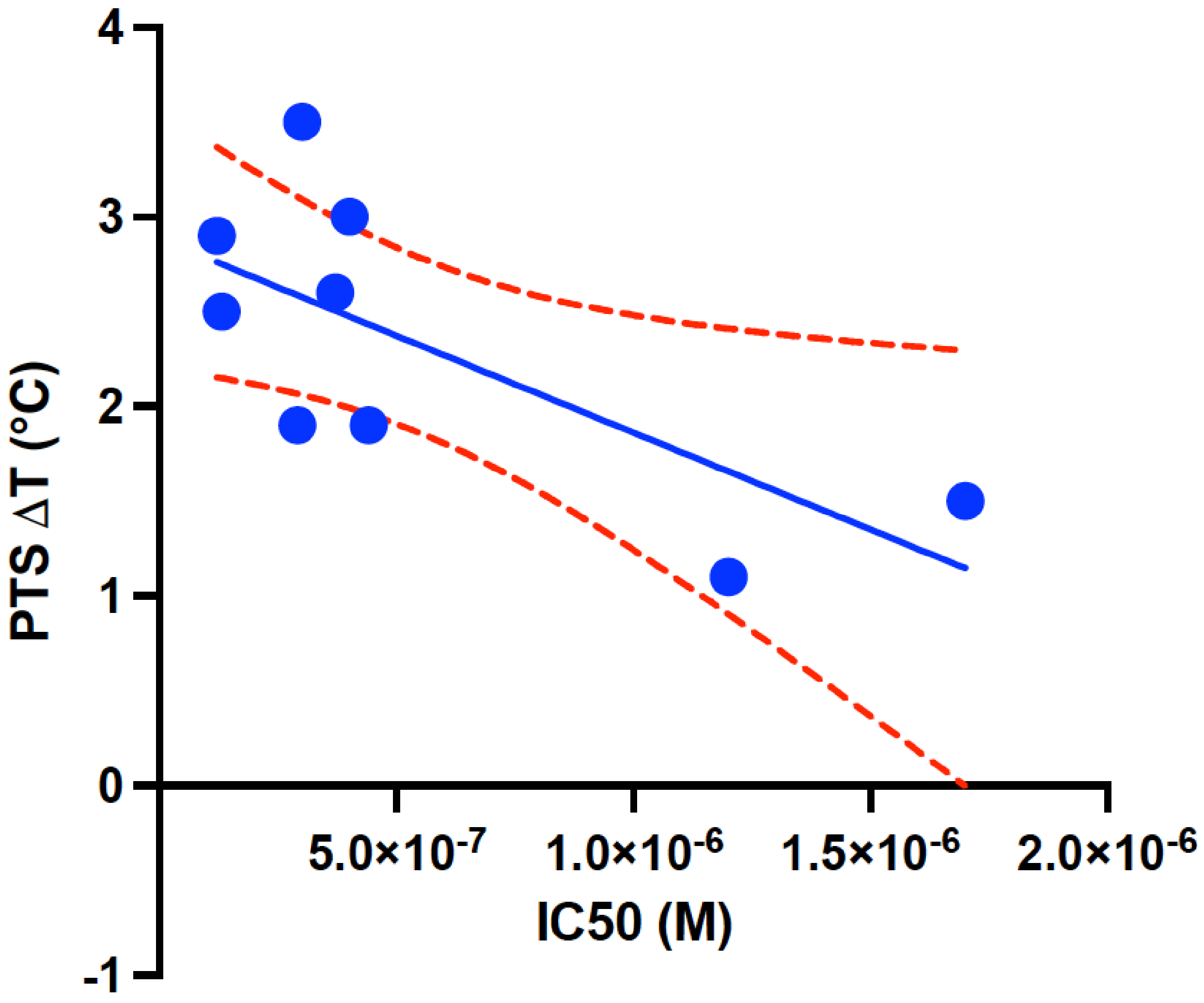

**SI Figure 3. Correlation plot: biochemical potency and protein thermal shift.** A correlation plot showing trend of stereopure arylthiazole compound biochemical $IC_{50}$ potency compared to PTS stabilization of the FASN-TE domain. Indicated is a least-squares fit of the data and a 95% confidence level.

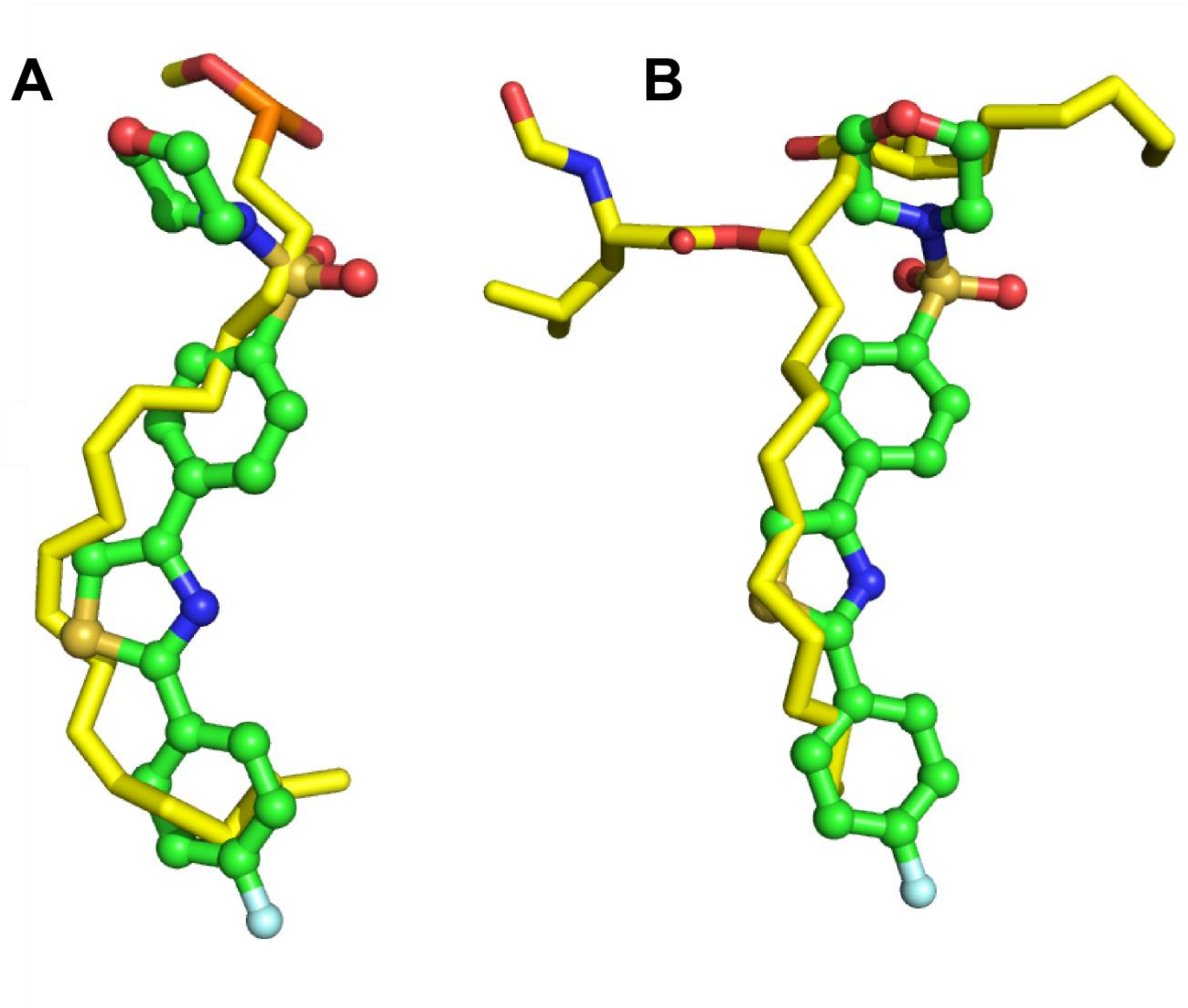

**SI Figure 4. Structural comparison of arylthiazole FASN-TE inhibitor conformation to bound lipid inhibitors.** A comparison of the FASN-TE bound confirmation of **66** and the A) bound state of dihomo-gamma-linolenic acid (3TJM) or B) covalent-Orlistat (2PX6) [12, 23]. The protein structures were superimposed by their backbone resides and the relative position of the bound ligands is shown as a basis for comparison. Both the polyalkyl chain of dihomo-gamma-linolenic acid (yellow) and Orlistat (yellow) follow a similar curved path through the hydrophobic tunnel. Evident is that **66** adopts a similar bent confirmation that would seem to mimic these more lipid-like inhibitors. Differences and similarities between the confirmation of these ligands suggest other possibilities for improved ligand spatial occupancy and optimization. The FASN-TE gatekeeper tunnel possesses an inherent capacity to accommodate flexible bound states.

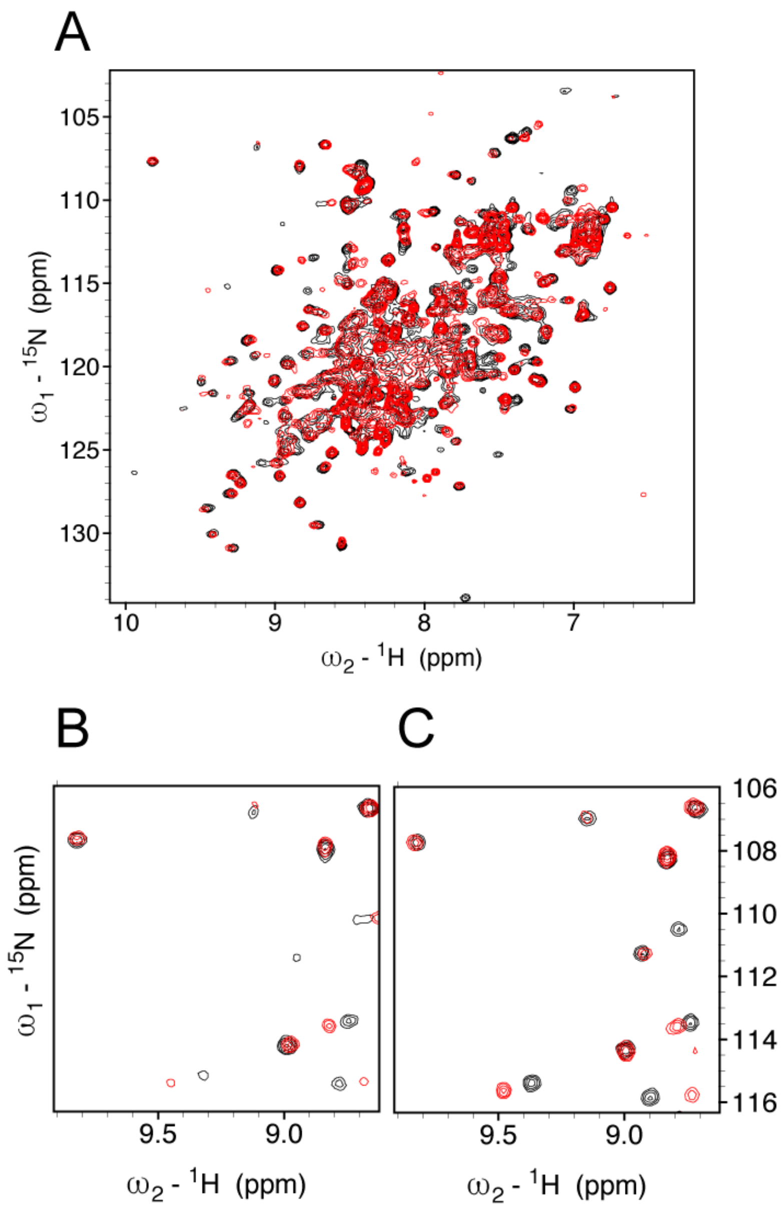

**SI Figure 5. Chemical shift perturbation of the FASN-TE indicates that the binding of 66 and 50 to the FASN-TE utilize common determinants.** A) The HSQC spectrum of $^{15}N$ labeled FASN-TE (2204-2511) (0.15 mM) in PBS buffer/0.6% DMSO recorded at 600 MHz (Black). Superimposed (Red) is the spectrum upon addition of 0.3 mM 50. B) Detailed HSQC spectral region of perturbed crosspeaks for 50. Indicated in black are crosspeaks arising from the unbound 15N labeled FASN-TE. Indicated in red are residues changed upon the addition of 50 at 0.3 mM. C) Detailed HSQC spectral region indicating perturbed crosspeaks for **66** (red) on FASN-TE (black). Shared perturbed residues indicate a common binding mode for these two related compounds.

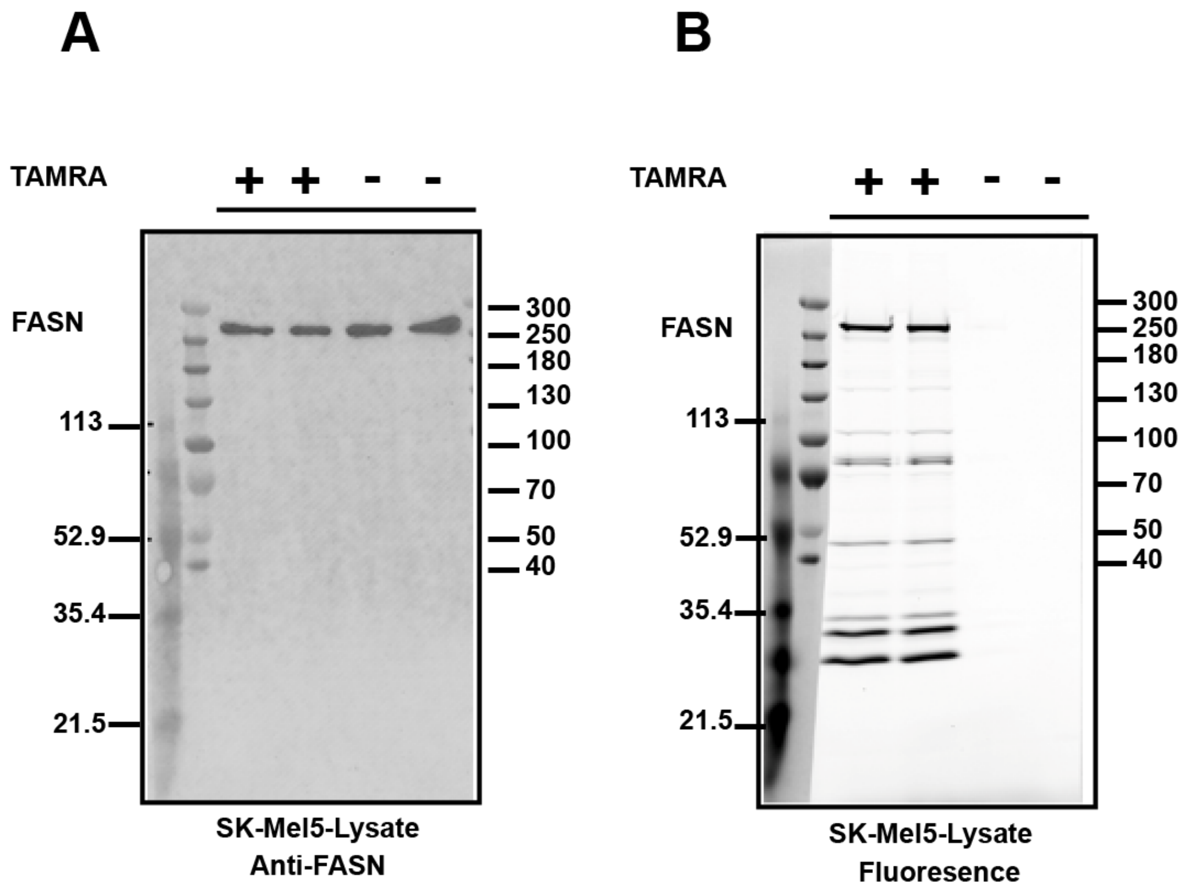

**SI Figure 6. FASN reacts with serine hydrolase activity probe TAMRA-FP.** A) Immunoblot probing of SK-Mel5 lysates with an Anti-FASN antibody confirms the molecular weight and identity of FASN in the cellular lysate. B) Fluorescent imaging of FASN indicates that it robustly reacts with the TAMRA-FP activity-based probe. Serine hydrolase proteins at 81.5, 29, and 26 kDa yield strong signals to the TAMRA-FP and can serve as a benchmark for selective inhibition by arylthiazole inhibitors.

**SI Table 2. Off-target protease activity of 50.** Inhibition studies performed at Reaction Biology (Malvern, PA).

| Target | Compound $IC_{50}$ (nM) | | Control compound ID |
|---|---|---|---|
| | **50** | **Control** | |
| ACE 1 | Inactive | 685 | Captopril |
| ADAM 10 | Inactive | 217 | GM6001 |
| BACE1 | Inactive | 171 | B-Secretase inhibitor IV |
| Calpain 1 | Inactive | 69 | E 64 |
| Caspase 1 | Inactive | 89 | IETD-CHO |
| Cathepsin G | >10000 | 1083 | Chymostatin |
| Chymotrypsin | 1720 | <1 | Chymostatin |
| DDP-IV | Inactive | 225 | P32/98 |
| Granzyme B | Inactive | 1670 | IETD-CHO |
| HCV NS3/4A | Inactive | 1682 | HCV inhibitor II |
| HIV 1 | Inactive | 25 | Pepstatin A |
| MMP 1 | Inactive | 1 | GM6001 |
| TACE/ADAM17 | Inactive | 3 | GM6001 |
| Tryptase beta II | Inactive | 3 | Gabexate Mesylate |

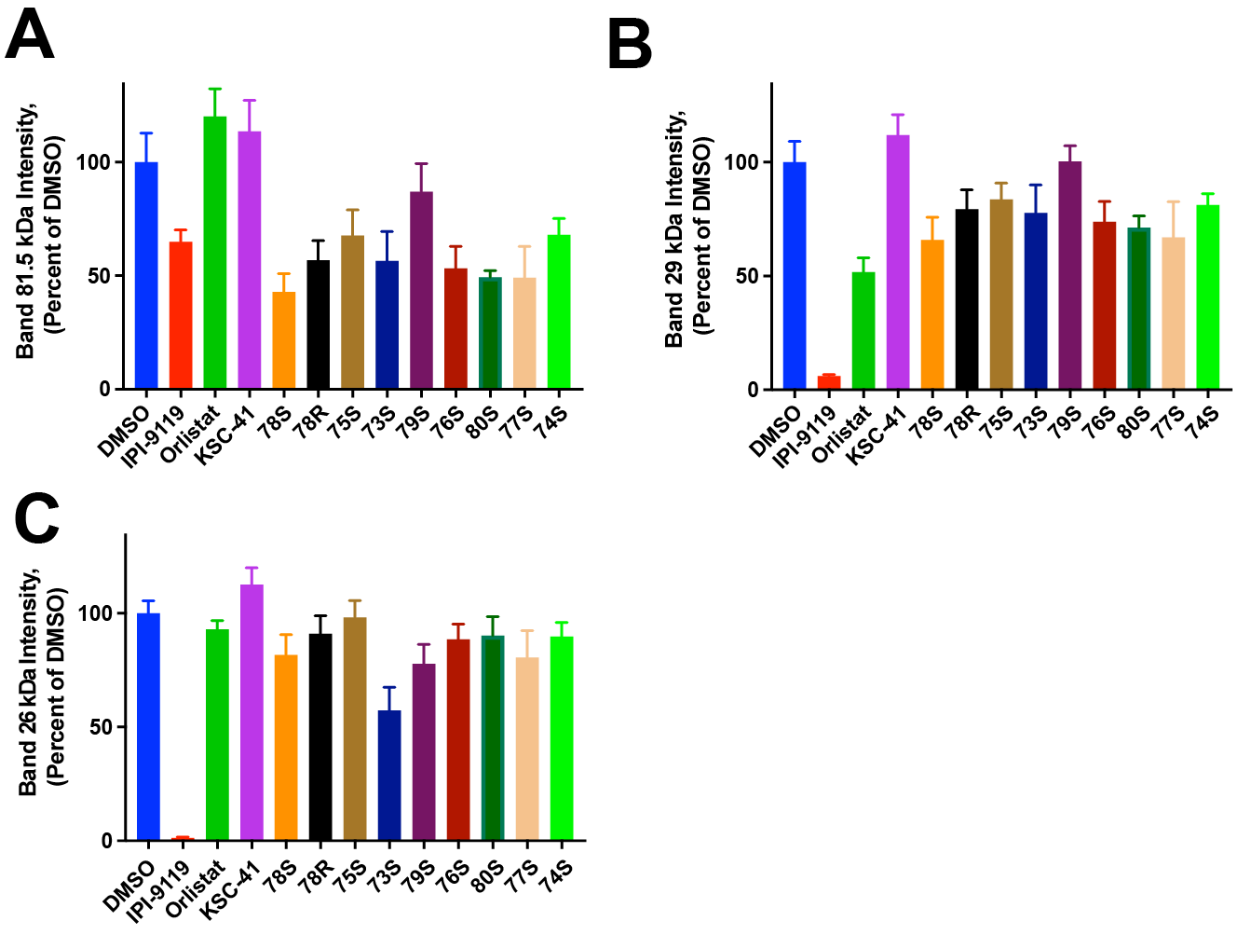

**SI Figure 7. Stereopure arylthiazoles have reduced off-target inhibitory activity against serine hydrolases in cell lysates compared to FASN.** Quantification of the fluorescent signal intensity of the TAMRA-FP active site probe in Figure 3. A) 81.5 kDa, B) 29 kDa, and C) 26 kDa bands in SK-Mel5 lysate treated with a panel of potent arylthiazole compounds relative to DMSO mean±SEM (n=3, duplicate).

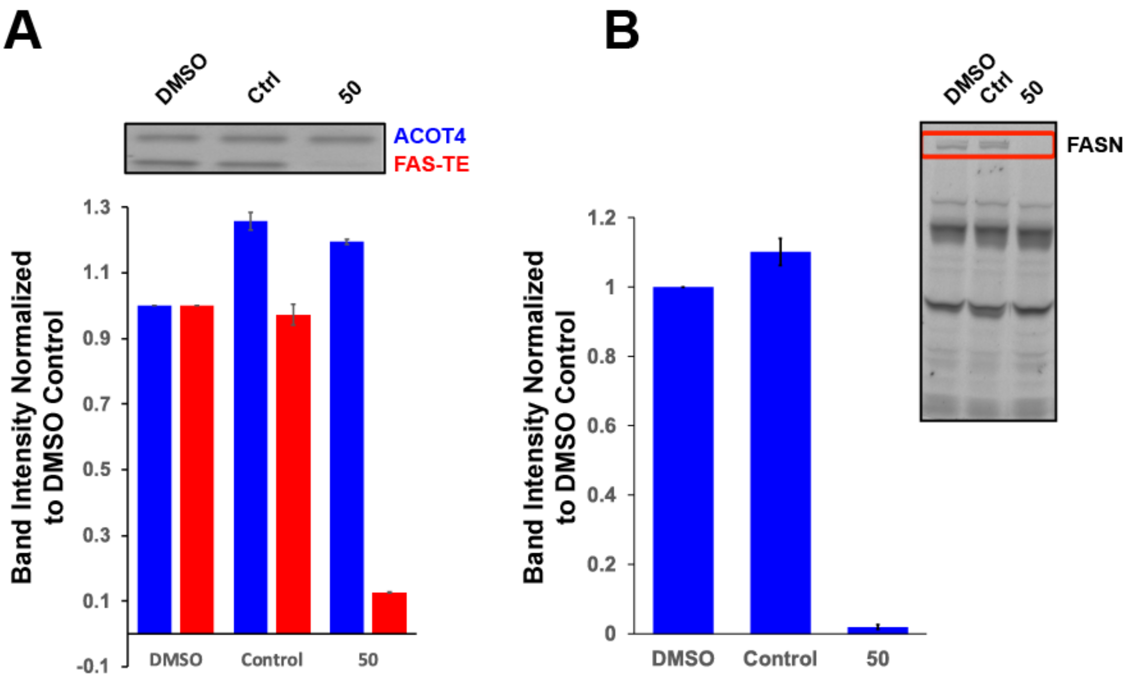

**SI Figure 8. Arylthiazoles have minimal off target activity for members of the thioesterase family.** The selectivity of FASN-TE inhibitors in an activity-based probe assay used methods previously described.[24] A) In an in vitro assay, recombinant acyl-coenzyme A thioesterase 4 (ACOT-4) and the FAS-TE were combined and incubated with the activity-based serine hydrolase probe fluorophosphonate-polyethylene glycol-6-carboxytetramethylrhodamine (FP-PEG-TAMRA). After the labeling reaction, the results are resolved on an SDS PAGE gel. Coincubation with compound **50** selectively inhibits FP-PEG-TAMRA binding to purified FAS-TE but not ACOT-4. An inactive control compound (4-methyl-*N*-(4-(4-(morpholinosulfonyl)phenyl)thiazol-2-yl)benzamide and DMSO vehicle do not inhibit labeling reaction with the FP-PEG-TAMRA. B) In the context of a cellular whole cell lysate prepared from PC-3 cells similar results are obtained. Arylthiazole compound **50** is able to diminish interaction of the full length FASN by the activity-based probe whereas DMSO and the control compound do not attenuate labeling. Other thioesterases in the PC-3 cell lysate, highlighted by the activity-based probe, show no observable change in band intensity.

### Chemistry Experimental

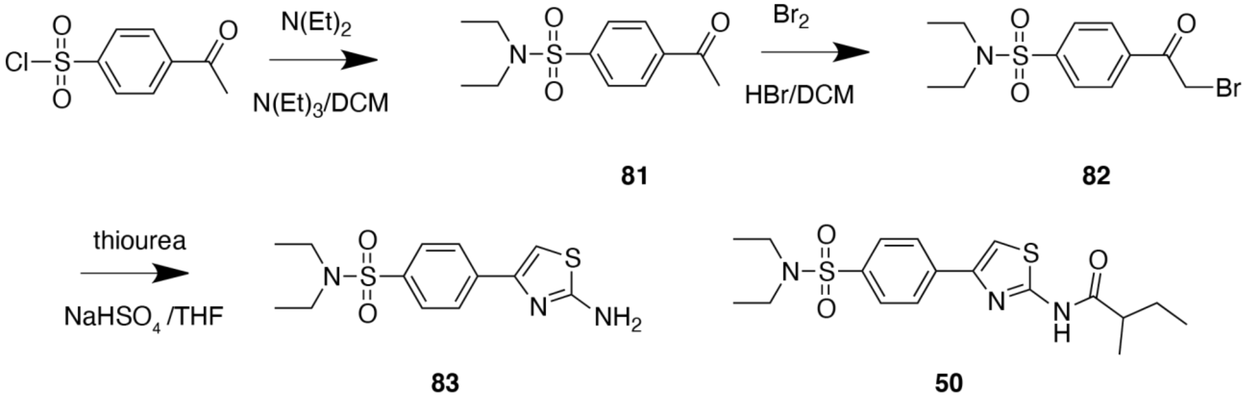

**SI Scheme 1. General Procedure for synthesis of arylthiazoles described in Table 1.** Details on the synthesis of **50** are provided in the scheme above and in the main text. Compounds **1-72** were prepared using analogous reagents and methods.

**4-Acetyl-N,N-diethyl-benzenesulfonamide (81).** A solution of 4-Acetylbenzenesulfonyl chloride (3.0 g, 13.7 mmol) was dissolved in dry DCM (150 mL) and combined with triethylamine (1.9 g, 18.7 mmol) and diethylamine (0.91g, 12.5 mmol). The reaction was stirred at ambient temperature for 12 h. The reaction was quenched with $Na_2CO_3$, washed with brine and dried over $Na_2SO_4$ before concentration to yield the crude reagent (3.3 g 12.5 mmol) which was used without further purification.

**4-(Bromoacetyl)-N,N-diethylbenzenesulfonamide (82**). A solution of 4-Acetyl-N,N-diethyl-benzenesulfonamide (3.2 g, 12.5 mmol) was dissolved in dry DCM (150 mL) and cooled to 0°C. Bromine (2.0 g, 12.8 mmol) was added dropwise and the reaction was allowed to warm to ambient temperature before proceeding for 12 hr. The reaction was poured over cold $H_2O$ and an extraction performed against brine. The organics were dried over $Na_2SO_4$ before concentration to yield the crude reagent (4.3 g, 12.9 mmol).

**4-(2-Amino-1,3-thiazol-4-yl)-N,N-diethylbenzenesulfonamide (83).** A solution of 4-(Bromoacetyl)-N,N-diethylbenzenesulfonamide (4.3 g, 12 mmol) in THF (120 mL) was combined with thiourea (1.6 g, 21 mmol) and sodium bisulfate monohydrate (1.8 g, 13.2 mmol). The reaction was stirred for 12 h at ambient temperature. The THF was removed in vacuo and dissolved in DCM. This was washed with $H_2O$ and the organic fraction dried over $Na_2SO_4$ before concentration to yield the crude reagent in ~ 83% yield (3.2 g 10 mmol).

### TABLE 1 COMPOUNDS

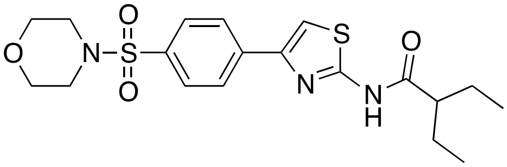

**2-ethyl-*N*-(4-(4-(morpholinosulfonyl)phenyl)thiazol-2-yl)butanamide (3).** The general three step synthesis sequence for the preparation of aryl thiazole **(50)** was followed to provide **(3).** HPLC/UV purity: 99%; LC-MS (ESI): 424.34 $[M+H]^+$. $^1$H NMR (400 MHz, CHLOROFORM-*D*) δ 11.97 (s, 1H), 8.10 (d, *J* = 8.4 Hz, 1H), 7.99 (d, *J* = 8.7 Hz, 1H), 7.84 (dd, *J* = 18.8, 8.2 Hz, 2H), 7.30 (s, 1H), 3.78 – 3.71 (m, 4H), 3.02 (dd, *J* = 5.9, 3.2 Hz, 4H), 2.45 (ddd, *J* = 14.0, 8.3, 5.6 Hz, 1H), 1.73 (dddd, *J* = 43.6, 23.5, 14.1, 7.3 Hz, 4H), 0.95 (q, *J* = 7.5 Hz, 6H). $^{13}$C NMR (101 MHz, CHLOROFORM-*D*) δ 198.57, 129.07, 128.53, 128.15, 126.76, 109.50, 66.16, 51.01, 46.08, 25.46, 25.05, 11.98, 11.85.

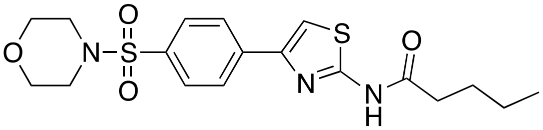

***N*-(4-(4-(morpholinosulfonyl)phenyl)thiazol-2-yl)pentanamide (4).** HPLC/UV purity: 97%; LC-MS (ESI): 410.3 $[M+H]^+$. $^1$H NMR (400 MHz, CHLOROFORM-*D*) δ 10.48 (s, 1H), 7.99 (s, 1H), 7.87 – 7.81 (m, 3H), 7.31 (s, 1H), 3.76 (m, 4H), 3.04 (m, 4H), 2.53 (dt, *J* = 30.0, 7.6 Hz, 2H), 1.83 – 1.70 (m, 3H), 1.50 – 1.36 (m, 3H), 0.97 (td, *J* = 7.4, 2.8 Hz, 3H). $^{13}$C NMR (101 MHz, CHLOROFORM-*D*) δ 171.42, 128.78, 128.55, 128.08, 126.67, 110.11, 66.17, 46.09, 36.25, 26.94, 22.31, 13.78.

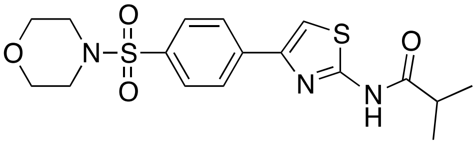

***N*-(4-(4-(morpholinosulfonyl)phenyl)thiazol-2-yl)isobutyramide (6).** The general three step synthesis sequence for the preparation of aryl thiazoles was followed to provide **(6).** HPLC/UV purity: 97%; LC-MS (ESI): 396.0 $[M+H]^+$. $^1$H NMR (400 MHz, CHLOROFORM-*D*) δ 10.25 (s, 1H), 7.99 (d, *J* = 8.3 Hz, 2H), 7.83 (d, *J* = 8.2 Hz, 1H), 7.31 (s, 1H), 3.79 – 3.72 (m, 4H), 3.04 (m, 4H), 2.76 (q, *J* = 6.9 Hz, 1H), 1.32 (dd, *J* = 13.2, 7.1 Hz, 6H). $^{13}$C NMR (101 MHz, CHLOROFORM-*D*) δ 175.16, 128.75, 128.55, 128.08, 126.65, 110.24, 66.18, 46.09, 35.76, 19.15, 19.08.

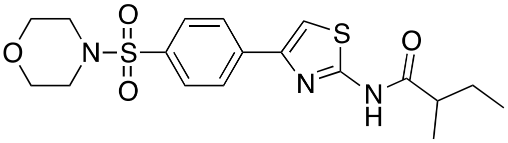

**2-methyl-*N*-(4-(4-(morpholinosulfonyl)phenyl)thiazol-2-yl)butanamide (7).** HPLC/UV purity: 90%; LC-MS (ESI): 410.3 $[M+H]^+$. $^1$H NMR (400 MHz, CHLOROFORM-*D*) δ 10.79 (s, 1H), 7.98 (d, *J* = 8.3 Hz, 2H), 7.82 (dd, *J* = 8.3, 6.0 Hz, 2H), 7.29 (s, 1H), 3.77 – 3.71 (m, 4H), 3.06 – 2.99 (m, 4H), 2.55 (m, *J* = 6.9 Hz, 1H), 1.91 – 1.73 (m, 1H), 1.70 – 1.55 (m, 1H), 1.28 (dd, *J* = 16.1,

6.9 Hz, 2H), 0.97 (t, *J* = 7.2 Hz, 3H). $^{13}$C NMR (101 MHz, CHLOROFORM-*D*) δ 207.47, 174.96, 128.84, 128.54, 128.09, 126.68, 110.02, 66.17, 46.09, 43.04, 27.12, 26.94, 16.74, 11.77, 11.70.

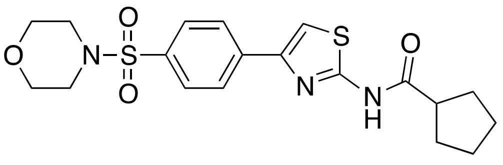

***N*-(4-(4-(morpholinosulfonyl)phenyl)thiazol-2-yl)cyclopentanecarboxamide (10).** HPLC/UV purity: 100%; LC-MS (ESI): 382.238 $[M+H]^+$. $^1$H NMR (400 MHz, DMSO-*D*) δ 8.11 (d, J = 8.4 Hz, 2H), 7.73 (d, J = 8.4 Hz, 2H), 7.43 (s, 1H), 3.63 (t, J = 4.4 Hz, 4H), 3.45 (t, J = 8.0 Hz, 4H), 2.87 (t, J = 8.0 Hz, 4H), 2.00 (m, 4H).

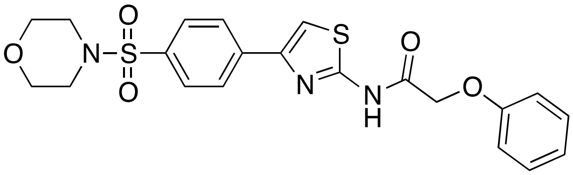

***N*-(4-(4-(morpholinosulfonyl)phenyl)thiazol-2-yl)-2-phenoxyacetamide (16).** HPLC/UV purity: 89%; LC-MS (ESI): 460.30 $[M+H]^+$. $^1$H NMR (400 MHz, DMSO-*D*) δ 12.48 (s, 1H), 8.17 (d, J = 8.4 Hz, 2H), 7.96 (s, 1H), 7.80 (d, J = 8.0 Hz, 2H), 7.33 (m, 2H), 6.97 (m, 3H), 3.65 (m, 4H), 2.90 (m, 4H), 2.50 (m, 4H).

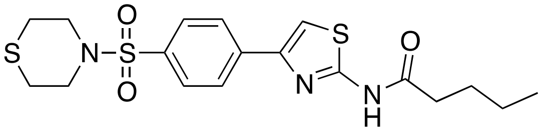

***N*-(4-(4-(thiomorpholinosulfonyl)phenyl)thiazol-2-yl)pentanamide (19).** HPLC/UV purity: 90%; LC-MS (ESI): 426.3 $[M+H]^+$. $^1$H NMR (400 MHz, CHLOROFORM-*D*) δ 7.97 (dd, *J* = 8.3, 5.4 Hz, 2H), 7.89 – 7.79 (m, 2H), 3.46 – 3.32 (m, 4H), 2.91 – 2.67 (m, 4H), 2.61 (td, *J* = 7.6, 3.8 Hz, 2H), 1.77 (p, *J* = 7.6 Hz, 2H), 1.49 – 1.35 (m, 2H), 0.95 (t, *J* = 7.3 Hz, 3H). $^{13}$C NMR (101 MHz, CHLOROFORM-*D*) δ 171.75, 128.58, 127.00, 126.82, 109.53, 47.99, 44.84, 36.49, 36.33, 27.42, 26.73, 22.26, 13.75.

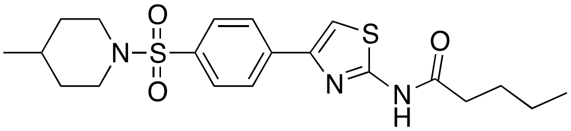

***N*-(4-(4-((4-methylpiperidin-1-yl)sulfonyl)phenyl)thiazol-2-yl)pentanamide (22).** HPLC/UV purity: 90%; LC-MS (ESI): 422.34 $[M+H]^+$.

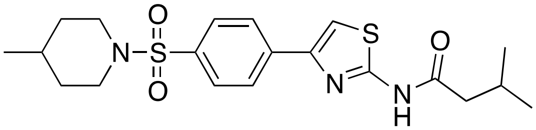

**3-methyl-*N*-(4-(4-((4-methylpiperidin-1-yl)sulfonyl)phenyl)thiazol-2-yl)butanamide (23).** HPLC/UV purity: 90%; LC-MS (ESI): 408.34 $[M+H]^+$. $^1$H NMR (400 MHz, CHLOROFORM-*D*) δ 7.96 (d, *J* = 8.6 Hz, 2H), 7.92 – 7.81 (m, 2H), 7.30 (s, 1H), 3.81 – 3.73 (m, 2H), 2.97 – 2.79 (m, 1H), 2.30 (t, *J* = 11.4 Hz, 3H), 1.42 – 1.24 (m, 10H), 0.92 (d, *J* = 5.5 Hz, 3H). $^{13}$C NMR (101 MHz,

CHLOROFORM-*D*) δ 175.79, 163.15, 138.95, 138.63, 130.67, 128.97, 128.31, 126.64, 108.88, 77.35, 77.24, 77.04, 76.72, 46.46, 35.92, 33.30, 30.07, 21.43, 18.79, 18.64.

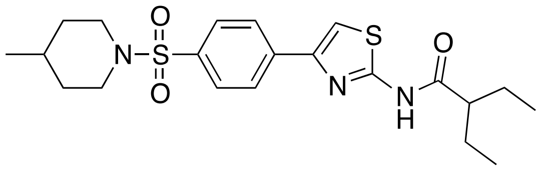

**2-ethyl-*N*-(4-(4-((4-methylpiperidin-1-yl)sulfonyl)phenyl)thiazol-2-yl)butanamide (24).** HPLC/UV purity: 99%; LC-MS (ESI): 436.21 $[M+H]^+$.

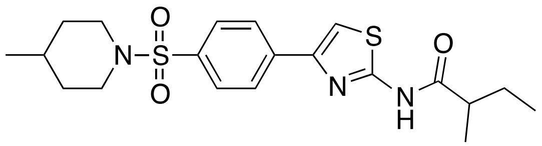

**2-methyl-*N*-(4-(4-((4-methylpiperidin-1-yl)sulfonyl)phenyl)thiazol-2-yl)butanamide (25).** HPLC/UV purity: 90%; LC-MS (ESI): 422.2 $[M+H]^+$. $^1$H NMR (400 MHz, CHLOROFORM-*D*) δ 12.39 (s, 1H), 8.08 – 8.01 (m, 2H), 8.00 – 7.86 (m, 3H), 3.78 (d, *J* = 11.3 Hz, 3H), 2.79 – 2.62 (m, 1H), 2.32 (t, *J* = 11.0 Hz, 3H), 1.87 (dt, *J* = 14.0, 7.0 Hz, 1H), 1.68 (t, *J* = 10.3 Hz, 2H), 1.40 – 1.29 (m, 7H), 1.04 – 0.90 (m, 6H).$^{13}$C NMR (101 MHz, CHLOROFORM-*D*) δ 128.61, 128.39, 46.55, 42.91, 33.41, 26.69, 21.54, 11.55.

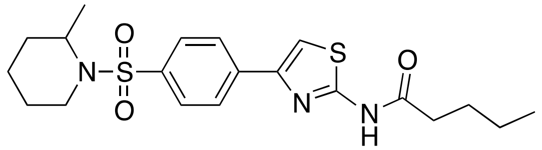

***N*-(4-(4-((2-methylpiperidin-1-yl)sulfonyl)phenyl)thiazol-2-yl)pentanamide (29).** HPLC/UV purity: 97%; LC-MS (ESI): 422.31 $[M+H]^+$. $^1$H NMR (400 MHz, CHLOROFORM-*D*) δ 7.94 (s, 3H), 7.31 (s, 1H), 3.78 – 3.70 (m, 1H), 3.07 – 2.96 (m, 1H), 2.74 – 2.65 (m, 2H), 2.63 (s, 9H), 1.84 – 1.76 (m, 1H), 1.63 – 1.52 (m, 3H), 1.50 – 1.40 (m, 2H), 1.39 (s, 1H), 1.08 (s, 1H), 0.97 (t, *J* = 7.3 Hz, 2H). $^{13}$C NMR (101 MHz, CHLOROFORM-*D*) δ 171.90, 162.10, 142.99, 140.33, 131.41, 128.14, 126.59, 108.90, 57.78, 48.78, 41.01, 40.48, 36.62, 36.24, 30.38, 26.57, 25.27, 22.15, 18.06, 15.57, 13.65.

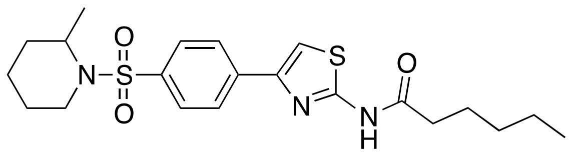

***N*-(4-(4-((2-methylpiperidin-1-yl)sulfonyl)phenyl)thiazol-2-yl)hexanamide (31).** HPLC/UV purity: 98%; LC-MS (ESI): 436.18 $[M+H]^+$. $^1$H NMR (400 MHz, CHLOROFORM-*D*) δ 11.12 (s, 1H), 7.92 (s, 4H), 7.27 (s, 1H), 4.31 – 4.23 (m, 1H), 3.79 – 3.69 (m, 1H), 3.07 – 2.96 (m, 1H), 2.57 (t, *J* = 7.6 Hz, 2H), 1.82 – 1.72 (m, 2H), 1.64 – 1.44 (m, 4H), 1.37 (dt, *J* = 7.6, 4.0 Hz, 6H), 1.09 (d, *J* = 6.9 Hz, 3H), 0.96 – 0.87 (m, 3H). $^{13}$C NMR (101 MHz, CHLOROFORM-*D*) δ 171.63, 160.40, 141.88, 128.47, 127.97, 127.20, 126.59, 109.43, 48.79, 40.52, 36.50, 31.24, 30.44, 25.33, 24.56, 22.38, 18.18, 15.60, 13.95.

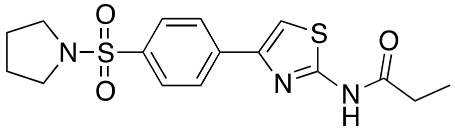

***N*-(4-(4-(pyrrolidin-1-ylsulfonyl)phenyl)thiazol-2-yl)propionamide (33).** HPLC/UV purity: 90%; LC-MS (ESI): 366.25 $[M+H]^+$. $^1$H NMR (400 MHz, CHLOROFORM-*D*) δ 7.97 (s, 5H), 7.29 (d, *J* = 17.3 Hz, 2H), 3.28 (td, *J* = 5.8, 3.0 Hz, 5H), 2.76 (q, *J* = 7.5 Hz, 2H), 1.86 – 1.75 (m, 2H), 1.36 (t, *J* = 7.5 Hz, 4H). $^{13}$C NMR (101 MHz, CHLOROFORM-*D*) δ 172.59, 162.95, 139.26, 138.92, 130.57, 128.77, 128.44, 128.16, 126.75, 108.82, 48.04, 29.97, 25.34, 8.54.

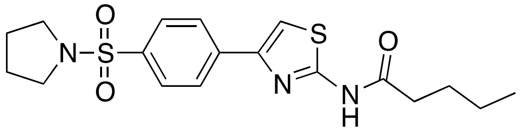

***N*-(4-(4-(pyrrolidin-1-ylsulfonyl)phenyl)thiazol-2-yl)pentanamide (35).** HPLC/UV purity: 95%; LC-MS (ESI): 394.3 $[M+H]^+$. $^1$H NMR (400 MHz, CHLOROFORM-*D*) δ 8.01 – 7.90 (m, 5H), 7.28 (s, 1H), 3.33 – 3.24 (m, 5H), 2.56 (d, *J* = 7.6 Hz, 2H), 1.82 – 1.70 (m, 6H), 1.47 – 1.37 (m, 2H), 0.96 (t, *J* = 7.3 Hz, 3H). $^{13}$C NMR (101 MHz, CHLOROFORM-*D*) δ 171.47, 136.90, 128.34, 127.67, 126.52, 109.68, 48.02, 36.13, 26.88, 25.29, 22.23, 13.71.

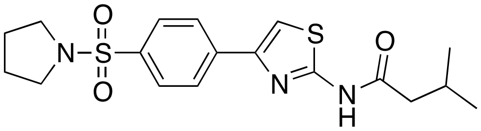

**3-methyl-*N*-(4-(4-(pyrrolidin-1-ylsulfonyl)phenyl)thiazol-2-yl)butanamide (36).** $^1$H NMR (400 MHz, CHLOROFORM-*D*) δ 12.31 (s, 1H), 8.00 (d, *J* = 7.8 Hz, 1H), 7.94 – 7.87 (m, 1H), 7.40 (d, *J* = 2.9 Hz, 1H), 4.08 (s, 8H), 3.82 (t, *J* = 10.9 Hz, 1H), 3.77 – 3.63 (m, 1H), 3.27 (t, *J* = 6.1 Hz, 2H), 3.07 – 2.96 (m, 2H), 2.54 – 2.47 (m, 1H), 1.81 – 1.74 (m, 1H), 1.04 (dd, *J* = 6.7, 1.9 Hz, 3H), 0.95 (dd, *J* = 7.3, 1.6 Hz, 1H).

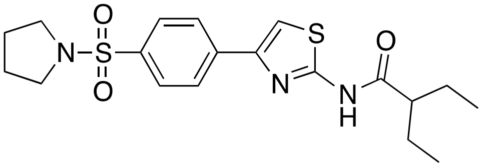

**2-ethyl-*N*-(4-(4-(pyrrolidin-1-ylsulfonyl)phenyl)thiazol-2-yl)butanamide (37).** HPLC/UV purity: 100%; LC-MS (ESI): 408.27. $^1$H NMR (400 MHz, CHLOROFORM-*D*) δ 7.99 (dd, *J* = 8.4, 2.8 Hz, 2H), 7.90 (dd, *J* = 8.4, 2.9 Hz, 1H), 7.29 (s, 1H), 3.28 (td, *J* = 6.6, 2.9 Hz, 4H), 1.77 (dq, *J* = 10.6, 5.2 Hz, 4H), 1.36 (dq, *J* = 7.3, 3.8 Hz, 6H), 0.91 (h, *J* = 2.9 Hz, 4H).

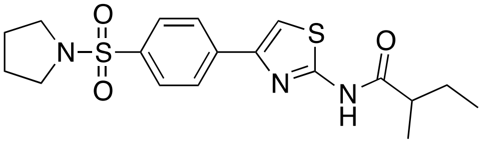

**2-methyl-*N*-(4-(4-(pyrrolidin-1-ylsulfonyl)phenyl)thiazol-2-yl)butanamide (38).** HPLC/UV purity: 99%; LC-MS (ESI): 394.23 $[M+H]^+$. $^1$H NMR (400 MHz, CHLOROFORM-*D*) δ 7.96 (d, *J* = 8.5 Hz, 1H), 7.86 (d, *J* = 8.5 Hz, 1H), 7.32 (s, 1H), 3.77 (d, *J* = 11.1 Hz, 1H), 2.63 (s, 8H), 2.29 (s, 1H), 1.85 – 1.77 (m, 1H), 1.42 – 1.31 (m, 3H), 1.31 (s, 2H), 0.92 (p, *J* = 3.4 Hz, 4H). $^{13}$C NMR

(101 MHz, CHLOROFORM-*D*) δ 171.86, 161.61, 137.61, 128.73, 126.56, 109.29, 46.46, 40.99, 36.42, 33.30, 31.11, 30.08, 24.32, 22.27, 21.44, 13.85.

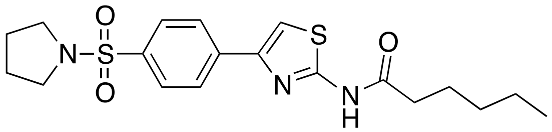

***N*-(4-(4-(pyrrolidin-1-ylsulfonyl)phenyl)thiazol-2-yl)hexanamide (39).** HPLC/UV purity: 99%; LC-MS (ESI): 408.23 $[M+H]^+$. $^1H$ NMR (400 MHz, DMSO-$D_6$) δ 12.03 (s, 1H), 8.12 (d, J = 8.4 Hz, 2H), 7.86 (d, J = 8.4 Hz, 2H), 3.71 (m, 4H), 2.46 (m, 2H), 1.64 (m, 4H), 0.87 (m, 3H).

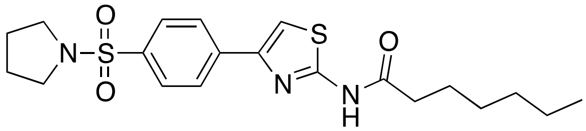

***N*-(4-(4-(pyrrolidin-1-ylsulfonyl)phenyl)thiazol-2-yl)heptanamide (40).** HPLC/UV purity: 99%; LC-MS (ESI): 422.04 $[M+H]^+$. $^1H$ NMR (400 MHz, DMSO-$D_6$) δ 8.12 (d, *J* = 8.3 Hz, 3H), 7.89 – 7.83 (m, 4H), 3.16 (dd, *J* = 9.5, 3.9 Hz, 3H), 2.53 – 2.43 (m, 3H), 1.69 – 1.59 (m, 6H), 1.28 (d, *J* = 4.3 Hz, 5H), 0.86 (d, *J* = 6.7 Hz, 3H).$^{13}C$ NMR (101 MHz, DMSO-$D_6$) δ 171.68, 158.23, 146.94, 138.11, 134.70, 127.88, 126.11, 110.79, 47.73, 34.78, 30.84, 28.10, 24.60, 24.49, 21.84, 13.83.

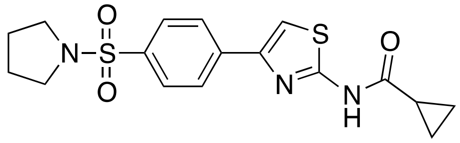

***N*-(4-(4-(pyrrolidin-1-ylsulfonyl)phenyl)thiazol-2-yl)cyclopropanecarboxamide (41).** HPLC/UV purity: 90%; LC-MS (ESI): 378.26 $[M+H]^+$. $^1H$ NMR (400 MHz, CHLOROFORM-*D*) δ 7.96 (s, 4H), 3.32 – 3.24 (m, 6H), 1.80 (h, *J* = 3.3 Hz, 4H), 1.30 (dd, *J* = 4.5, 3.2 Hz, 2H), 1.20 (ddd, *J* = 14.9, 7.5, 3.7 Hz, 3H). $^{13}C$ NMR (101 MHz, CHLOROFORM-*D*) δ 172.73, 128.67, 128.40, 127.75, 126.67, 108.92, 48.03, 25.33, 25.29, 15.69, 11.06.

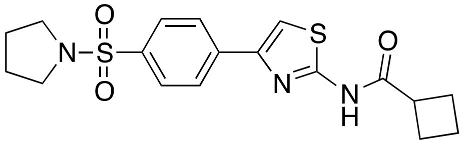

***N*-(4-(4-(pyrrolidin-1-ylsulfonyl)phenyl)thiazol-2-yl)cyclobutanecarboxamide (42). HPLC**/UV purity: 85%; LC-MS (ESI): 392.3 $[M+H]^+$. $^1H$ NMR (400 MHz, CHLOROFORM-*D*) δ 7.95 (s, 4H), 3.32 – 3.24 (m, 6H), 2.51 – 2.31 (m, 5H), 2.16 – 1.96 (m, 2H), 1.79 (dd, *J* = 10.2, 3.4 Hz, 3H). $^{13}C$ NMR (101 MHz, CHLOROFORM-*D*) δ 173.48, 161.56, 141.75, 138.06, 133.09, 128.54, 128.38, 127.68, 126.61, 109.29, 48.03, 39.69, 25.31, 25.27, 25.16, 18.26.

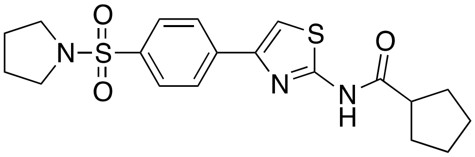

***N*-(4-(4-(pyrrolidin-1-ylsulfonyl)phenyl)thiazol-2-yl)cyclopentanecarboxamide (43).** HPLC/UV purity: 90%; LC-MS (ESI): 406.31 $[M+H]^+$. $^1H$ NMR (400 MHz, CHLOROFORM-*D*)

δ 11.50 (s, 1H), 7.98 – 7.87 (m, 4H), 7.26 (dd, *J* = 4.4, 2.5 Hz, 2H), 3.28 (tt, *J* = 5.8, 3.1 Hz, 4H), 2.98 (t, *J* = 7.8 Hz, 1H), 2.11 – 2.03 (m, 2H), 1.85 – 1.74 (m, 7H), 1.69 (dt, *J* = 13.0, 4.2 Hz, 3H). $^{13}$C NMR (101 MHz, CHLOROFORM-*D*) δ 128.69, 128.45, 126.71, 48.11, 45.95, 30.18, 26.07, 26.01, 25.40.

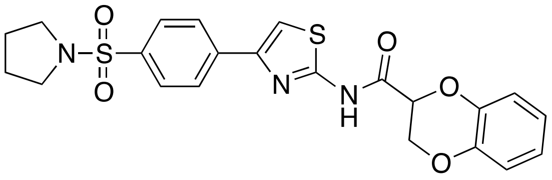

***N*-(4-(4-(pyrrolidin-1-ylsulfonyl)phenyl)thiazol-2-yl)-2,3dihydrobenzo[*b*][1,4]dioxine -2-carboxamide (45).** HPLC/UV purity: 100%; LC-MS (ESI): 472.047 [M+H]$^+$. $^1$H NMR (400 MHz, DMSO-*D*) δ 12.67 (s, 1H), 8.15-8.12 (m, 2H), 7.95 (s, 1H), 7.88-7.86 (m, 2H), 7.03 (m, 1H), 6.86 (m, 3H), 5.19 (s, 1H), 4.47 (m, 2H), 3.19 (m, 4H), 1.66 (m, 4H).

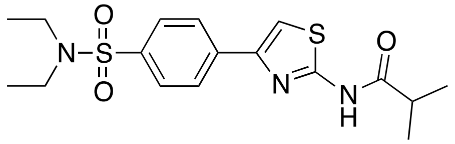

***N*-(4-(4-(*N*,*N*-diethylsulfamoyl)phenyl)thiazol-2-yl)isobutyramide (47).** HPLC/UV purity: 95%; LC-MS (ESI): 382.29 [M+H]$^+$. $^1$H NMR (400 MHz, CHLOROFORM-*D*) δ 7.95 (d, *J* = 1.9 Hz, 3H), 7.27 (s, 1H), 3.28 (q, *J* = 7.2 Hz, 5H), 2.94 – 2.88 (m, 1H), 2.67 (d, *J* = 4.0 Hz, 3H), 1.39 (s, 3H), 1.15 (t, *J* = 7.1 Hz, 6H). $^{13}$C NMR (101 MHz, CHLOROFORM-*D*) δ 175.79, 163.25, 142.71, 138.73, 130.10, 128.36, 126.68, 108.62, 42.33, 42.19, 40.65, 35.94, 18.62, 14.34, 14.21.

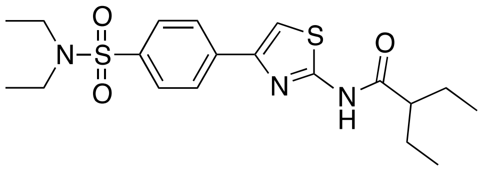

***N*-(4-(4-(*N*,*N*-diethylsulfamoyl)phenyl)thiazol-2-yl)-2-ethylbutanamide (49).** HPLC/UV purity: 95%; LC-MS (ESI): 410.23 [M+H]$^+$. $^1$H NMR (400 MHz, CHLOROFORM-*D*) δ 8.04 (s, 1H), 8.02 (s, 1H), 7.98 – 7.92 (m, 3H), 3.33 – 3.26 (m, 4H), 3.25 (s, 1H), 2.77 (s, 1H), 1.77 (dq, *J* = 21.3, 6.9 Hz, 3H), 1.16 (t, *J* = 7.1 Hz, 6H), 0.96 (t, *J* = 7.4 Hz, 6H). $^{13}$C NMR (101 MHz, CHLOROFORM-*D*) δ 128.45, 42.30, 14.34, 11.85.

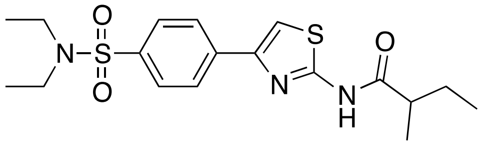

***N*-(4-(4-(*N*,*N*-diethylsulfamoyl)phenyl)thiazol-2-yl)-2-methylbutanamide (50).** HPLC/UV purity: 95%; LC-MS (ESI): 396.33 [M+H]$^+$. $^1$H NMR (400 MHz, CHLOROFORM-*D*) δ 9.79 – 9.70 (m, 1H), 7.96 - 7.87 (m, 2H), 7.25 (m, 2H), 3.25 (dt, *J* = 14.7, 7.2 Hz, 4H), 1.26 – 1.19 (m, 3H), 1.11 (m, 6H), 0.97 – 0.86 (m, 3H). $^{13}$C NMR (101 MHz, CHLOROFORM-*D*) δ 174.90, 158.52, 148.01, 139.26, 138.20, 127.70, 126.55, 110.22, 42.78, 42.13, 27.19, 17.02, 14.20, 11.81.

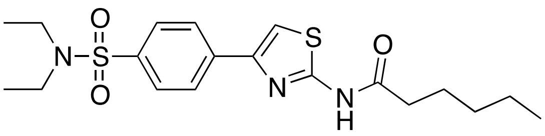

***N*-(4-(4-(*N*,*N*-diethylsulfamoyl)phenyl)thiazol-2-yl)hexanamide (51).** HPLC/UV purity: 95%; LC-MS (ESI): 410.23 $[M+H]^+$. $^1H$ NMR (400 MHz, CHLOROFORM-*D*) δ 10.97 (s, 1H), 8.09 (d, *J* = 10.2 Hz, 0H), 7.98 (q, *J* = 7.8 Hz, 1H), 7.87 (q, *J* = 7.6 Hz, 2H), 7.29 (d, *J* = 6.4 Hz, 2H), 3.77 (d, *J* = 11.9 Hz, 2H), 2.82 – 2.77 (m, 4H), 2.59 (q, *J* = 7.2 Hz, 1H), 2.50 (d, *J* = 7.9 Hz, 1H), 2.30 (s, 2H), 1.77 (dt, *J* = 15.1, 7.6 Hz, 1H), 1.44 (p, *J* = 7.0 Hz, 2H), 1.32 (s, 3H), 0.95 (dq, *J* = 18.5, 5.5 Hz, 5H). $^{13}C$ NMR (101 MHz, CHLOROFORM-*D*) δ 171.48, 128.58, 126.47, 109.56, 46.47, 36.17, 33.32, 30.11, 26.80, 22.21, 21.45, 13.70.

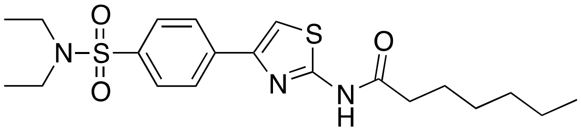

***N*-(4-(4-(*N*,*N*-diethylsulfamoyl)phenyl)thiazol-2-yl)heptanamide (52).** HPLC/UV purity: 95%; LC-MS (ESI): 424.27 $[M+H]^+$. $^1H$ NMR (400 MHz, CHLOROFORM-*D*) δ 10.08 (s, 1H), 8.01 – 7.94 (m, 1H), 7.85 (dd, *J* = 8.5, 2.7 Hz, 2H), 7.32 – 7.24 (m, 1H), 3.82 – 3.74 (m, 2H), 2.50 (dt, *J* = 7.5, 3.7 Hz, 2H), 2.28 (t, *J* = 11.1 Hz, 2H), 1.80 – 1.64 (m, 4H), 1.41 – 1.26 (m, 9H), 0.90 (tt, *J* = 6.7, 2.8 Hz, 6H). $^{13}C$ NMR (101 MHz, CHLOROFORM-*D*) δ 171.36, 128.39, 126.41, 109.96, 46.48, 36.38, 33.32, 31.43, 30.12, 28.79, 24.91, 22.46, 21.45, 14.03.

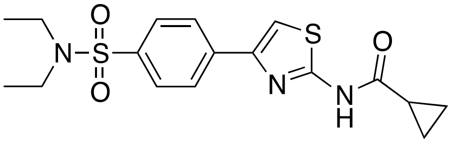

***N*-(4-(4-(*N*,*N*-diethylsulfamoyl)phenyl)thiazol-2-yl)cyclopropanecarboxamide (53).** HPLC/UV purity: 90%; LC-MS (ESI): 380.25 $[M+H]^+$. $^1H$ NMR (400 MHz, CHLOROFORM-*D*) δ 7.96 (m, 3H), 7.27 (d, *J* = 3.2 Hz, 2H), 3.33 – 3.25 (m, 4H), 2.63 (d, *J* = 4.5 Hz, 1H), 1.33 (m, 1H), 1.25 – 1.12 (m, 7H). $^{13}C$ NMR (101 MHz, CHLOROFORM-*D*) δ 128.37, 126.69, 42.21, 15.87, 14.23, 11.43.

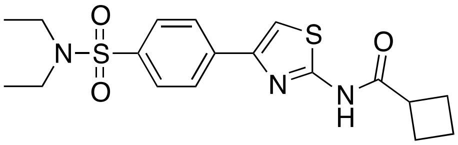

***N*-(4-(4-(*N*,*N*-diethylsulfamoyl)phenyl)thiazol-2-yl)cyclobutanecarboxamide (54).** HPLC/UV purity: 91%; LC-MS (ESI): 394.30 $[M+H]^+$. $^1H$ NMR (400 MHz, CHLOROFORM-*D*) δ 7.93 (m, 4H), 7.30 (m, 1H), 3.52 – 3.43 (m, 1H), 3.27 (q, *J* = 7.2 Hz, 4H), 2.63 (s, 4H), 2.50 – 2.35 (m, 3H), 2.11 (d, *J* = 9.0 Hz, 1H), 1.15 (t, *J* = 7.1 Hz, 6H). $^{13}C$ NMR (101 MHz, CHLOROFORM-*D*) δ 173.63, 162.27, 142.03, 140.36, 131.56, 128.18, 126.65, 108.97, 42.17, 40.96, 39.73, 25.19, 18.27, 14.20.

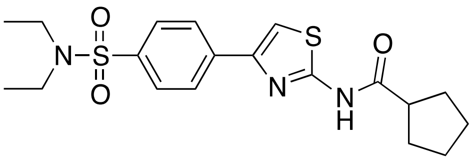

***N*-(4-(4-(*N*,*N*-diethylsulfamoyl)phenyl)thiazol-2-yl)cyclopentanecarboxamide (55).** HPLC/UV purity: 96%; LC-MS (ESI): 408.3 $[M+H]^+$. $^1H$ NMR (400 MHz, CHLOROFORM-*D*) δ 11.93 (s, 1H), 8.08 (d, *J* = 8.3 Hz, 1H), 7.96 (d, *J* = 8.2 Hz, 1H), 7.88 (s, 1H), 7.87 – 7.80 (m, 1H), 7.29 (s, 1H), 3.03 (t, *J* = 5.4 Hz, 4H), 2.65 (d, *J* = 7.0 Hz, 1H), 1.66 (p, *J* = 5.6 Hz, 5H), 1.46 (d, *J* = 5.6 Hz, 1H), 1.45 – 1.41 (m, 1H), 1.35 (s, 1H), 1.33 (s, 1H), 1.29 (d, *J* = 6.9 Hz, 1H), 0.99 (q, *J* = 7.5 Hz, 3H). $^{13}C$ NMR (101 MHz, CHLOROFORM-*D*) δ 128.77, 128.29, 127.88, 126.55, 109.19, 46.98, 42.97, 42.73, 27.01, 26.70, 25.16, 23.47, 16.80, 16.47, 11.67, 11.54.

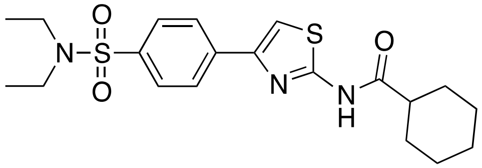

***N*-(4-(4-(*N*,*N*-diethylsulfamoyl)phenyl)thiazol-2-yl)cyclohexanecarboxamide (56).** HPLC/UV purity: 95%; LC-MS (ESI): 396.33 $[M+H]^+$. $^1H$ NMR (400 MHz, CHLOROFORM-*D*) δ 7.95 (m, 5H), 3.28 (q, *J* = 7.1 Hz, 3H), 2.10 (d, *J* = 13.0 Hz, 2H), 1.94 – 1.82 (m, 3H), 1.68 – 1.57 (m, 3H), 1.48 – 1.22 (m, 5H), 1.15 (t, *J* = 7.0 Hz, 7H). $^{13}C$ NMR (101 MHz, CHLOROFORM-*D*) δ 179.96, 174.88, 162.96, 142.41, 139.27, 130.67, 128.29, 126.72, 108.90, 44.91, 42.79, 42.21, 40.87, 28.91, 28.71, 25.73, 25.39, 25.35, 25.29, 25.18, 14.22.

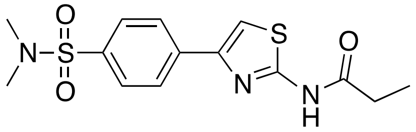

***N*-(4-(4-(*N*,*N*-dimethylsulfamoyl)phenyl)thiazol-2-yl)propionamide (57).** HPLC/UV purity: 99%; LC-MS (ESI): 340.2 $[M+H]^+$. $^1H$ NMR (400 MHz, CHLOROFORM-*D*) δ 8.01 – 7.95 (m, 1H), 7.89 (d, *J* = 8.3 Hz, 1H), 7.27 (d, *J* = 1.9 Hz, 1H), 2.77 – 2.69 (m, 6H), 2.63 (d, *J* = 1.8 Hz, 6H), 1.34 (t, *J* = 7.5 Hz, 2H). $^{13}C$ NMR (101 MHz, CHLOROFORM-*D*) δ 172.51, 161.96, 140.82, 137.05, 132.49, 128.85, 126.70, 109.32, 42.69, 40.95, 37.91, 29.83, 8.65.

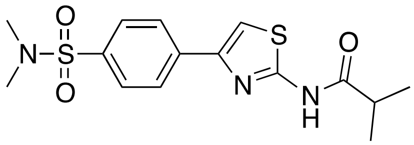

**s**

***N*-(4-(4-(*N*,*N*-dimethylsulfamoyl)phenyl)thiazol-2-yl)isobutyramide (58).** HPLC/UV purity: 97%; LC-MS (ESI): 354.24 $[M+H]^+$. $^1H$ NMR (400 MHz, CHLOROFORM-*D*) δ 8.02 – 7.96 (m, 1H), 7.92 – 7.85 (m, 1H), 2.75 (s, 4H), 2.63 (s, 11H), 1.35 (s, 2H). $^{13}C$ NMR (101 MHz, CHLOROFORM-*D*) δ 128.75, 126.60, 109.55, 40.98, 37.92, 35.68, 18.83.

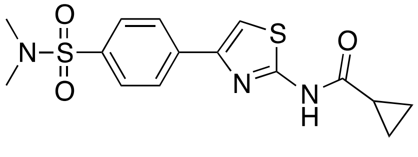

***N*-(4-(4-(*N*,*N*-dimethylsulfamoyl)phenyl)thiazol-2-yl)cyclopropanecarboxamide (59).** HPLC/UV purity: 90%; LC-MS (ESI): 352.21 $[M+H]^+$. $^1H$ NMR (400 MHz, CHLOROFORM-*D*) δ 8.00 – 7.90 (m, 2H), 7.86 (m, 1H), 7.82 (m, 1H), 7.32 – 7.22 (m, 1H), 2.77 – 2.67 (m, 4H), 2.67 – 2.57 (m, 6H), 1.25 – 1.15 (m, 2H), 1.07 – 0.99 (m, 1H). $^{13}C$ NMR (101 MHz, CHLOROFORM-

*D*) δ 172.67, 160.69, 143.48, 135.76, 135.02, 128.56, 126.60, 126.56, 109.68, 40.90, 40.86, 37.94, 15.13, 10.18.

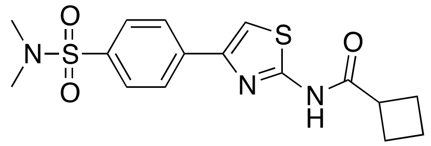

***N*-(4-(4-(*N*,*N*-dimethylsulfamoyl)phenyl)thiazol-2-yl)cyclobutanecarboxamide (60).** HPLC/UV purity: 90%; LC-MS (ESI): 366.25 $[M+H]^+$. $^1$H NMR (400 MHz, CHLOROFORM-*D*) δ 7.99 (d, *J* = 8.5 Hz, 2H), 7.93 (d, *J* = 8.3 Hz, 2H), 7.33 (s, 1H), 2.76 (s, 4H), 2.64 (s, 6H), 2.53 – 2.38 (m, 3H), 2.19 – 1.98 (m, 1H). $^{13}$C NMR (101 MHz, CHLOROFORM-*D*) δ 173.75, 163.13, 138.77, 137.88, 130.73, 129.04, 126.72, 108.94, 40.88, 39.78, 37.89, 25.22, 18.28.

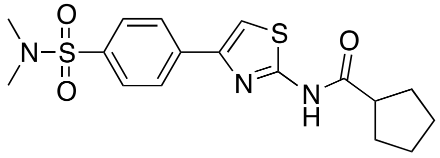

***N*-(4-(4-(*N*,*N*-dimethylsulfamoyl)phenyl)thiazol-2-yl)cyclopentanecarboxamide (61).** HPLC/UV purity: 90%; LC-MS (ESI): 380.29 $[M+H]^+$. $^1$H NMR (400 MHz, CHLOROFORM-*D*) δ 7.96 (s, 3H), 7.29 (d, *J* = 16.6 Hz, 1H), 3.32 – 3.24 (m, 5H), 2.89 (hept, *J* = 6.9 Hz, 1H), 1.80 (h, *J* = 6.4 Hz, 3H), 1.38 (d, *J* = 6.8 Hz, 6H), 1.32 (m, 2H). $^{13}$C NMR (101 MHz, CHLOROFORM-*D*) δ 175.64, 162.34, 140.43, 138.61, 131.94, 128.65, 128.39, 127.72, 126.67, 109.14, 48.04, 35.85, 25.33, 19.02, 18.73.

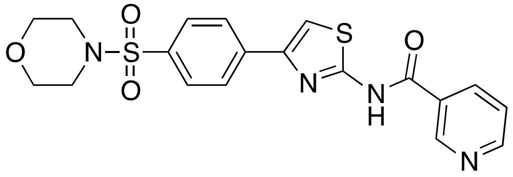

***N*-(4-(4-(morpholinosulfonyl)phenyl)thiazol-2-yl)nicotinamide (62).** HPLC/UV purity: 94%; LC-MS (ESI): 388.2 $[M+H]^+$. $^1$H NMR (400 MHz, CHLOROFORM-*D*) δ 9.48 (d, *J* = 5.5 Hz, 1H), 9.00 (s, 1H), 8.85 (s, 1H), 8.17 (d, *J* = 7.8 Hz, 2H), 8.09 (s, 1H), 7.92 (dd, *J* = 17.0, 6.5 Hz, 3H), 7.27 (s, 2H), 3.79 (dt, *J* = 8.8, 4.8 Hz, 4H), 3.08 (s, 4H). $^{13}$C NMR (101 MHz, CHLOROFORM-*D*) δ 128.73, 127.25, 66.19, 46.10.

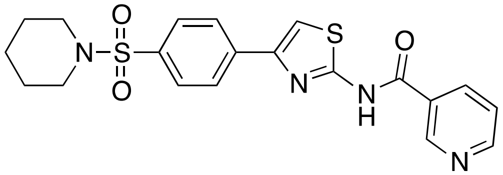

***N*-(4-(4-(piperidin-1-ylsulfonyl)phenyl)thiazol-2-yl)nicotinamide (63).** HPLC/UV purity: 98%; LC-MS (ESI): 387.24 $[M+H]^+$.$^1$H NMR (400 MHz, DMSO-*D*) δ 9.25 (s, 1H), 8.72 (m, 1H), 8.53 (s, 1H), 8.43 (m, 2H), 7.84 (m, 2H), 7.63 (m, 1.0), 2.91 (m, 4H), 1.57 (m, 4H), 1.38 (m, 2H).

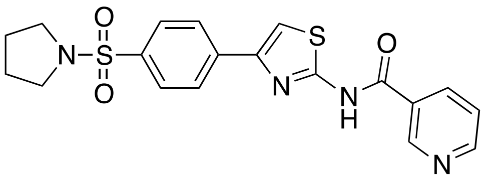

***N*-(4-(4-(pyrrolidin-1-ylsulfonyl)phenyl)thiazol-2-yl)nicotinamide (64).** HPLC/UV purity: 100%; LC-MS (ESI): 373.04 $[M+H]^+$. $^1$H NMR (400 MHz, DMSO-*D*) δ 9.25 (s, 1H), 8.73 (t, J = 1.2 Hz, 1H), 8.53 (s, 1H), 8.42 (m, 1H), 8.30 (t, J = 6.8 Hz, 2H), 7.92 (d, J = 8.4 Hz, 2H), 7.60 (m, 1H), 3.19 (m, 4H), 1.67 (m, 4H).

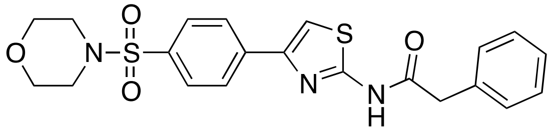

***N*-(4-(4-(morpholinosulfonyl)phenyl)thiazol-2-yl)-2-phenylacetamide (65).** HPLC/UV purity: 95%; LC-MS (ESI): 401.2 $[M+H]^+$. $^1$H NMR (400 MHz, CHLOROFORM-*D*) δ 8.16 – 8.09 (m, 2H), 7.86 – 7.78 (m, 3H), 7.55 (s, 1H), 7.41 – 7.30 (m, 6H), 4.51 (s, 1H), 3.79 – 3.72 (m, 4H), 3.03 (dt, *J* = 4.8, 3.4 Hz, 4H). $^{13}$C NMR (101 MHz, CHLOROFORM-*D*) δ 172.60, 152.41, 134.47, 129.15, 129.08, 129.02, 128.77, 128.47, 127.96, 127.57, 127.04, 115.78, 66.11, 46.01, 39.41.

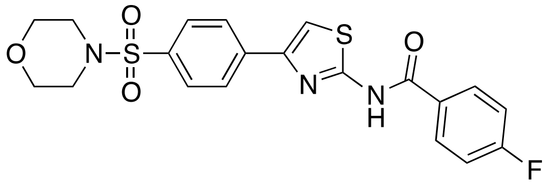

**4-fluoro-*N*-(4-(4-(morpholinosulfonyl)phenyl)thiazol-2-yl)benzamide (66).** $^1$H NMR (400 MHz, $CDCl_3$) δ, ppm: 8.17 (d, 2H), 8.03 (dd. 2H), 7.83 (d, 2H), 7.64 (s, 1H), 7.18 (t, 2H), 3.75 (dd, 4H), 3.04 (dd, 4H). $^{13}$C NMR (400 MHz, $CDCl_3$) δ, ppm: 167.45, 165.40, 162.82, 154.28, 138.72, 134.19, 128.66, 128.56, 128.41, 126.92, 116.33, 116.01, 115.15, 66.11, 46.02. HRMS (ESI): *m/z* calcd for $C_{19}H_{17}FN_2O_3S_2$ 404.48, found $(M + H)^+$ 405.08.

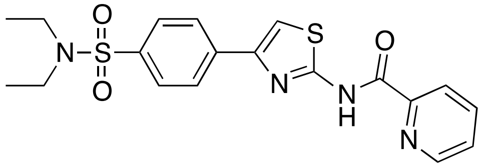

***N*-(4-(4-(*N*,*N*-diethylsulfamoyl)phenyl)thiazol-2-yl)picolinamide (67).** HPLC/UV purity: 98%; LC-MS (ESI): 374.15 $[M+H]^+$. $^1$H NMR (400 MHz, DMSO-*D*) δ 8.66 (d, *J* = 8.0 Hz, 1H), 8.49 (s, 1H), 8.26 (m, 3H), 8.02 (m, 1H), 7.98 (m, 2H), 7.55 (m, 1H) 3.19 (m, 4H), 1.05 (t, J = 4 Hz, 6H). $^{13}$C NMR (101 MHz, DMSO-*D*) δ 168.83, 153.82, 150.09, 149.80, 138.98, 137.88, 137.49, 127.40, 126.73, 125.47, 119.65, 119.36, 41.76, 14.03.

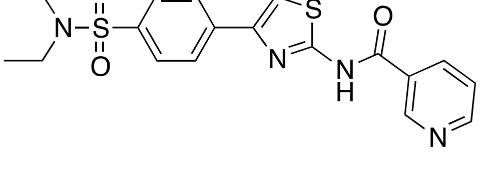

***N*-(4-(4-(*N*,*N*-diethylsulfamoyl)phenyl)thiazol-2-yl)nicotinamide (68).** HPLC/UV purity: 98%; LC-MS (ESI): 374.60 $[M+H]^+$. $^1$H NMR (400 MHz, DMSO-*D*) δ 9.25 (d, *J* = 4.0 Hz, 1H), 8.71 (s, 1H), 8.51 (s, 1H), 8.27 (t, J = 2.0 Hz, 2H), 7.89 (d, J = 8.0 Hz, 2H), 4.13 (m, 4H) 1.05 (m, 6H). $^{13}$C NMR (101 MHz, DMSO-*D*) δ 168.83, 153.82, 150.09, 149.80, 138.98, 137.89, 137.49, 127.41, 126.73, 125.47, 119.64, 119.36, 41.76, 14.03.

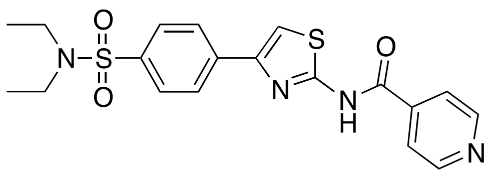

***N*-(4-(4-(*N*,*N*-diethylsulfamoyl)phenyl)thiazol-2-yl)isonicotinamide (69).** HPLC/UV purity: 95%; LC-MS (ESI): 373.8 $[M+H]^+$. $^1$H NMR (400 MHz, CHLOROFORM-*D*) δ 8.89 (s, 2H), 8.51 (s, 2H), 8.13 (dd, *J* = 8.5, 3.2 Hz, 1H), 7.98 – 7.91 (m, 2H), 7.27 (d, *J* = 2.3 Hz, 2H), 3.31 (qd, *J* = 7.2, 3.2 Hz, 4H), 1.17 (td, *J* = 7.0, 3.0 Hz, 6H). $^{13}$C NMR (101 MHz, CHLOROFORM-*D*) δ 141.82, 127.83, 127.17, 123.11, 42.05, 14.15.

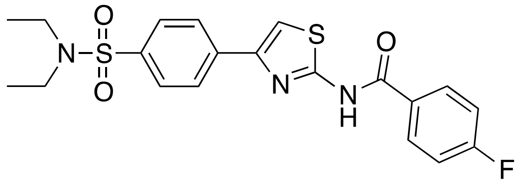

***N*-(4-(4-(*N*,*N*-diethylsulfamoyl)phenyl)thiazol-2-yl)-4-fluorobenzamide (70).** $^1$H NMR (400 MHz, $CDCl_3$) δ, ppm: 8.12 (d, 2H), 8.04 (m, 2H), 7.88 (d, 2H), 7.61 (s, 1H), 7.17 (t, 2H), 3.27 (q, 4H), 1.15 (t, 6H). $^{13}$C NMR (400 MHz, $CDCl_3$) δ, ppm: 167.30, 165.30, 162.80, 154.45, 139.62, 137.89, 128.52, 127.53, 126.82, 116.22, 116.00, 114.78, 41.98, 14.10. HRMS (ESI): *m/z* calcd for $C_{19}H_{19}FN_2O_2S_2$ 390.09, found $(M + H)^+$ 391.09.

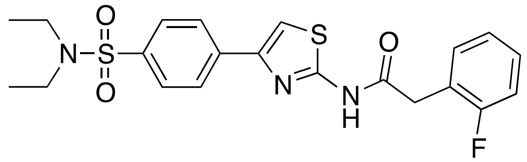

***N*-(4-(4-(*N*,*N*-diethylsulfamoyl)phenyl)thiazol-2-yl)-2-(2-fluorophenyl)acetamide (71).** $^1$H NMR (400 MHz, $CDCl_3$) δ, ppm: 8.02 (d, 2H), 7.86 (d, 2H), 7.64 (s, 1H), 7.51 (m, 4H), 4.45 (s, 2H), 3.26 (m, 4H), 1.13 (m, 6H). $^{13}$C NMR (400 MHz, $CDCl_3$) δ, ppm: 169.32, 153.71, 139.54, 138.35, 137.89, 132.42, 129.30, 127.51, 126.74, 125.76, 124.13, 115.31, 41.95, 39.36, 14.06. HRMS (ESI): *m/z* calcd for $C_{20}H_{21}FN_2O_2S_2$ 404.52, found $(M + H)^+$ 405.11.

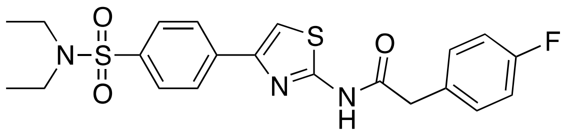

***N*-(4-(4-(*N*,*N*-diethylsulfamoyl)phenyl)thiazol-2-yl)-2-(4-fluorophenyl)acetamide (72).** $^1$H NMR (400 MHz, $CDCl_3$) δ, ppm: 8.02 (d, 2H), 7.86 (d, 2H), 7.49 (s, 1H), 7.32 (dd, 2H), 7.05 (t, 2H), 4.36 (s, 2H), 3.26 (q, 4H), 1.13 (t, 6H). $^{13}$C NMR (400 MHz, $CDCl_3$) δ, ppm: 170.82, 163.30, 160.86, 153.57, 139.49, 138.02, 133.24, 130.67, 127.52, 126.73, 115.82, 115.61, 115.19, 41.95, 38.94, 14.07. HRMS (ESI): *m/z* calcd for $C_{20}H_{21}FN_2O_2S_2$ 404.52, found $(M + H)^+$ 405.11.

### TABLE 2 COMPOUNDS

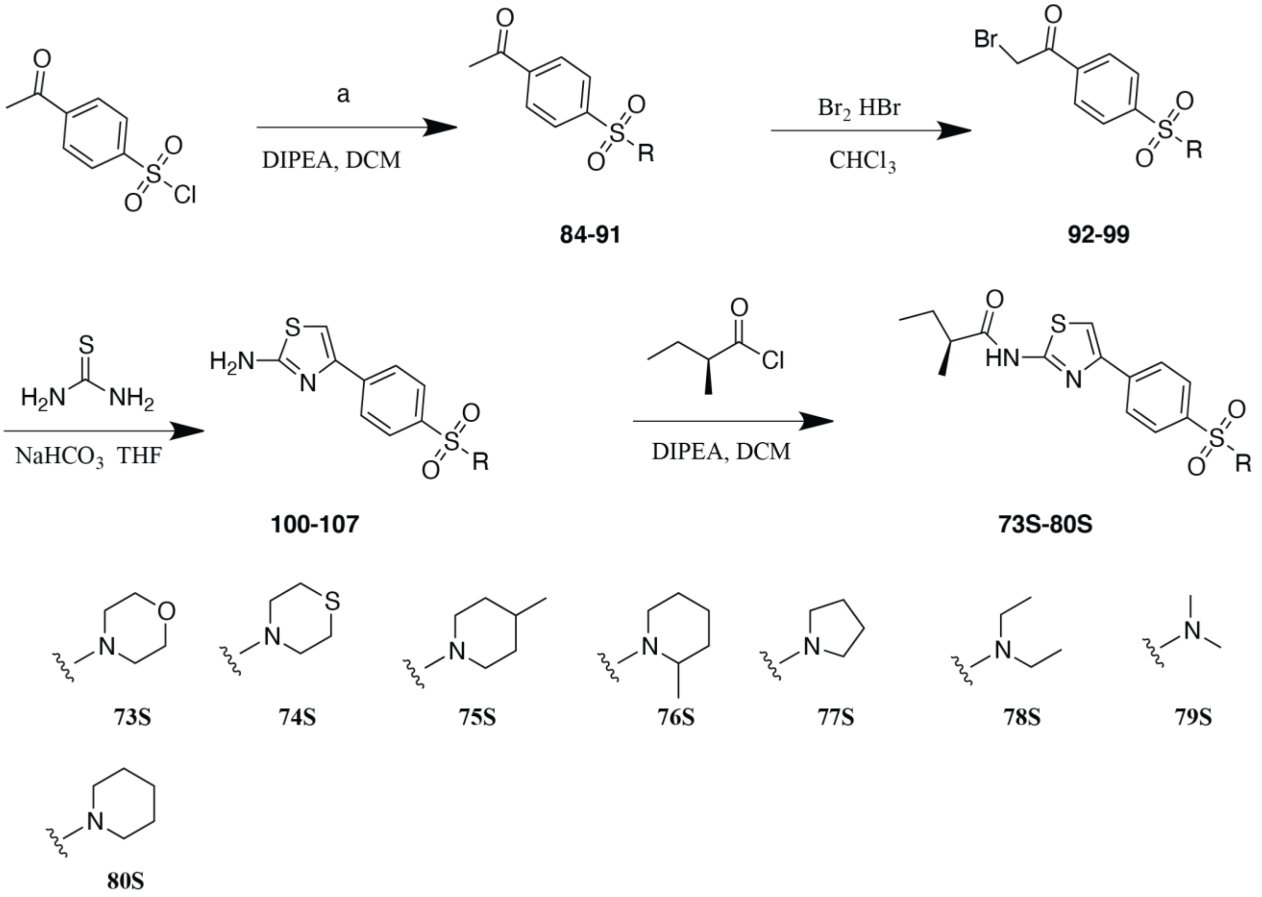

**Scheme SI 2: Chemical Synthesis of Intermediates for Stereopure S- R- aminiothiazole FASN-TE inhibitors**. (a) morpholine **(84)**, thiomorpholine **(85)**, 4-pipecoline **(86)**, 2-pipecoline **(87)**, pyrrolidine **(88)**, diisopropylamine **(89),** dimethylamine **(90),** piperidine **(91)**.

**Synthesis of 73S**

**1-(4-(morpholinosulfonyl)phenyl)ethan-1-one (84)** To a solution of 4-acetylbenzenesulfonyl chloride (3 g, 13.76 mmol) in DCM (30 mL) were added morpholine (1317 mg, 15.14 mmol) and *N*,*N*-Diisopropylethylamine (3557.1 mg, 27.52 mmol). The mixture was stirred at room temperature for 2 hours. The reaction mixture was quenched with water (30 mL), and then extracted with DCM (15 mL × 3). The combined organic layers were washed with brine (20 mL), and dried over $Na_2SO_4$. This was filtered and the filtrate was concentrated to provide the crude product which was purified by flash column chromatography (silica gel, eluting with 10% to 15% EA/PE) to afford 1-(4-(morpholinosulfonyl)phenyl)ethan-1-one (**84**) (3.38 g, 91.3%) as a white solid. HPLC/UV purity: 95%; LC-MS (ESI): 270.0 $(M + H)^+$.

**2-bromo-1-(4-(morpholinosulfonyl)phenyl)ethan-1-one (92)** To a solution of bromine (2.01 g, 12.56 mmol) in chloroform (65 mL) were added 1-(4-(morpholinosulfonyl)phenyl)ethan-1-one (3.38 g, 12.56 mmol) and hydrobromic acid (45% water) (1.5 mL) at 0°C. The mixture was stirred at room temperature for 3 hours. The reaction mixture was quenched with water (60 mL), and then extracted with DCM (30 mL × 3). The combined organic layers were washed with brine (30 mL × 2) and then dried over $Na_2SO_4$. This was filtered and the filtrate was concentrated to give the crude product and purified by flash column chromatography (silica gel, eluting with 10% to 15% EA/PE) to afford 2-bromo-1-(4-(morpholinosulfonyl)phenyl)ethan-1-one (**92**) (3.23 g, 74.1%) as a light yellow solid. HPLC/UV purity: 95%; LC-MS (ESI): 348.2 $(M + H)^+$. $^1H$ NMR (400 MHz, DMSO-$d_6$) δ 8.24 (d, $J$ = 2.0, 8.8 Hz, 2H), 7.91 (d, $J$ = 8.8 Hz, 2H), 5.04 (s, 2H), 3.64 (t, $J$ = 4.4 Hz, 4H), 1.92 (t, $J$ = 4.8 Hz, 4H).

**4-(4-(morpholinosulfonyl)phenyl)thiazol-2-amine (100)** To a solution of 2-bromo-1-(4-(morpholinosulfonyl)phenyl)ethan-1-one (1.63 g, 4.7 mmol) in THF (16 mL) were added thiourea (624.7 mg, 8.22 mmol) and sodium bicarbonate (848.3 mg, 10.1 mmol). The mixture was stirred at room temperature for 16 hours. The reaction mixture was quenched with water (40 mL), and then extracted with EA (20 mL × 3). The combined organic layers were washed with brine (30 mL) followed by a drying over $Na_2SO_4$. Subsequently the sample was filtered and the filtrate was concentrated to yield the crude product which was purified by flash column chromatography (silica gel, eluting with 30% to 40% EA/PE) to afford 4-(4-(morpholinosulfonyl)phenyl)thiazol-2-amine (**100**) (1.12 g, 73.4%) as a white solid. HPLC/UV purity: 95%; LC-MS (ESI): 326.1 $(M + H)^+$.$^1H$ NMR (400 MHz, DMSO-$d_6$) δ 8.05 (d, $J$ = 8.4 Hz, 2H), 7.72 (d, $J$ = 8.4 Hz), 7.31 (s, 1H), 7.12 (s, 2H), 3.63 (t , $J$ = 4.4 Hz, 4H), 2.88 (t, $J$ = 4.4 hz, 4H).

**Synthesis of 74S**

**1-(4-(thiomorpholinosulfonyl)phenyl)ethan-1-one (85).** To a solution of 4-acetylbenzenesulfonyl chloride (200 mg, 0.92 mmol) in DCM (5 mL) were added thiomorpholine (103.9 mg, 1.01 mmol) and *N*,*N*-Diisopropylethylamine (237.1 mg, 1.83 mmol). The mixture was stirred at room temperature for 2 h. The reaction mixture was quenched with water (15 mL), and then extracted with DCM (10 mL × 3). The combined organic layers were washed with brine (20 mL) and dried over $Na_2SO_4$. This solution was then filtered and then subsequently concentrated to give the crude product which was purified by flash column chromatography (silica gel, eluting with 15% to 20% EA/PE) to afford the reagent 1-(4-(thiomorpholinosulfonyl)phenyl)ethan-1-one (**85**) (215 mg, 81.9%) as a white solid. HPLC/UV purity: 95%; LC-MS (ESI): 286.1 $(M + H)^+$.

**2-bromo-1-(4-(thiomorpholinosulfonyl)phenyl)ethan-1-one (93).** To a solution of bromine (120.7 mg, 0.75 mmol) in chloroform (5 mL) were added 1-(4-(thiomorpholinosulfonyl)phenyl)ethan-1-one (215 mg, 0.75 mmol) and hydrobromic acid (45% water) (0.1 mL) at 0°C. The mixture was stirred at room temperature for 3 h. The reaction mixture was quenched with water (20 mL), and then extracted with DCM (10 mL × 3). The combined organic layers were washed with brine (15 mL × 2). The organic layer was then dried over $Na_2SO_4$. The sample was filtered and the filtrate was concentrated to give the crude product which was purified by flash column chromatography (silica gel, eluting with 10% to 15% EA/PE) to afford 2-bromo-1-(4-(thiomorpholinosulfonyl)phenyl)ethan-1-one (**93**) (165 mg, 60.2%) as a white solid. HPLC/UV purity: 95%; LC-MS (ESI): 364.2 $(M + H)^+$.

**4-(4-(thiomorpholinosulfonyl)phenyl)thiazol-2-amine (101).** To a solution of 2-bromo-1-(4-(thiomorpholinosulfonyl)phenyl)ethan-1-one (165 mg, 0.45 mmol) in THF (5 mL) were added thiourea (60.5 mg, 0.80 mmol) and sodium bicarbonate (82.1 mg, 0.98 mmol). The mixture was stirred at room temperature for 16 h. The reaction mixture was quenched with water (15 mL), and then extracted into EtOAc (10 mL × 3). The combined organic layers were washed with brine (15 mL) and the solution was dried over $Na_2SO_4$. This was filtered and concentrated to give the crude product which was purified by flash column chromatography (silica gel, eluting with 35% to 45% EA/PE) to afford 4-(4-(thiomorpholinosulfonyl)phenyl)thiazol-2-amine (**101**) (125 mg, 80.6%) as a white solid. HPLC/UV purity: 95%; LC-MS (ESI): 342.1 $(M + H)^+$.

**Synthesis of 75S**

**1-(4-((4-methylpiperidin-1-yl)sulfonyl)phenyl)ethan-1-one (86).** To a solution of 4-acetylbenzenesulfonyl chloride (400 mg, 1.83 mmol) in DCM (5 mL) were added 4-pipecoline (199.8 mg, 2.02 mmol) and *N*,*N*-Diisopropylethylamine (474.3 g, 3.67 mmol). The mixture was stirred at room temperature for 2 h. The reaction mixture was quenched with water (20 mL), and then extracted with DCM (10 mL × 3). The combined organic layers were washed with brine (10 mL), dried over $Na_2SO_4$ and then filtered. The filtrate solution was concentrated to give the crude product and this was followed by purification by flash column chromatography (silica gel, eluting with 10% to 15% EA/PE) to afford 1-(4-((4-methylpiperidin-1-yl)sulfonyl)phenyl)ethan-1-one (**86**) (377 mg, 73.1%) as a white solid. HPLC/UV purity: 95%; LC-MS (ESI): 282.1 $(M + H)^+$.

**2-bromo-1-(4-((4-methylpiperidin-1-yl)sulfonyl)phenyl)ethan-1-one (94)** To a solution of bromine (214.7 mg, 1.34 mmol) in chloroform (9.5 mL) were added 1-(4-(pyrrolidin-1-ylsulfonyl)phenyl)ethan-1-one (377 mg, 1.34 mmol) and Hydrobromic acid (45% water) (0.3 mL) at 0°C. The mixture was stirred at room temperature for 3 h. The reaction mixture was quenched with water (15 mL), and then extracted with DCM (10 mL × 3). The combined organic layers were washed with brine (15 mL × 2), dried over $Na_2SO_4$, and then filtered. The filtrate was concentrated to give the crude product and the sample was purified by flash column chromatography (silica gel, eluting with 10% to 15% EA/PE) to afford 2-bromo-1-(4-((4-methylpiperidin-1-yl)sulfonyl)phenyl)ethan-1-one (**94**) (355 mg, 73.7%) as a white solid. HPLC/UV purity: 95%; LC-MS (ESI): 360.0 $(M + H)^+$.

**4-(4-((4-methylpiperidin-1-yl)sulfonyl)phenyl)thiazol-2-amine (102).** To a solution of 2-bromo-1-(4-((4-methylpiperidin-1-yl)sulfonyl)phenyl)ethan-1-one (355 mg, 0.99 mmol) in THF (8 mL) were added thiourea (131.5 mg, 1.73 mmol) and sodium bicarbonate (178.6 mg, 2.12 mmol). The mixture was stirred at room temperature for 16 h. The reaction mixture was quenched with water (20 mL), and then extracted with EA (10 mL × 3). The combined organic layers were washed with brine (15 mL) followed by addition of $Na_2SO_4$. The solution was filtered, and the filtrate was concentrated to give the crude product. Final purification was achieved by flash column chromatography (silica gel, eluting with 30% to 40% EA/PE) to afford 4-(4-(pyrrolidin-1-ylsulfonyl)phenyl)thiazol-2-amine (**102**) (289 mg, 86.7%) as a white solid. HPLC/UV purity: 95%; LC-MS (ESI): 338.1 $(M + H)^+$.

**Synthesis of 76S**

**1-(4-((2-methylpiperidin-1-yl)sulfonyl)phenyl)ethan-1-one (87).** To a solution of 4-acetylbenzenesulfonyl chloride (200 mg, 0.92 mmol) in DCM (5 mL) were added 2-methylpiperidine (99.9 mg, 1.01 mmol) and *N*,*N*-Diisopropylethylamine (237.1 mg, 1.83 mmol). The mixture was stirred at room temperature for 2 h. The reaction mixture was quenched with water (15 mL), and then extracted with DCM (10 mL × 3). The combined organic layers were washed with brine (20 mL) and then dried over $Na_2SO_4$. The solids removed by filtration and the filtrate was concentrated to obtain the crude product which was then purified by flash column chromatography (silica gel, eluting with 10% to 15% EA/PE) to afford 1-(4-((2-methylpiperidin-1-yl)sulfonyl)phenyl)ethan-1-one (**87**) (220 mg, 85.3%) as a white solid. HPLC/UV purity: 95%; LC-MS (ESI): 282.1 $(M + H)^+$.

**2-bromo-1-(4-((2-methylpiperidin-1-yl)sulfonyl)phenyl)ethan-1-one (95).** To a solution of bromine (125.3 mg, 0.78 mmol) in chloroform (5 mL) were added 1-(4-((2-methylpiperidin-1-yl)sulfonyl)phenyl)ethan-1-one (220 mg, 0.78 mmoL) and Hydrobromic acid (45% water) (0.1 mL) at 0°C. The mixture was stirred at room temperature for 3 h. The reaction mixture was quenched with water (20 mL), and then extracted with DCM (10 mL × 3). The combine organic layers were washed with brine (15 mL × 2) and subsequently was dried over $Na_2SO_4$. This solution was filtered and the filtrate was concentrated to obtain the crude product which was purified by flash column chromatography (silica gel, eluting with 10% to 15% EA/PE) to afford 2-bromo-1-(4-((2-methylpiperidin-1-yl)sulfonyl)phenyl)ethan-1-one (**95**) (166 mg, 59.1%) as a white solid. HPLC/UV purity: 95%; LC-MS (ESI): 360.1 $(M + H)^+$.

**4-(4-((2-methylpiperidin-1-yl)sulfonyl)phenyl)thiazol-2-amine (103).** To a solution of 2-bromo-1-(4-((2-methylpiperidin-1-yl)sulfonyl)phenyl)ethan-1-one (166 mg, 0.46 mmol) in THF (5 mL) were added thiourea (61.5 mg, 0.81 mmol) and sodium bicarbonate (83.5 mg, 0.99 mmol). The mixture was stirred at room temperature for 16 h. The reaction mixture was quenched with water (15 mL), and then extracted with EA (10 mL × 3). The combined organic layers were washed with brine (15 mL) followed by treatment with $Na_2SO_4$ to dry the solution. This crude product was filtered, and the filtrate was concentrated to obtain the impure product which was purified by flash column chromatography (silica gel, eluting with 30% to 40% EA/PE) to afford 4-(4-((2-methylpiperidin-1-yl)sulfonyl)phenyl)thiazol-2-amine (**103**) (130 mg, 83.4%) as a white solid. HPLC/UV purity: 95%; LC-MS (ESI): 338.1 $(M + H)^+$.

**Synthesis of 77S**

**1-(4-(pyrrolidin-1-ylsulfonyl)phenyl)ethan-1-one (88).** To a solution of 4-acetylbenzenesulfonyl chloride (1.5 g, 6.88 mmol) in DCM (10 mL) were added pyrrolidine (537.4 mg, 7.57 mmol) and *N*,*N*-Diisopropylethylamine (1.78 g, 13.76 mmol). The mixture was stirred at room temperature for 2 h. The reaction mixture was quenched with water (20 mL), and then extracted with DCM (10 mL × 3). The combined organic layers were washed with brine (10 mL) and subsequently dried over $Na_2SO_4$. This solution was then filtered, and the filtrate was concentrated to give the crude product. Final purification was by flash column chromatography (silica gel, eluting with 10% to 15% EA/PE) to afford 1-(4-(pyrrolidin-1-ylsulfonyl)phenyl)ethan-1-one (**88**) (1.38 g, 79%) as a white solid. HPLC/UV purity: 95%; LC-MS (ESI): 254.1 $(M + H)^+$.

**2-bromo-1-(4-(pyrrolidin-1-ylsulfonyl)phenyl)ethan-1-one (96).** To a solution of bromine (869.3 mg, 5.43 mmol) in chloroform (35 mL) were added 1-(4-(pyrrolidin-1-ylsulfonyl)phenyl)ethan-1-one (1.38 g, 5.43 mmol) and hydrobromic acid (45% water) (0.5 mL) at 0°C. The mixture was stirred at room temperature for 3 h. The reaction mixture was quenched with water (30 mL), and then extracted with DCM (15 mL × 3). The combined organic layers were washed with brine (15 mL × 2) followed by a drying step over $Na_2SO_4$. The solution was filtered, and the filtrate was concentrated to give the crude product which was then purified by flash column chromatography (silica gel, eluting with 10% to 15% EA/PE) to afford 2-bromo-1-(4-(pyrrolidin-1-ylsulfonyl)phenyl)ethan-1-one (**96**) (1.25 g, 69.5%) as a light-yellow solid. HPLC/UV purity: 95%; LC-MS (ESI): 332.1 $(M + H)^+$.

**4-(4-(pyrrolidin-1-ylsulfonyl)phenyl)thiazol-2-amine (104).** To a solution of 2-bromo-1-(4-(pyrrolidin-1-ylsulfonyl)phenyl)ethan-1-one (1.25 g, 3.77 mmol) in THF (25 mL) were added thiourea (502 mg, 6.61 mmol) and sodium bicarbonate (682 mg, 8.12 mmol). The mixture was stirred at room temperature for 16 h. The reaction mixture was quenched with water (30 mL), and then extracted with EA (15 mL × 3). The combined organic layers were washed with brine (15 mL) and then dried over $Na_2SO_4$. This was then filtered and the filtrate was concentrated to give the crude product. The sample was purified by flash column chromatography (silica gel, eluting with 30% to 40% EA/PE) to afford 4-(4-(pyrrolidin-1-ylsulfonyl)phenyl)thiazol-2-amine (**104**) (954 mg, 81.7%) as a white solid. HPLC/UV purity: 95%; LC-MS (ESI): 310.1 $(M + H)^+$.

**Synthesis of 78S/78R**

**4-acetyl-*N*,*N*-diethylbenzenesulfonamide (89)** To a solution of 4-acetylbenzenesulfonyl chloride (1.5 g, 6.88 mmol) in DCM (10 mL) were added diethylamine (552.5 mg, 7.57 mmol) and *N*,*N*-Diisopropylethylamine (1.78 g, 13.76 mmol). The mixture was stirred at room temperature for 2 hours. The reaction mixture was quenched with water (20 mL), and then extracted with DCM (10 mL × 3). The combined organic layers were washed with brine (10 mL) followed by a drying step over $Na_2SO_4$. Filtration of this reaction was followed by a concentration step to give the crude product. The parent compound was purified by flash column chromatography (silica gel, eluting with 10% to 15% EA/PE) to afford 4-acetyl-*N*,*N*-diethylbenzenesulfonamide (**89**) (1.52 g, 79%) as a white solid. HPLC/UV purity: 95%; LC-MS (ESI): 256.1 $(M + H)^+$.

**4-(2-bromoacetyl)-*N*,*N*-diethylbenzenesulfonamide (97)** To a solution of bromine (953.7 mg, 5.43 mmol) in chloroform (35 mL) were added 4-acetyl-*N*,*N*-diethylbenzenesulfonamide (1.52 g, 5.43 mmol) and hydrobromic acid (45% water) (0.5 mL) at 0ºC. The mixture was stirred at room temperature for 3 hours. The reaction mixture was quenched with water (30 mL), and then extracted with DCM (15 mL × 3). The combined organic layers were washed with brine (15 mL × 2) and dried over $Na_2SO_4$. The crude product was filtered, concentrated and purified by flash column chromatography (silica gel, eluting with 10% to 15% EA/PE) to afford 4-(2-bromoacetyl)-*N*,*N*-diethylbenzenesulfonamide (**97**) (1.72 g, 86.6%) as a white solid. HPLC/UV purity: 95%; LC-MS (ESI): 334.1 $(M + H)^+$.

**4-(2-aminothiazol-4-yl)-*N*,*N*-diethylbenzenesulfonamide (105)** To a solution of 4-(2-bromoacetyl)-*N*,*N*-diethylbenzenesulfonamide (1.12 g, 3.36 mmol) in THF (12 mL) were added thiourea (447.3 mg, 5.89 mmol) and sodium bicarbonate (607.4 mg, 7.23 mmol). The mixture was stirred at room temperature for 16 hours. The reaction mixture was quenched with water (30 mL), and then extracted with EA (15 mL × 3). The combined organic layers were washed with brine (15 mL), dried over $Na_2SO_4$, filtered and then concentrated. Purification by flash column chromatography (silica gel, eluting with 30% to 40% EA/PE) afforded 4-(2-aminothiazol-4-yl)-*N*,*N*-diethylbenzenesulfonamide (**105**) (763 mg, 72.9%) as a white solid. HPLC/UV purity: 95%; LC-MS (ESI): 312.1 $(M + H)^+$.

**Synthesis of 79S**

**4-acetyl-*N,N*-dimethylbenzenesulfonamide (90).** To a solution of 4-acetylbenzenesulfonyl chloride (200 mg, 0.92 mmol) in DCM (5 mL) were added dimethylamine (0.69 mL, 1.37 mmol,2 mol/L in THF) and *N*,*N*-Diisopropylethylamine (237.1 mg, 1.83 mmol). The mixture was stirred at room temperature for 2 h. The reaction mixture was quenched with water (15 mL), and then extracted with DCM (10 mL × 3). The combined organic layers were washed with brine (20 mL) and dried over $Na_2SO_4$. The solution was filtered, and the filtrate was concentrated to obtain the crude product which was purified by flash column chromatography (silica gel, eluting with 10% to 15% EA/PE) to afford 4-acetyl-*N,N*-dimethylbenzenesulfonamide (**90**) (195 mg, 93.6%) as a white solid. HPLC/UV purity: 95%; LC-MS (ESI): 228.1 $(M + H)^+$.

**4-(2-bromoacetyl)-*N*,*N*-dimethylbenzenesulfonamide (98).** To a solution of bromine (137.4 mg, 0.86 mmol) in chloroform (5 mL) were added 4-acetyl-*N,N*-dimethylbenzenesulfonamide (195 mg, 0.86 mmol) and Hydrobromic acid (45% water) (0.1 mL) at 0°C. The mixture was stirred at room temperature for 3 h. The reaction mixture was quenched with water (20 mL), and then extracted with DCM (10 mL × 3). The combined organic layers were washed with brine (15 mL × 2) and were dried over $Na_2SO_4$. This solution was filtered, and the filtrate was concentrated to obtain the crude product which was purified by flash column chromatography (silica gel, eluting with 10% to 15% EA/PE) to afford 4-(2-bromoacetyl)-*N*,*N*-dimethylbenzenesulfonamide (**98**) (150 mg, 57.3%) as a white solid. HPLC/UV purity: 95%; LC-MS (ESI): 306.0 $(M + H)^+$.

**4-(2-aminothiazol-4-yl)-*N*,*N*-dimethylbenzenesulfonamide (106).** To a solution of 4-(2-bromoacetyl)-*N*,*N*-dimethylbenzenesulfonamide (150 mg, 0.49 mmol) in THF (5 mL) were added thiourea (65.4 mg, 0.86 mmol) and sodium bicarbonate (88.8 mg, 1.06 mmol). The mixture was stirred at room temperature for 16 h. The reaction mixture was quenched with water (15 mL), and then extracted with EA (10 mL × 3). The combined organic layers were washed with brine (15 mL) and were then dried over $Na_2SO_4$. The solution was filtered, and the filtrate was concentrated to obtain the crude product which was purified by flash column chromatography (silica gel, eluting with 30% to 40% EA/PE) to afford 4-(2-aminothiazol-4-yl)-*N*,*N*-dimethylbenzenesulfonamide (**106**) (115 mg, 82.3%) as a white solid. HPLC/UV purity: 95%; LC-MS (ESI): 284.1 $(M + H)^+$.

**Synthesis of 80S**

**1-(4-(piperidin-1-ylsulfonyl)phenyl)ethan-1-one (91).** To a solution of 4-acetylbenzenesulfonyl chloride (800 mg, 3.67 mmol) in DCM (20 mL) were added piperidine (343.1 mg, 4.04 mmol) and *N*,*N*-Diisopropylethylamine (948.5 mg, 7.34 mmol). The mixture was stirred at room temperature for 2 h. The reaction mixture was quenched with water (30 mL), and then extracted with DCM (15 mL × 3). The combined organic layers were washed with brine (30 mL), dried over $Na_2SO_4$. Filtered and the filtrate was concentrated to give the crude product which was purified by flash column chromatography (silica gel, eluting with 10% to 15% EA/PE) to afford 1-(4-(piperidin-1-ylsulfonyl)phenyl)ethan-1-one (**91**) (820 mg, 83.7%) as a white solid. HPLC/UV purity: 95%; LC-MS (ESI): 268.1 $(M + H)^+$.

**2-bromo-1-(4-(piperidin-1-ylsulfonyl)phenyl)ethan-1-one (99).** To a solution of bromine (491.4 mg, 3.07 mmol) in chloroform (20 mL) were added 1-(4-(piperidin-1-ylsulfonyl)phenyl)ethan-1-one (820 mg, 3.07 mmol) and Hydrobromic acid (45% water) (0.2 mL) at 0°C. The mixture was stirred at room temperature for 3 h. The reaction mixture was quenched with water (40 mL), and then extracted with DCM (20 mL × 3). The combined organic layers were washed with brine (20 mL × 2) and dried over $Na_2SO_4$. The sample was filtered, and the filtrate was concentrated to give the crude product which was purified by flash column chromatography (silica gel, eluting with 10% to 15% EA/PE) to afford 2-bromo-1-(4-(piperidin-1-ylsulfonyl)phenyl)ethan-1-one (**99**) (766 mg, 72.3%) as a white solid. HPLC/UV purity: 95%; LC-MS (ESI): 346.0 $(M + H)^+$.

**4-(4-(piperidin-1-ylsulfonyl)phenyl)thiazol-2-amine (107).** To a solution of 2-bromo-1-(4-(piperidin-1-ylsulfonyl)phenyl)ethan-1-one (766 mg, 2.22 mmol) in THF (8 mL) were added thiourea (295.3 mg, 3.88 mmol) and sodium bicarbonate (401 mg, 4.77 mmol). The mixture was stirred at room temperature for 16 h. The reaction mixture was quenched with water (30 mL), and then extracted with EA (15 mL × 3). The combined organic layers were washed with brine (15 mL) and then dried over $Na_2SO_4$. The solution was filtered, and the filtrate was concentrated to give the crude product which was purified by flash column chromatography (silica gel, eluting with 30% to 40% EA/PE) to afford 4-(4-(piperidin-1-ylsulfonyl)phenyl)thiazol-2-amine (**107**) (560 mg, 78.1%) as a white solid. HPLC/UV purity: 95%; LC-MS (ESI): 324.1 $(M + H)^+$. $^1$H NMR (400 MHz, DMSO-$d_6$) δ 8.02 (d, $J$ = 8.4 Hz, 2H), 7.70 (d, $J$ = 8.4 Hz, 2H), 7.28 (s, 1H), 7.18 (s,2H), 2.89 (t, $J$ = 5.2 Hz, 4H), 1.53 (t, $J$ = 4.8 Hz, 4H), 1.36 (d, $J$ = 4.8 Hz, 2H).

**HPLC purity of compounds used for in vivo studies**

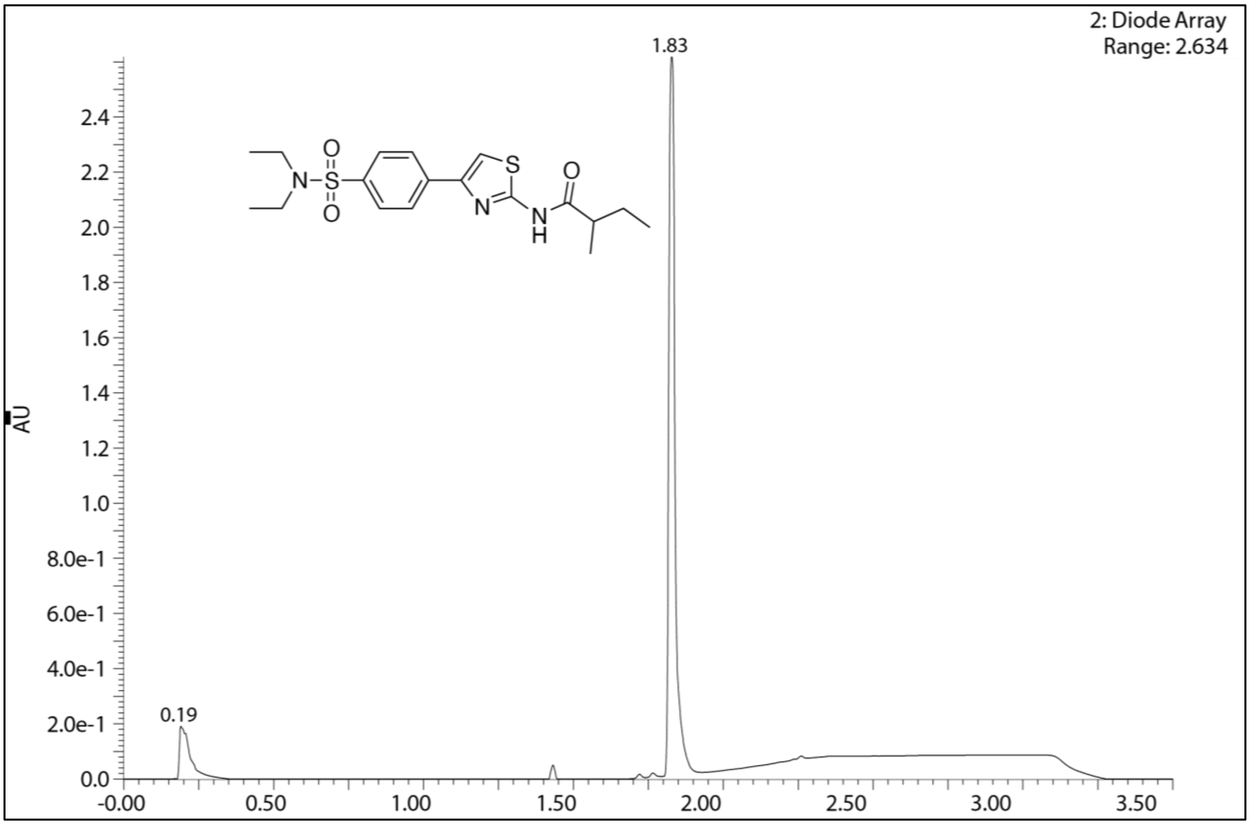

**HPLC chromatogram for (50)**

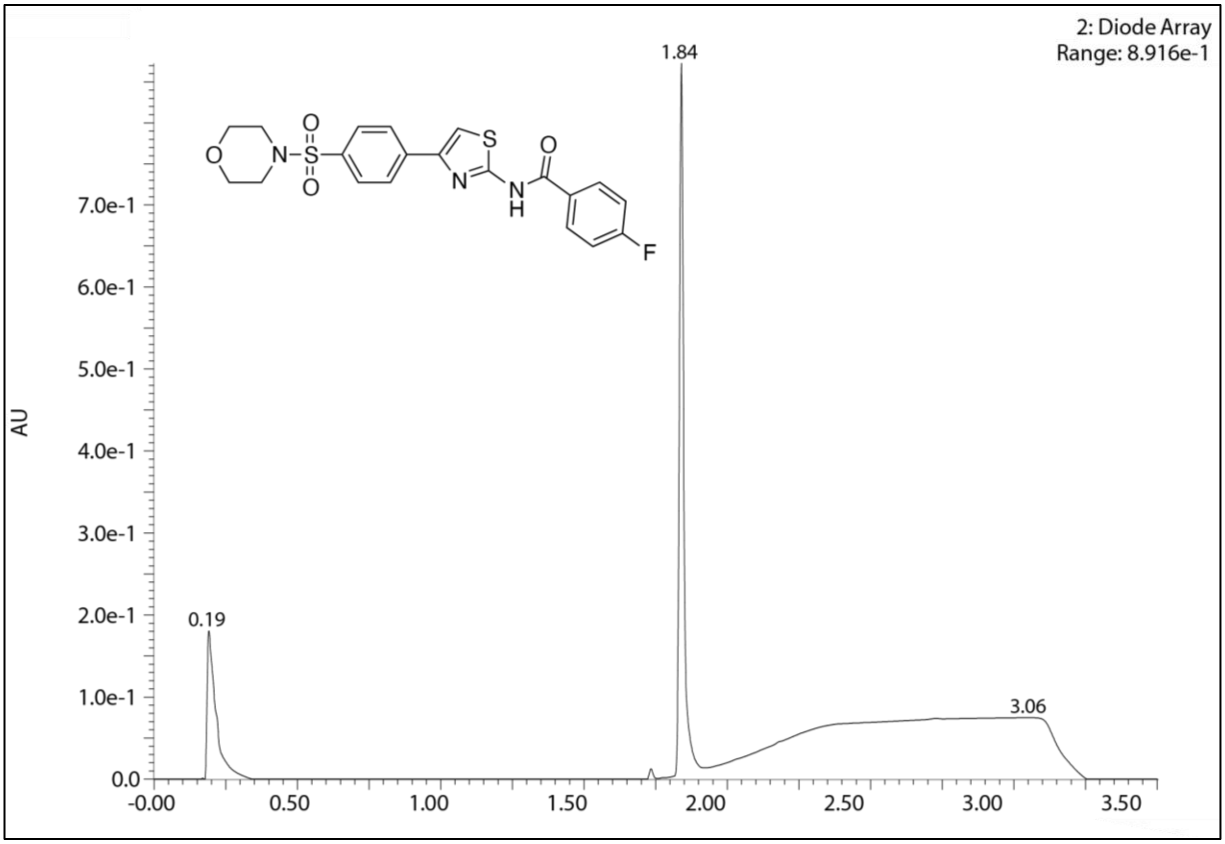

**HPLC chromatogram for (66)**

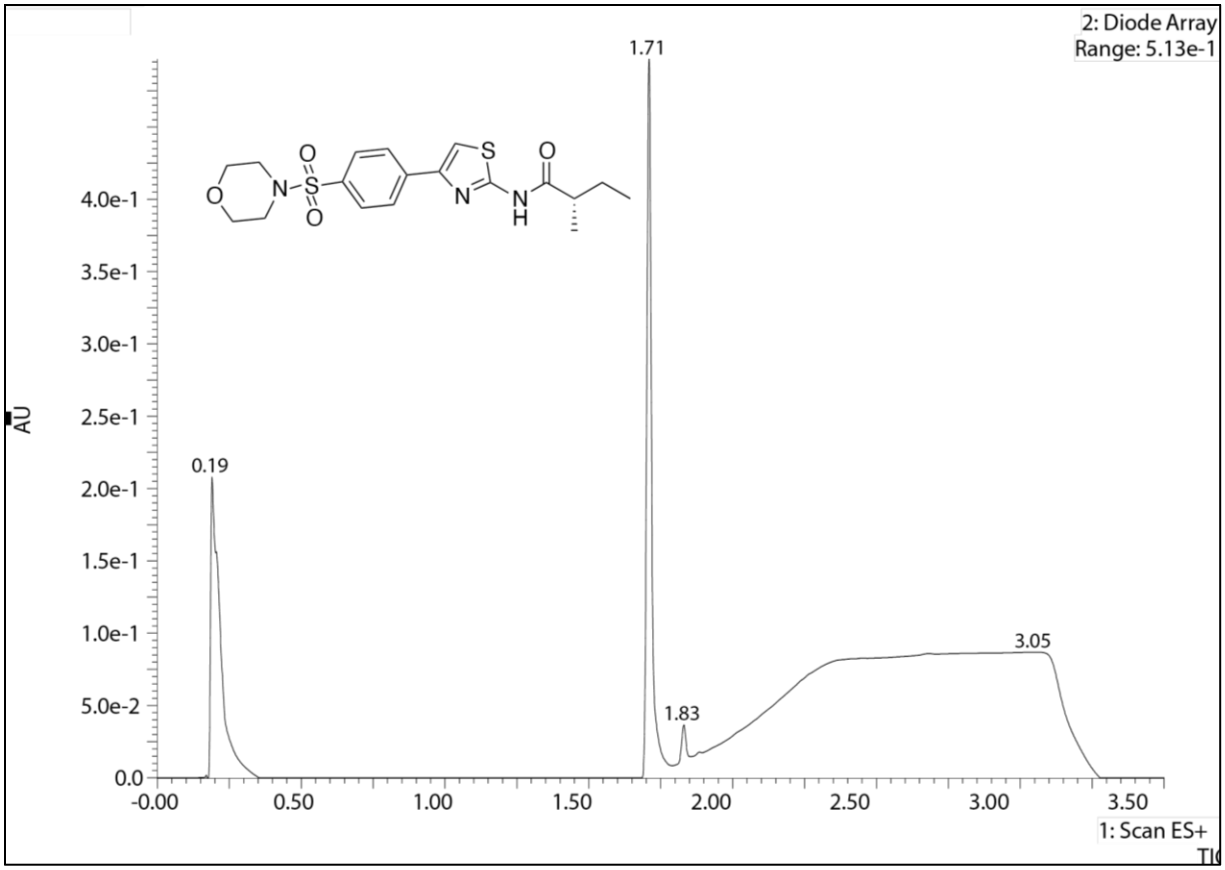

**HPLC chromatogram for (73S)**

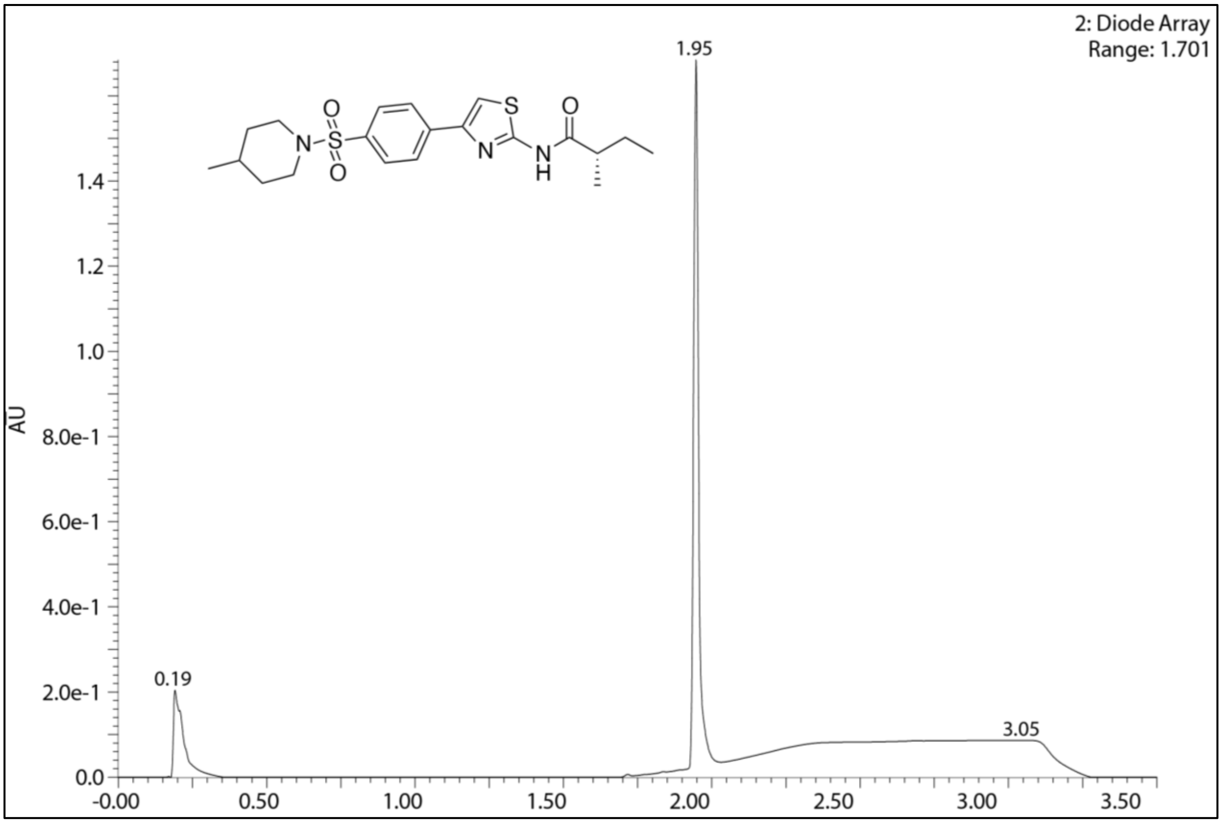

**HPLC chromatogram for (75S)**

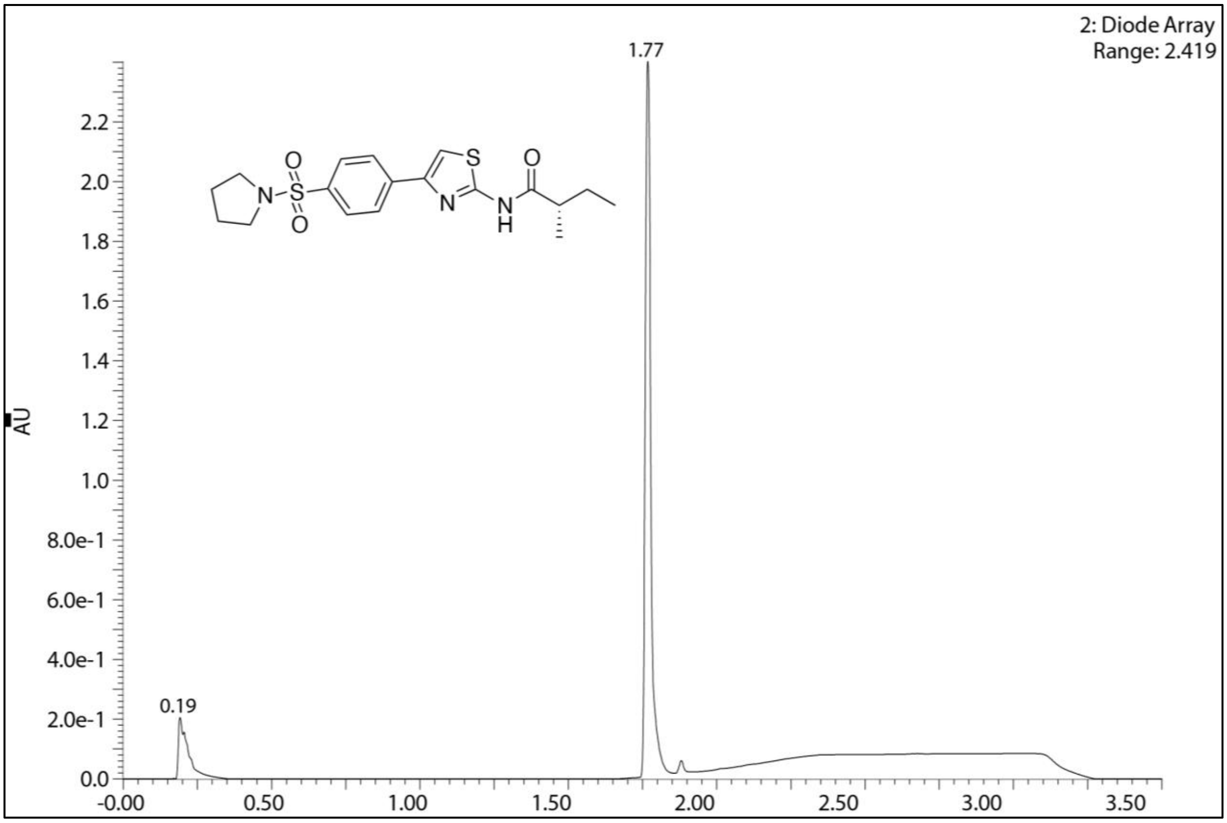

**HPLC chromatogram for (77S)**

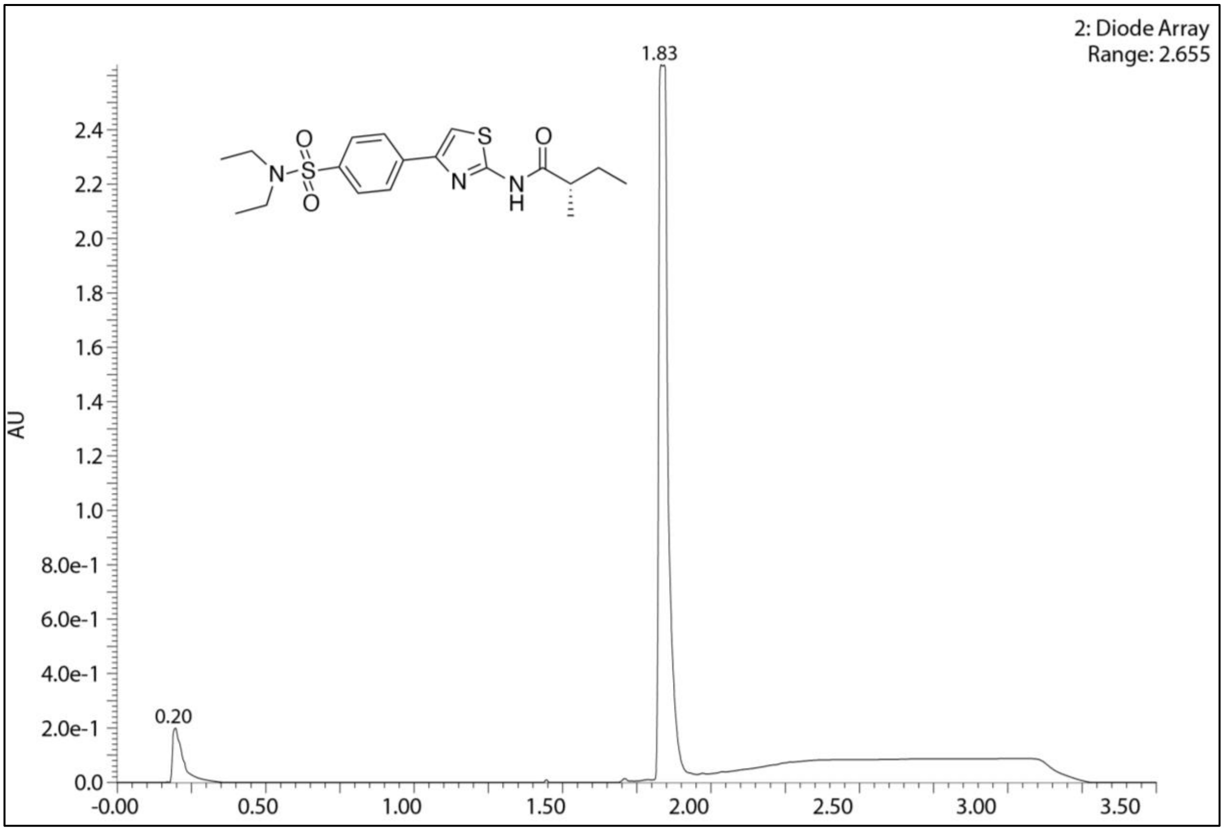

**HPLC chromatogram for (78S)**

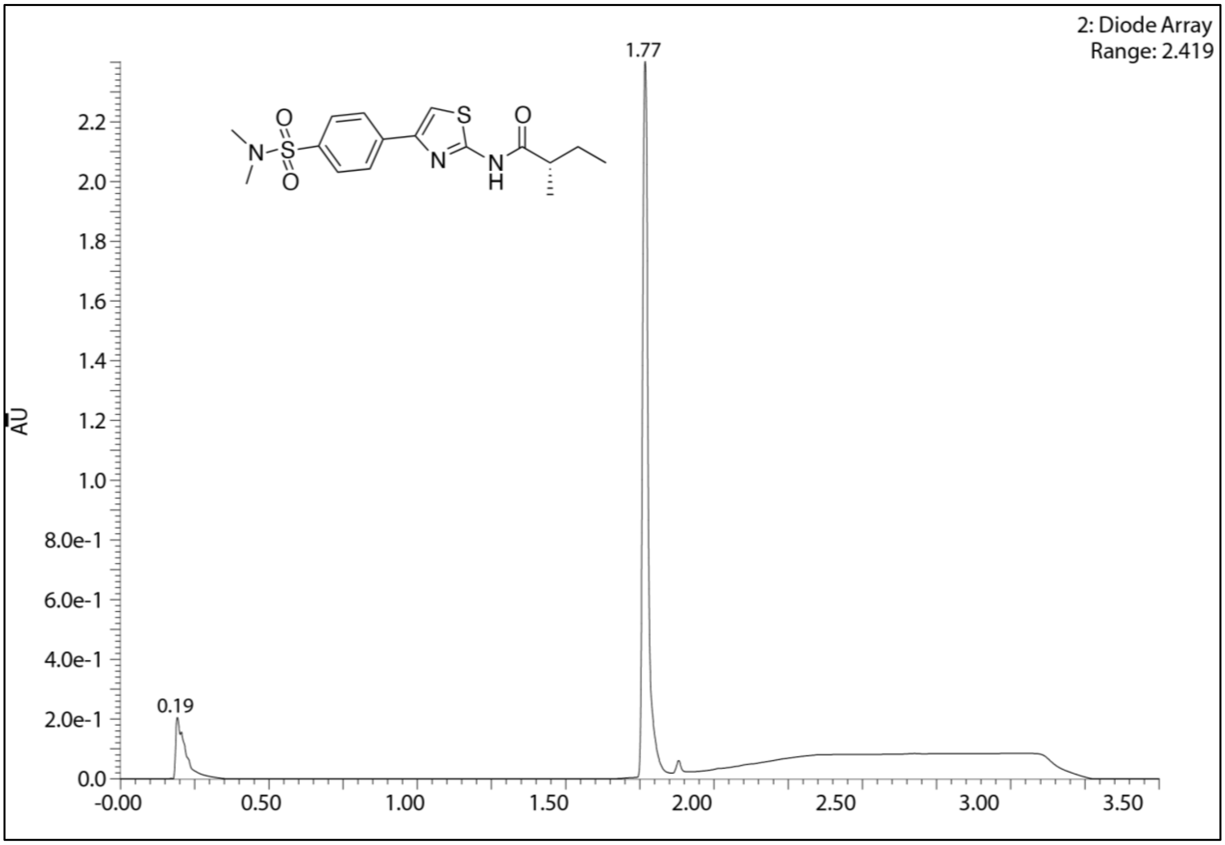

**HPLC chromatogram for (79S)**

**Chiral resolution HPLC chromatograms for 50, 78S, 78R**

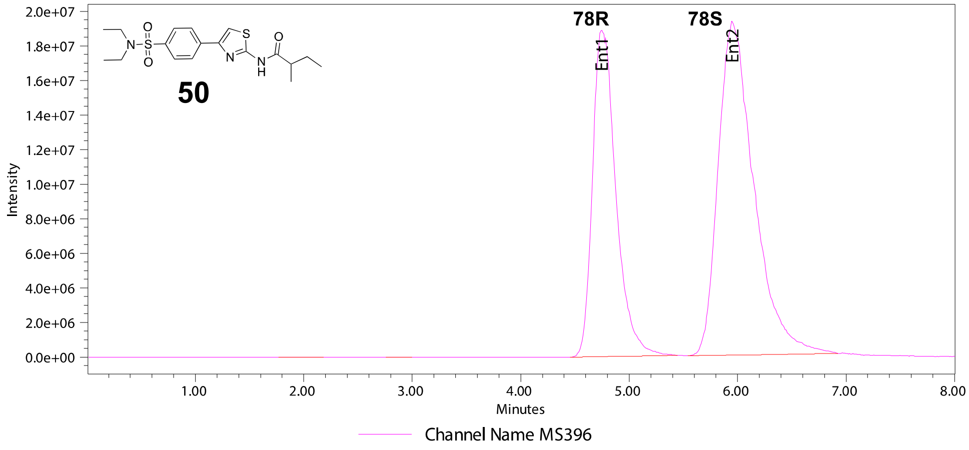

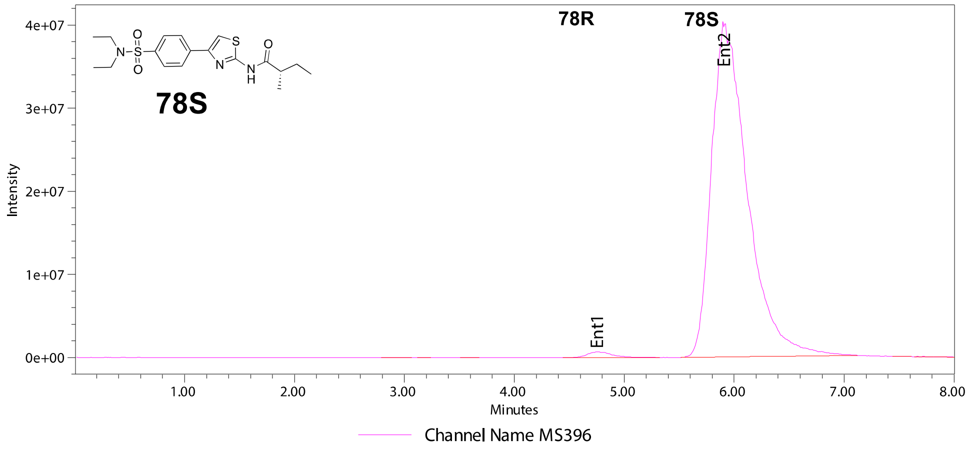

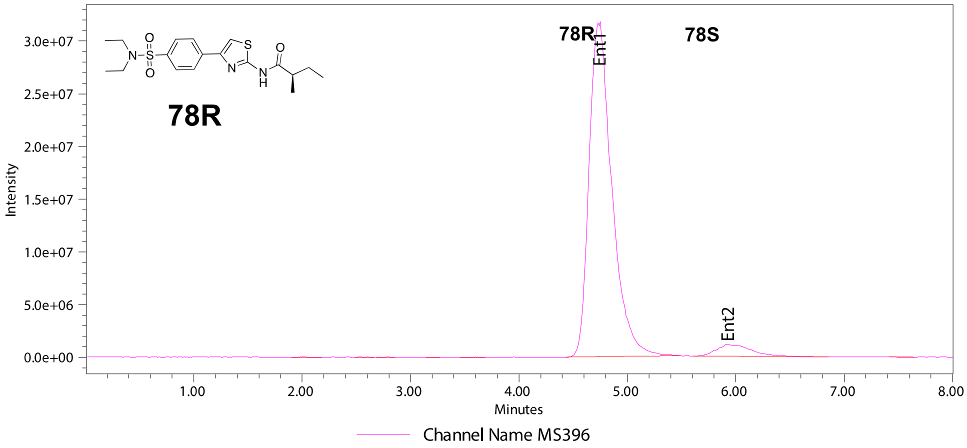

| | Ent / Retention Time | | | | | |
|---|---|---|---|---|---|---|
| Sample Name | Ent 1 / 4.75 | Ent 2 / 5.90 | EE | Ent 1 (μV*sec) | Ent 1 (μV*sec) | %EE |
| **50-** mixture | 38.99 | 61.01 | -22.02 | 282774953 | 442504434 | |
| **78S** | 1.14 | 98.86 | -97.72 | 10520698 | 912489691 | 97.72 |
| **78R** | 94.90 | 5.10 | 89.79 | 463618991 | 24938014 | 89.79 |
